# Immunosuppression, rather than inflammation, is a salient feature of sepsis in an Indian cohort

**DOI:** 10.1101/742924

**Authors:** Samanwoy Mukhopadhyay, Pravat K Thatoi, Bidyut K Das, Saroj K Mohapatra

## Abstract

Sepsis remains a lethal ailment with imprecise treatment and ill-understood biology. A clinical transcriptomic analysis of sepsis patients was performed for the first time in India and revealed large-scale change in blood gene expression in patients of severe sepsis and septic shock admitted to ICU. Three biological processes were quantified using scores derived from the corresponding transcriptional modules. Comparison of the module scores revealed that genes associated with immune response were more suppressed compared to the inflammation-associated genes. These findings will have great implication in the treatment and prognosis of severe sepsis/septic shock if it can be translated into a bedside tool.

## 2 Introduction

Sepsis is a condition with severe systemic inflammation accompanied by dysregualted host response to infection. It is one of the leading causes of hospital stay, death and economic burden in worldwide health-care [1–3]. Sepsis is generally considered a disease continuum from bacterial infection through systemic inflammation, progressing to severe sepsis with onset of organ failure, ultimately developing into the most lethal septic shock [1]. Pneumonia, urinary tract infection and intra-abdominal infection are the chief causes leading to sepsis [1]. Bacteria form the majority of etiologic microorganisms but the bacterial classes differ geographically. Gram-negative bacteria in India, such as, *Escherichia coli*, *Klebsiella species* and *Pseudomonas aeruginosa* [4], dominate in contrast to the gram-positives in the West. While there are many pathogens causing sepsis, human host response to the infection is very complex. A large body of literature has provided insights into the genome-scale biology of sepsis [5–8]. However, the clinical predictions from these discoveries remain to be tested in Indian patients. We set out to perform unbiased blood transcriptomics in patient with severe sepsis or septic shock (SS) from an Indian ICU. One of the major focus of our study was to find the genes and gene sets associated with poor outcome. Accordingly analysis was performed at multiple levels (genes, gene sets and networks) converging on key molecular processes (transcriptional modules) associated with outcome.

## 3 Results

The analysis plan for our study contained two arms (Figure 1)–firstly detection of transcriptomic changes associated with sepsis and secondly to identify pathways associated with survival.

**Figure 1.**
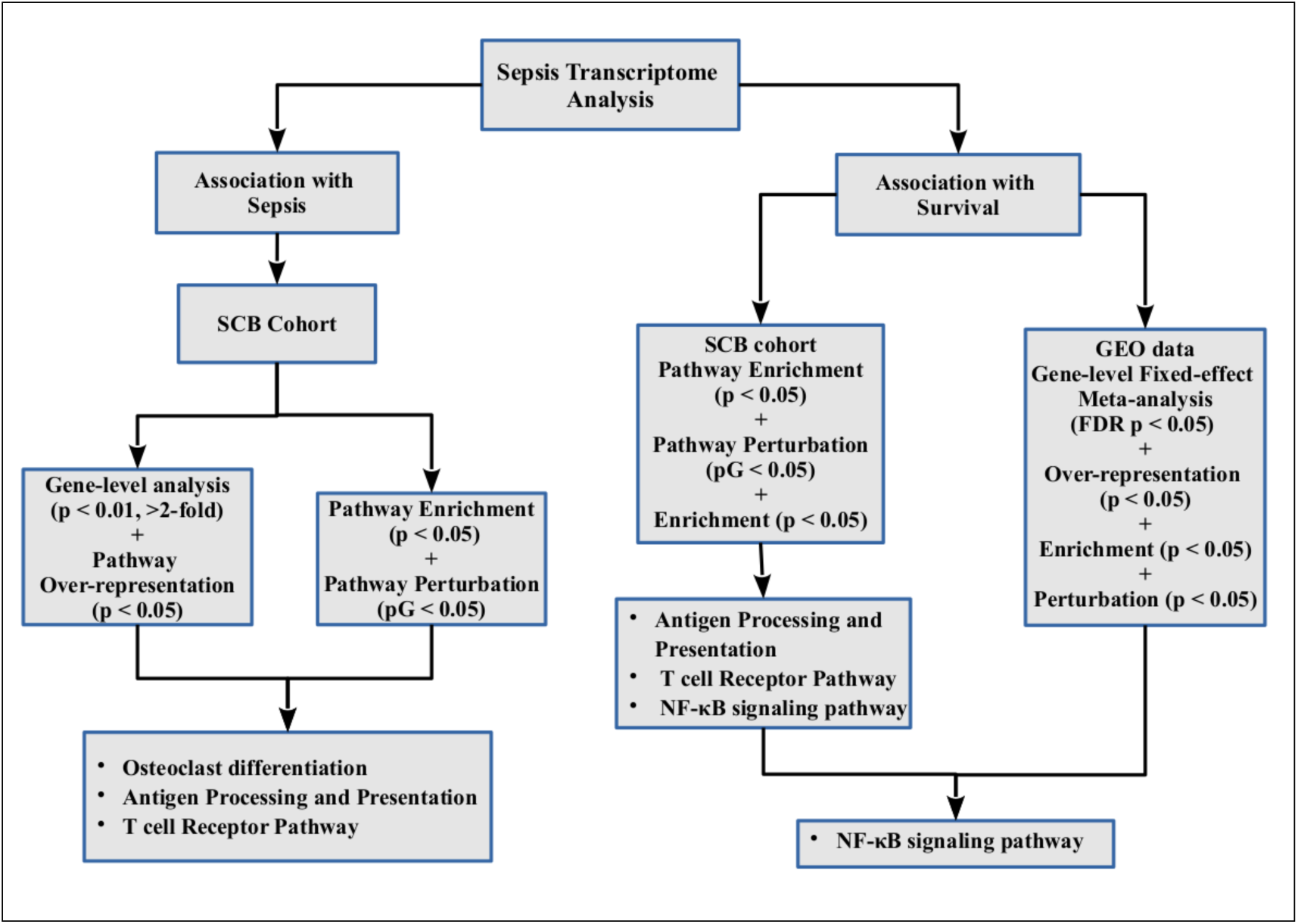
Analysis flow with results: The left arm describes the steps for case-control analysis and the right arm describes the analysis for association with survival.

### 3.1 Genome-level changes in gene expression

Differential gene expression analysis revealed 4221 genes (24% of the total number of 17513 genes assayed) to be significantly altered in sepsis patients compared to age- and gender-matched healthy controls (Figure 2). This established large scale change in gene expression in sepsis, impacting multiple pathways.

**Figure 2.**
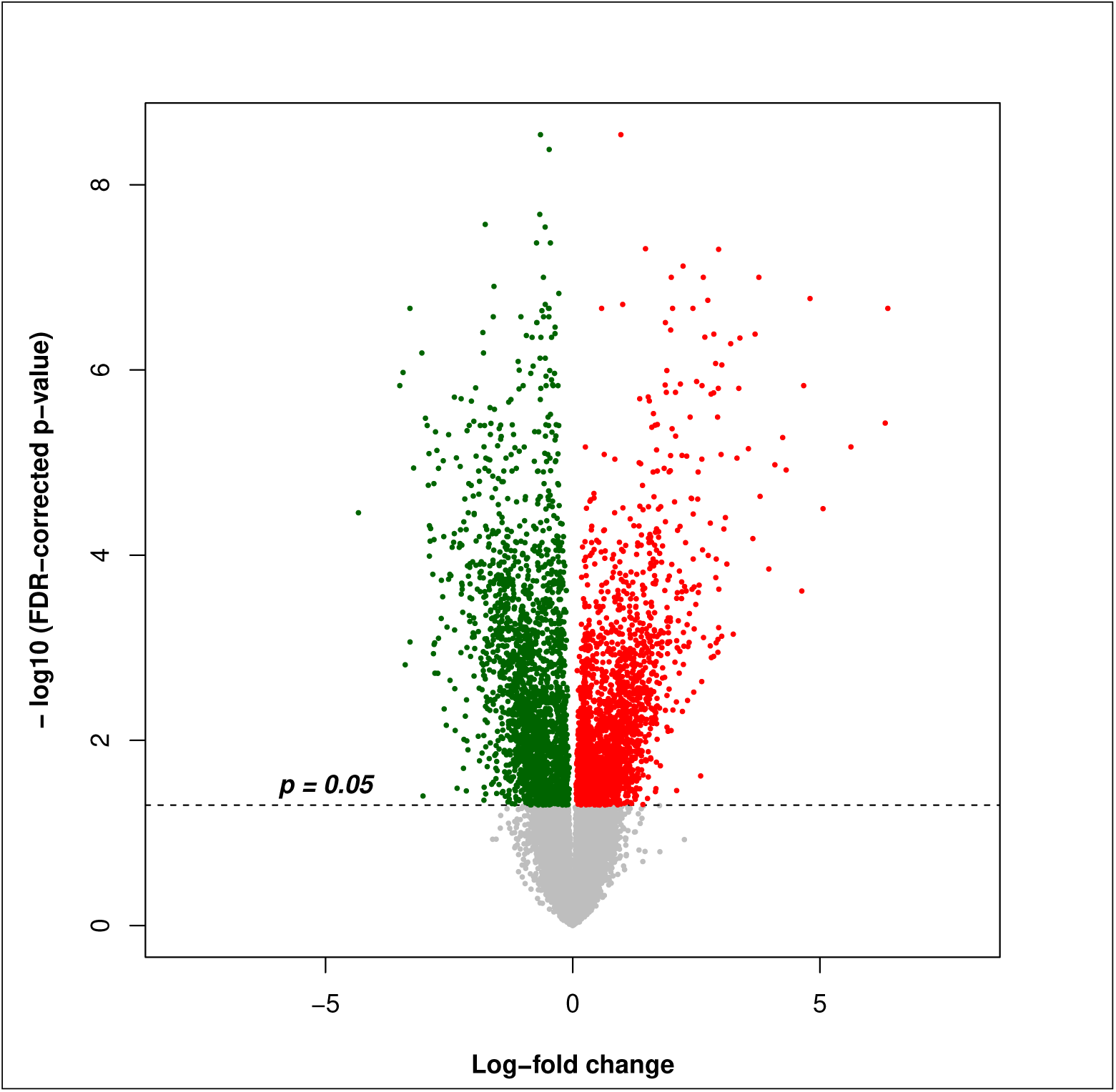
Volcano plot showing genome-wide change in gene expression in Sepsis: The Y-axis is in the negative logarithmic scale, showing more significant genes (smaller p-values) at the top. Significant genes (FDR p *<* 0.05) are shown above the dashed line. Green and red dots represent down- and up-regulated genes respectively. Approximately 24% of the genome (4221 out of 17513 genes) was observed to be differentially expressed.

### 3.2 Temporal change in gene expression

Temporal analysis of differentially expressed (DE) genes (FDR p *<* 0.05, 2 fold or greater change) revealed a non-random trend toward the baseline with time (Figure 3). Additionally, we observed a difference between survivors and non-survivors (Figure S4) suggesting delayed recovery of the transcriptome in non-survivors compared to survivors. This is consistent with the trends observed in critically ill subjects [9].

**Figure 3.**
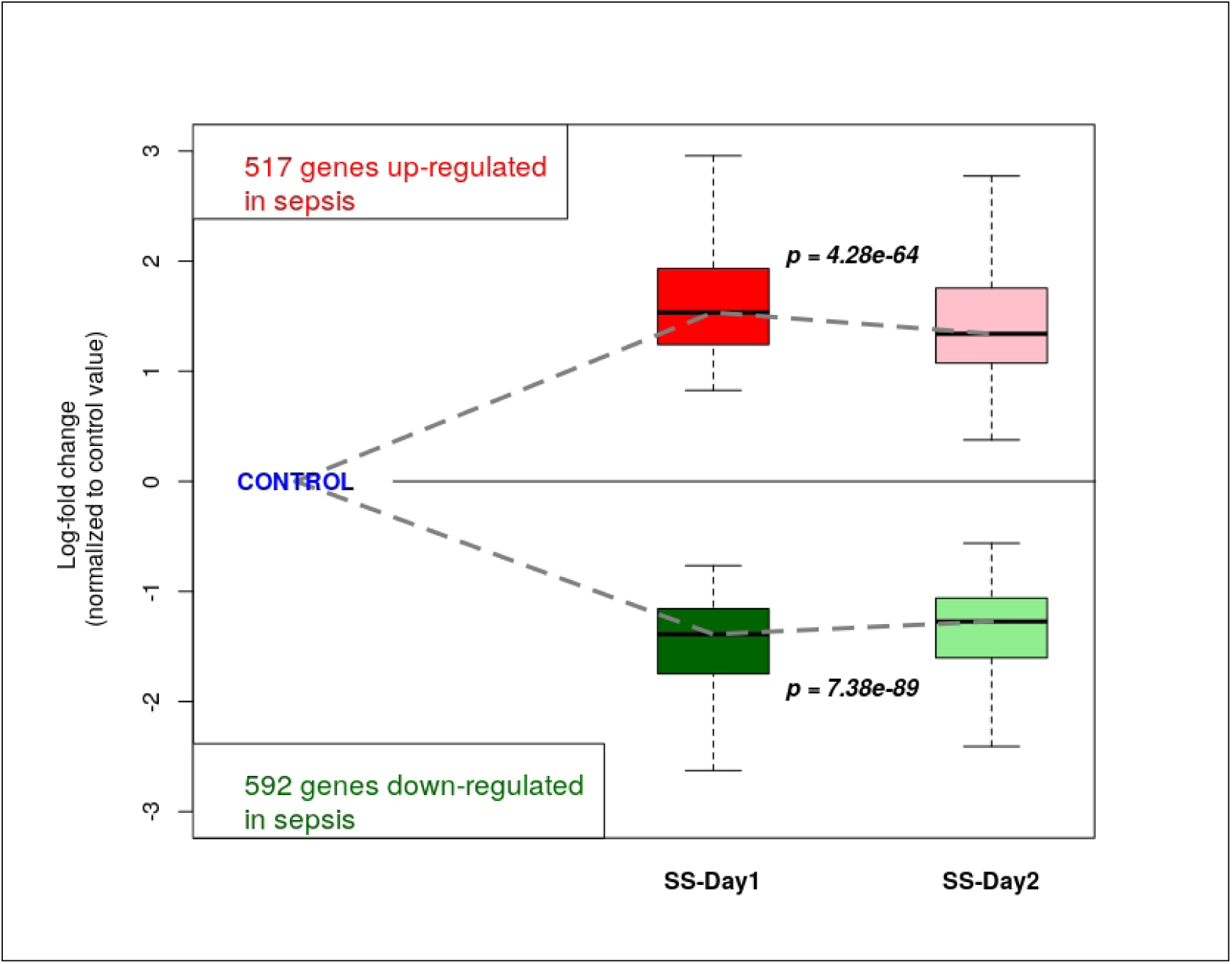
Temporal change of DE genes: Temporal change of DE genes (FDR p *<* 0.05, 2 fold-change or more), there is a non-random trend toward the baseline with time (p-values from paired t-tests are provided). This is consistent with earlier findings from patients with trauma [9].

### 3.3 Pathways associated with disease

To extract higher-order signal, we conducted analysis at the level of “gene sets” (pathways). Three complementary approaches were pursued: (over-representation) analysis based on hypergeometric test (ORA), permutation-based gene set enrichment analysis (GSEA) and topology based perturbation analysis. The intersection from the three analyses consisted of the pathways Osteoclast Differentiation, Antigen Processing and Presentation (AgPP) and T-cell Receptor (TCR) signalling (Table 1, Figure 4).

**Table 1.** Key pathways observed to be perturbed in SS compared to control.

| KEGG ID | KEGG pathway name | Direction of change in SS compared to Control | p value |
| --- | --- | --- | --- |
| hsa04612 | Antigen processing and presentation (AgPP) | Down-regulated | 3.51e-05 |
| hsa04660 | T-cell receptor signaling (TCR) | Down-regulated | 5.82e-07 |
| hsa04380 | Osteoclast differentiation | Up-regulated | 2.91e-06 |

**Figure 4.**
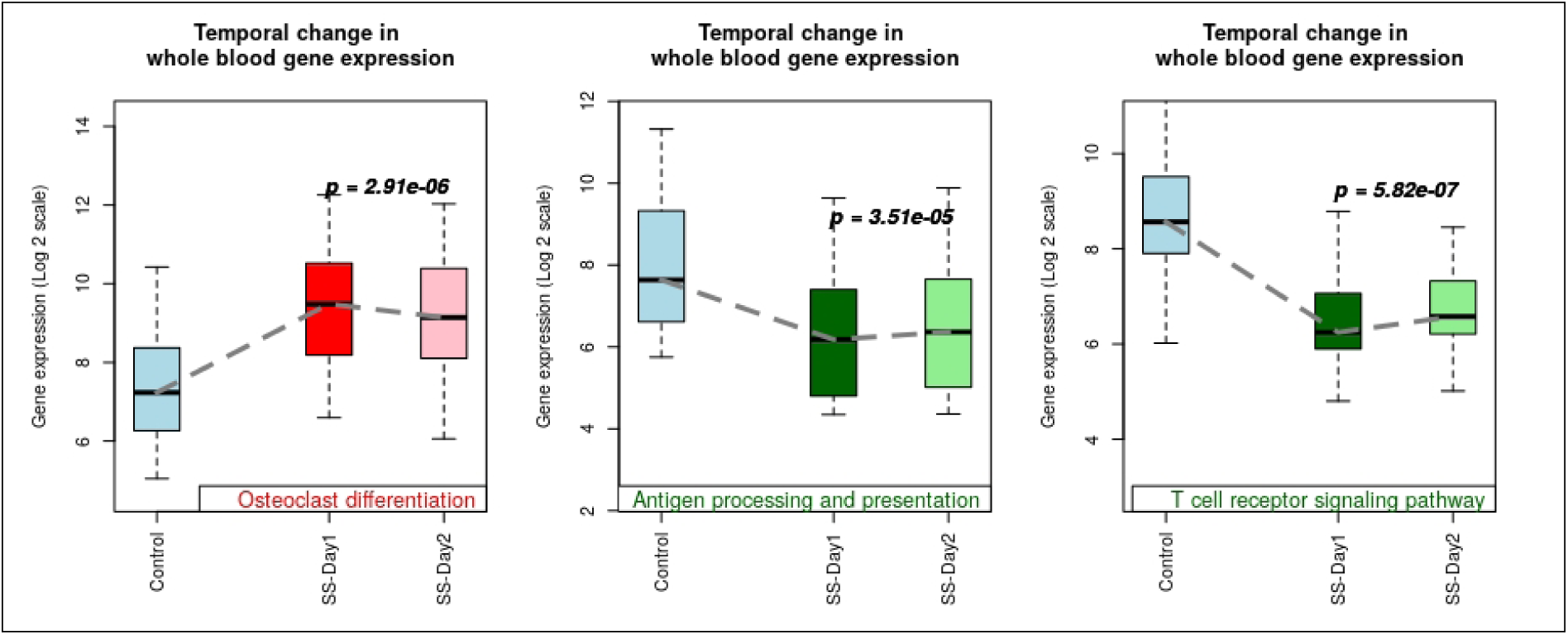
Pathways perturbed in disease: Each box represents median expression of genes in the given pathway. Data are plotted for three time points. “Control” (Healthy individuals), 1^st^ day of sepsis (SS-Day1), 2^nd^ day of sepsis (SS-Day2). The dotted line connects the median values of the three boxes and represents the trajectory of the pathway gene expression during sepsis.

### 3.4 Pathways associated with poor outcome in sepsis

Testing for differential expression in non-survivors compared to survivors revealed 1333 genes (7.6% of the genome) to be associated with survival (p *<* 0.05). Compared to the extent of transcriptional alteration in disease process, difference between survivors and non-survivors of sepsis was small, in terms of both the number of genes significantly affected (7.6% with survival compared to the 24% with sepsis) and magnitude of fold-change. Combined over-representation and perturbation analysis [16] revealed three pathways (Table 2) associated with survival. Each pathway was interrogated further as described below.

**Table 2.** Key pathways observed to be down-regulated in non-survivors in SCB cohort compared to survivors.

| KEGG ID | KEGG pathway name | Direction of change in Non-survivors | p value |
| --- | --- | --- | --- |
| hsa04064 | NF-kB signalling | Down-regulated | 3.5e-03 |
| hsa04612 | Antigen processing and presentation (AgPP) | Down-regulated | 3.6e-09 |
| hsa04660 | T-cell receptor signalling (TCR) | Down-regulated | 9.33-05 |

### 3.5 NF-*κ*B signalling pathway

NF-*κ*B signalling pathway is known to be associated with cell survival, immunity and inflammation. This pathway was observed to be significantly down-regulated in non-survivors compared to survivors (p = 0.0012) in SCB cohort as well as in other cohorts (Figure 5, Figure S5). Next, we considered the role of NF-*κ*B, a transcription factor in regulation of several genes of this pathway. We observed that the target genes of NF-*κ*B (involved in antigen presentation, T-cell activity and macrophage activation) were significantly (p = 0.004) down-regulated (Figure 8) in non-survivors. The differential impact of NF-*κ*B on host immunity was tested through relative difference in expression of genes associated with different macrophage types i.e. M1-specific pro-inflammatory genes and M2-specific hypo-inflammatory genes. Downregulation of M2-specific gene expression (p = 0.07) was observed in non-survivors of SCB cohort while M1-specific gene expression was observed to be up-regulated in survivors.

**Figure 5.**
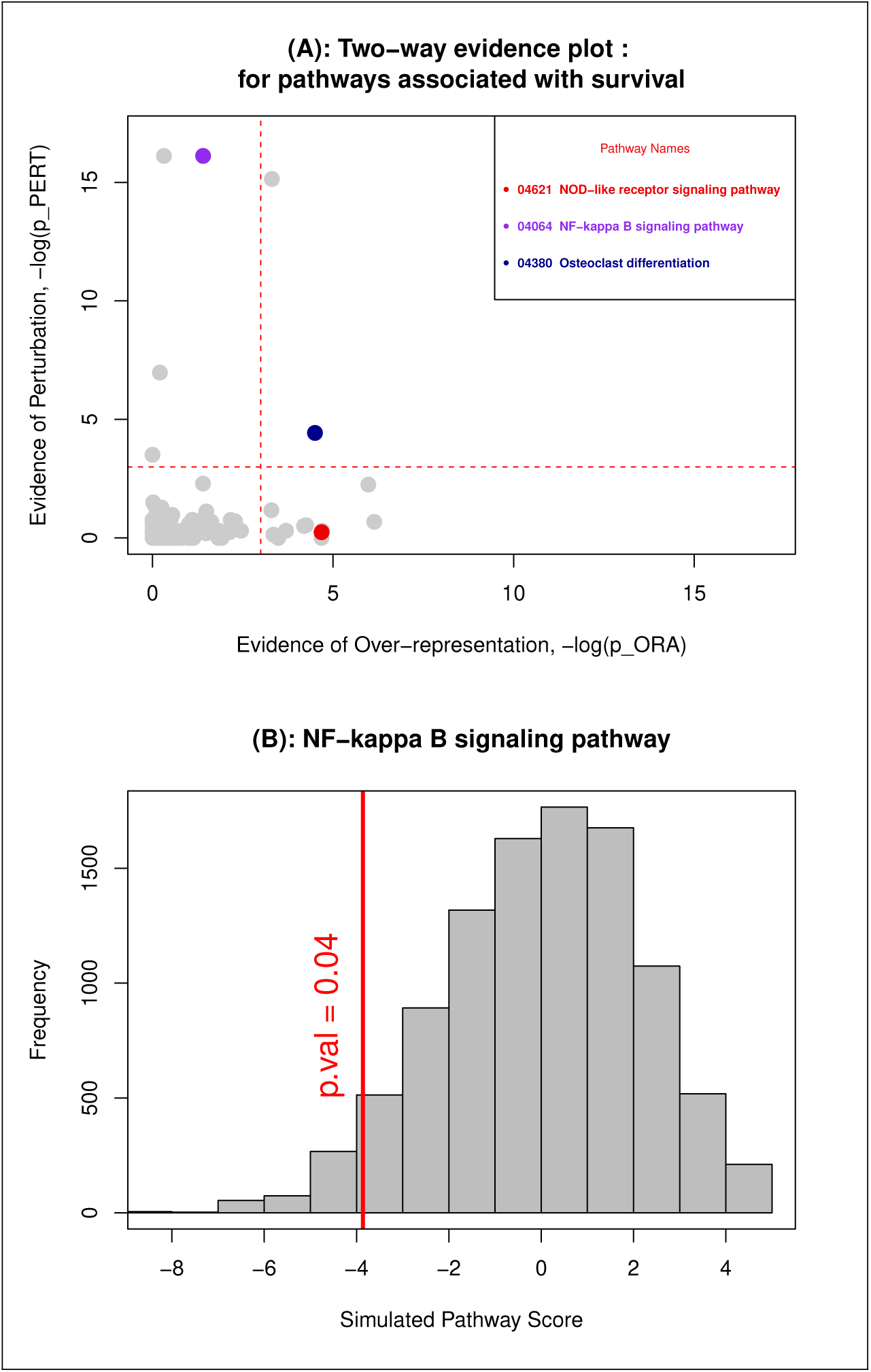
Evidence of perturbation of NF-*κ*B signalling pathway: NF-*κ*B signalling pathway is associated with survival. (A) NF-*κ*B signalling pathway, Osteoclast differentiation and NOD-like receptor signalling pathway. Each point in this plot represents a single KEGG pathway. The horizontal axis records over-representation (negative log p-value) evidence and the vertical axis records the perturbation (negative log p-value) evidence. The three pathways mentioned here are all significant when both the evidences are combined (i.e., Fisher’s product of the p-values). This plot is obtained from analysing data from multiple cohorts published earlier. (B) NF-*κ*B pathway was also observed to be significantly perturbed in the SCB cohort by permutation based GSEA method with significant difference between survivors and non-survivors (p = 0.04).

**Figure 8.**
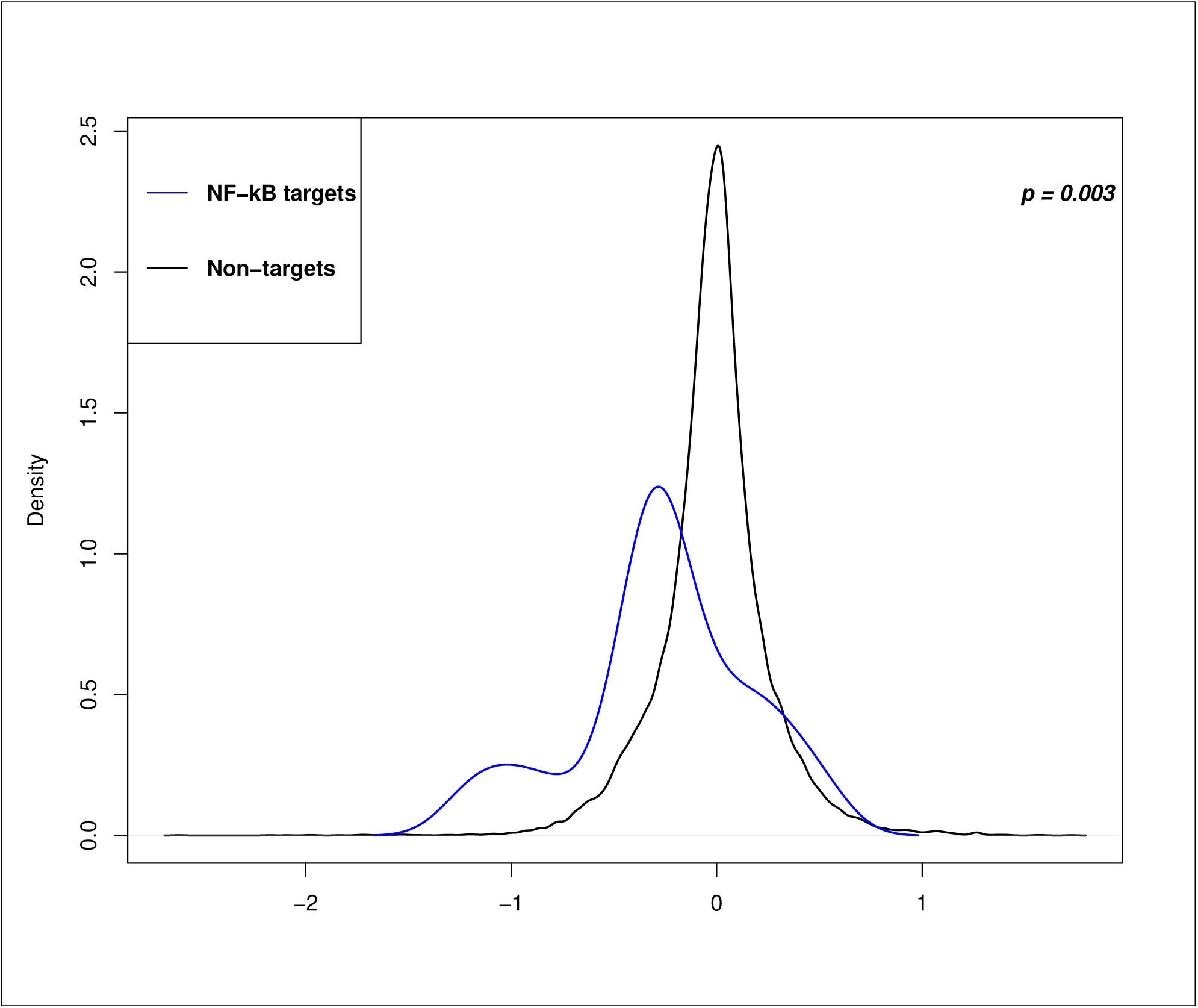
NF-*κ*B target gene expression density plot: A density plot of targets of NF-*κ*B between non-survivors and survivors that include antigen processing and presentation genes and various immune receptor genes. The gray peak in the background represents all the non-target genes in the genome and the blue peak represents the NF-*κ*B targets. There is a shift in the target gene expression to the left (p= 0.005), suggesting NF-*κ*B-induced down-regulation of its targets.

### 3.6 Antigen processing and presentation (AgPP)

This pathway was observed to be down-regulated in non-survivors compared to survivors (Figure S6). MHC class II genes (Table S3) of this pathway were found to be down-regulated in non-survivors compared to survivors, suggesting a role in impaired adaptive immunity functions (potentially mediated by T helper cells) in non-survival.

### 3.7 T-cell receptor signalling (TCR)

The T-cell receptor (TCR) signalling pathway was observed to be down-regulated in survivors compared to control, and it was further down-regulated in non-survivors (Table 1; Figure S7). Down-regulation of this pathway (along with antigen presentation) in non-survivors is consistent with the association of diminished adaptive immune response with poor outcome.

### 3.8 Transcriptional modules underlying survival

Following the principle of monotonic change across disease severity, KEGG pathways [10] were reconstructed into three transcriptional modules associated with SS. Each of the three modules was observed to be significantly (*p <* 0.05) associated with outcome (Figure 6) and immunosuppression was observed to be the module with the highest magnitude of change (Figure S8). A detailed description of modules is provided in Supplementary Text 2.

**Figure 6.**
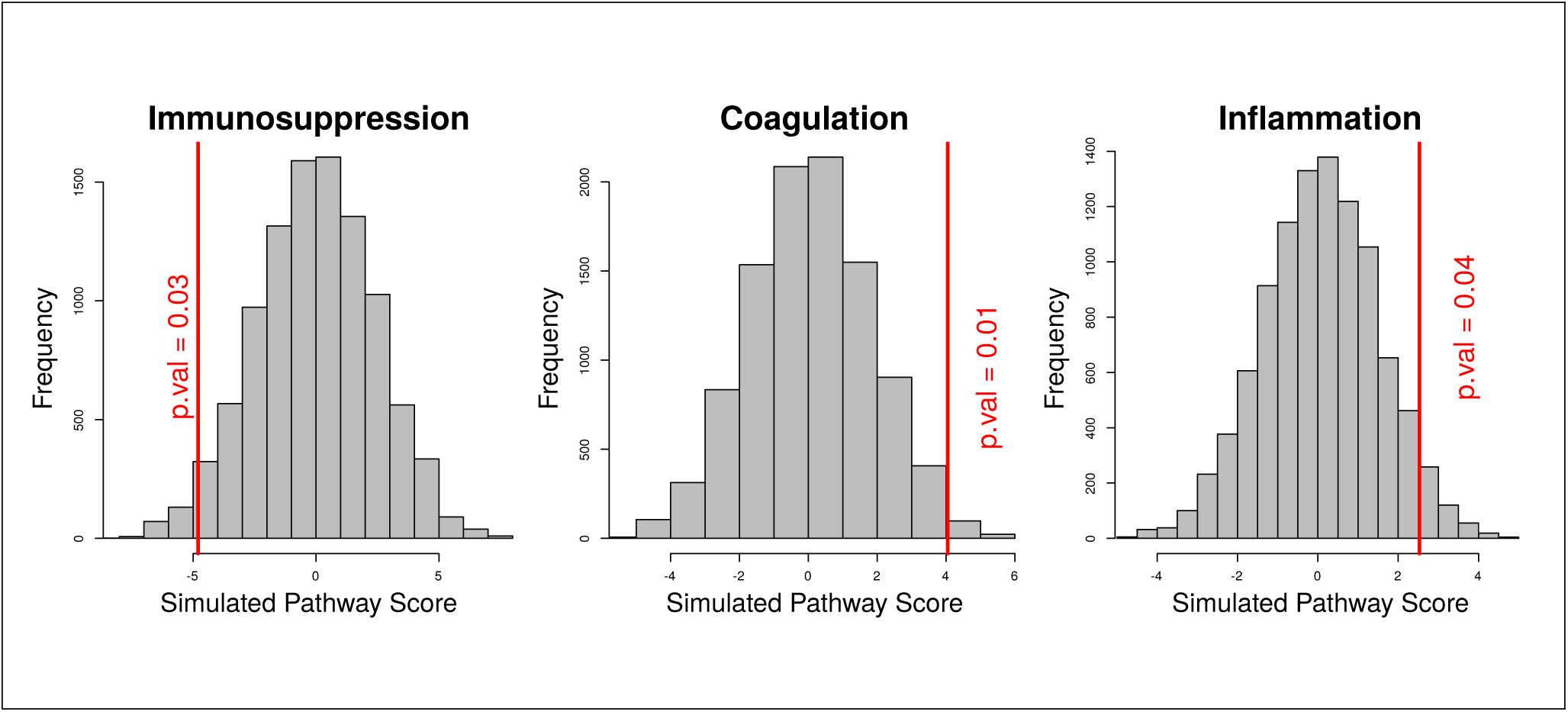
Significance of 3 biological modules: Histogram of simulated scores (GSEA permutation, 10000 times) and observed score (in red straight line) for three key biological modules. The histogram represents the null distribution of the gene set perturbation score. The deviation of the observed score for this gene set from the histogram suggests association between the gene sets and the disease outcome (non-survival). Significance of the module perturbation is denoted by the p-value.

#### 3.8.1 Immunosuppression

This module, is derived from the KEGG pathways: Antigen processing and presentation; T-cell receptor signalling and NK cell mediated cytotoxicity. Each of these pathways was observed to be significantly down-regulated in survivors compared to control, and further down in non-survivors (monotonic decrease). Antigen processing and presentation and T cell receptor mediated pathway genes were observed to be significantly down-regulated (GSEA p value 0.02 and 0.09 respectively) in non-survivors compared to survivors (Figure S6, S7). Both MHC class I and MHC class II genes were observed to be down-regulated (Table S3).

**Figure 7.**
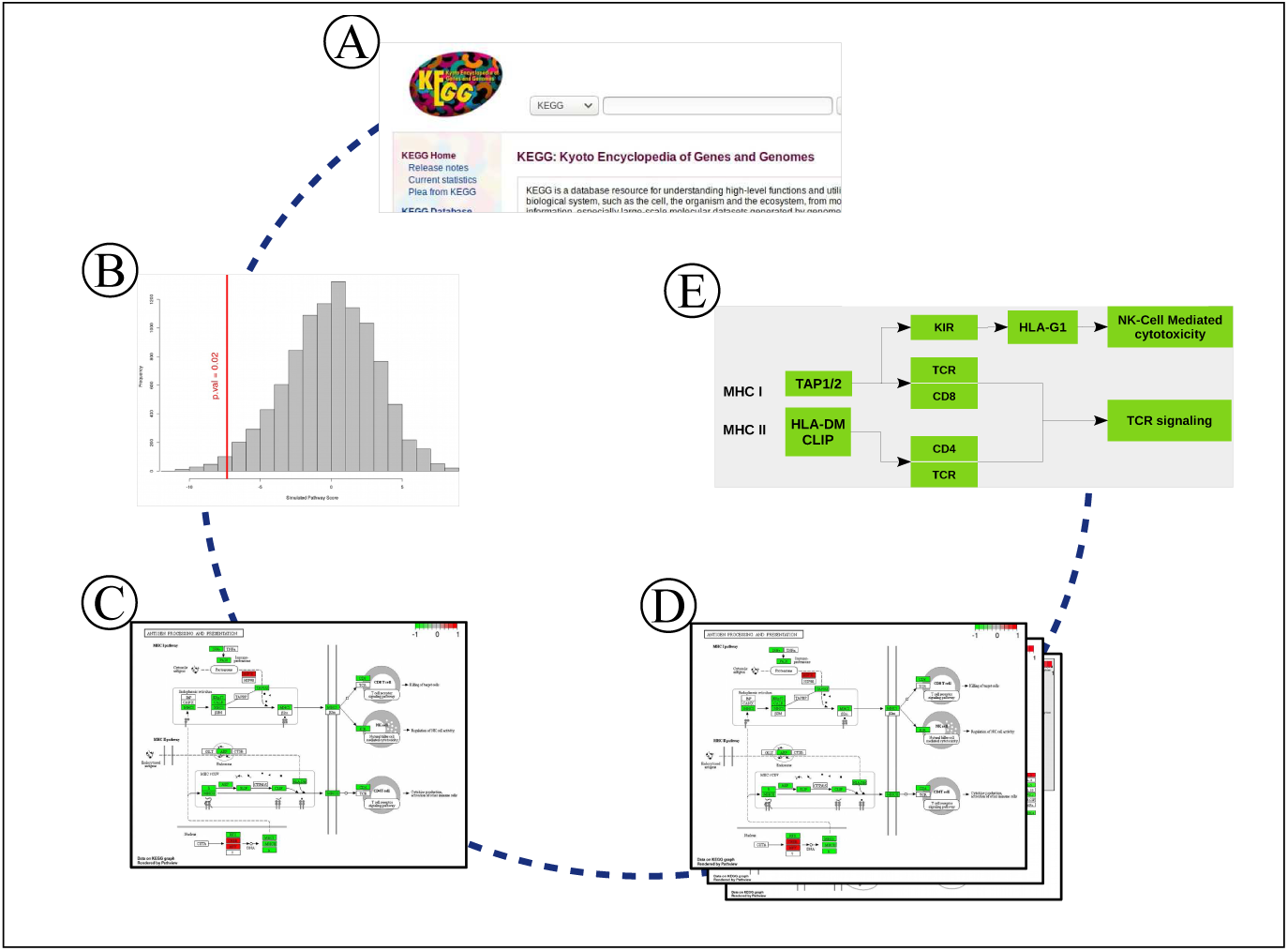
Steps of module construction: The sequence of steps leading to the construction of the three biological modules. (A) Selection of key biological processes (e. g. Immune pathways, Signalling pathways, Metabolic pathways) from KEGG database. (B) Testing for significant perturbation of the pathway in non-survivors. (C) Selection of pathways with genes differentially expressed and monotonic change from control through survivors to non-survivors. Finding the pathways which are monotonically changed. (D) Merging the monotonic pathways into a set of essentially connected nodes. (E) Construction of three key transcriptional modules (one is displayed here).

#### 3.8.2 Coagulation

A well known key pathophysiological characteristics of sepsis is coagulopathy. A pro-coagulant state is observed in septic patients due to the interaction of pro-inflammatory cytokines and tissue factor [11]. We observed Factor III, V, VII, XI to be up-regulated monotonically from healthy to survivors and further up-regulated in non-survivors, suggesting a more severe coagulopathy for those patients that did not survive.

#### 3.8.3 Inflammation

We observed several genes associated with inflammation to be monotonically up-regulated from control through survivors to non-survivors. Up-regulation of many chemokines (TGF-*β*, IL-13, IL-4, IL-6, and IL-21) and key signalling molecules (PI3K/AKT, PKC, WASP, ROCK) are associated with regulation of actin cytoskeleton (GSEA, p value = 0.04) (which helps in trans-endothelial migration of leukocytes).

## 4 Discussion

Unbiased and stringent testing revealed about one-fourth of the genome to be transcriptionally altered in day 1 of SS compared to control (Figure 2). This profound change is induced by the disease state, and is comparable with the scale of change generally observed in critical illness or sepsis (Table S1). With passage of time in the ICU, the transcriptome slowly returns to the baseline, with a small but significant difference between day 1 and day 2 of SS (Figure 3). Of note, there is a deviation in the trajectories of non-survivors compared to survivors, with the magnitude of differential expression being greater in non-survivors at both the time points. In general, delayed genomic trajectory in non-survivors suggests an increased disease severity that resists restoration of baseline expression. This trend is part of the generic host response seen in critically ill humans [9]. We have leveraged this observation in the quest for transcriptional modules with stable association with outcome.

Functionally, there is evidence of up-regulation of osteoclast differentiation and down-regulation of antigen processing and presentation, T-cell receptor signaling (Table 2). While immuno-suppression continues to be a recurring theme in sepsis biology, osteoclast differentiation has only recently been reported in our previous work in the context of septic shock with some evidence of its association with survival [12].

Immunosuppression in our patients is a result of down-regulation of NF-*κ*B signalling in non-survivors as established by stringent testing (Figure 5). Monotonic decrease in non-survivors, i.e., positive correlation between disease severity and differential expression, suggests that this dysregulation is a stable consequence of SS. Targets of NF-*κ*B include molecules of the immune system, for example, those associated with T-cell receptor signaling, antigen presentation and alternative macrophage activation. These are significantly down-regulated in non-survivors (Figure 8-9; Table 2). Similarly, NF-*κ*B-mediated down-regulation of M2-type genes (Figure 9) was consistent with an immunosuppressed state in the non-survivors. In spite of the tremendous interest in the role of NF-*κ*B in inflammation in general and sepsis in particular [13, 14], NF-*κ*B has resisted being a successful target of sepsis therapy [15–18]. Further investigation is required to improve our understanding of the precise role of NF-*κ*B in determining the outcome of SS.

**Figure 9.**
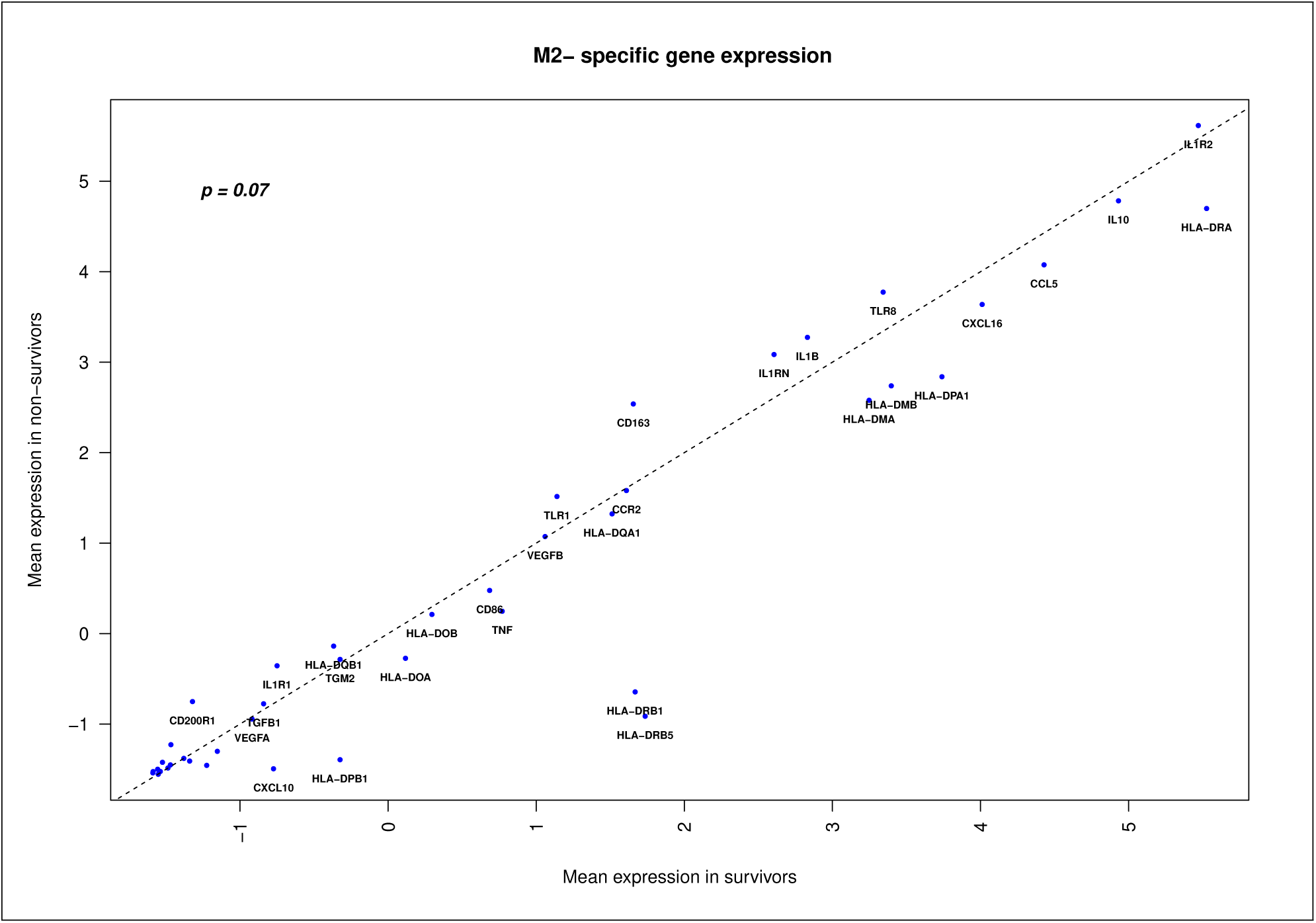
M2-specific gene expression: Scatterplot of M2 target gene expression in non-survivors compared to survivors. Each point represents a single gene with mean expression in the two groups of patients: survivors (x-axis) and non-survivors (y-axis). For most of the genes there is higher mean expression in survivors, suggesting significant M2-specific (p = 0.07) under-expression in non-survivors.

Monotonic change across disease severity spectrum (healthy control, survivors, non-survivors) has the potential to better define molecular underpinning of SS outcome. This is true not only at the individual gene level but also at the level of functionally connected gene sets, i.e., transcriptional modules (Figure 7). The modules correspond to the three major dysregulated pathophysiologic processes considered part of the hallmark of sepsis: immunosuppression, coagulopathy and inflammation. These transcriptional modules capture progressive dysfunctional processes in SS, culminating in poor outcome (Figure 6). Although each of the three modules is significantly different between survivors and non-survivors, the magnitude of immunosuppression is greater than that of coagulation, with least perturbation for the inflammation module (Figure S8).

Our finding raises questions for management of patients in this region. If validated on a larger set of patients, it appears that immune-enhancement e.g., immune adjuvant therapy [19]) shall be a better strategy than inhibition of inflammation.

One caveat is that this study included a small cohort of samples (n=40). Therefore, the results must be considered preliminary in nature. Secondly all our patients were recruited from a single clinical center. In order to adequately capture the demographic diversity of the region, a future multi-centric study shall need to include patients from different geographical locations across the country.

## 5 Conclusion

Profound transcriptional reprogramming in SS was interrogated by complementary bioinformatic strategies revealing down-regulation of immune pathways. NF-*κ*B appears to be at the center of this change. Data-driven integration of KEGG annotation and sepsis pathophysiology led to identification of three key transcriptional modules associated with survival – immunosuppression, coagulopathy and inflammation. Quantitatively, the magnitude of immunosuppression is much more than that of inflammation, with potential role in patient stratification. The three modules (containing 288 genes) can be tested (with the help of quick gene expression profiling technologies at bedside) to determine which category each patient belongs. Specific therapy can then be catered to each patient based on that categorisation.

## 6 Materials and methods

### 6.1 Differential gene expression analysis

Welch t-test [20] was performed to detect genes DE in SS compared to gender and age matched healthy controls. Correction for multiple testing was performed to reduce False Discovery Rate (FDR) [21]. Threshold of FDR p-value of *<* 0.01 and 2-fold or greater change; was applied to detect DE genes.

### 6.2 Pathway analysis methods

#### 6.2.1 Over-Representation Analysis (ORA)

Significantly up-regulated genes were subjected to over-representation analysis by applying hypergeometric test [22]. Correction for multiple testing was performed to reduce FDR. A significant association was detected at threshold of FDR p *<* 0.001.

#### 6.2.2 Gene Set Enrichment Analysis (GSEA)

Permutation test was performed by randomly scrambling the sample labels (case/control) and computing the enrichment score (t-statistics) of the gene set for this permuted data set [23]. Multiple rounds of this process generated the null-distribution of the enrichment score. Pathways with observed enrichment score significantly deviated (FDR p *<* 0.05) from the null distribution were considered significant [23].

#### 6.2.3 Perturbation Analysis of Signalling Pathways

Analysis based on pathway topology (derived from [24]) revealed the probability of perturbation of the pathway. In this process the positions of the differentially expressed genes are scrambled in the pathway and for each combination a perturbation score is calculated. Ultimately a total perturbation score is calculated by summing up the cumulative perturbation score from each iteration of gene swapping. In our analysis perturbation score was applied along with over-representation score visualise pathways (Figure 5). By combining two different evidences (one from the hypergeometric test model and the other from the probability of perturbation that takes the pathway topology into account), a two way evidence visualisation was made possible [24].

### 6.3 Construction of Biological Modules

We reconstructed three modules (immunosuppression, coagulation and inflammation) from KEGG pathways associated with SS (Figure 7). First, we selected genes (nodes) from significantly perturbed pathways, that were also monotonically dysregulated from baseline in survivors and further in non-survivors. We reasoned that the extent of perturbation of biological process shall increase with severity of disease, and therefore, the genes that are monotonically changing are key nodes of outcome-associated modules. Then we extended the network of genes based on the connectivity as described in a well-curated pathway database (KEGG [10]). Wherever possible, we made the connectivity parsimonious with minimal number of non-significant genes (included only to preserve the continuity of signal flow, in keeping with the directed nature of curated KEGG graph). Three modules were constructed to describe three aspects of sepsis pathophysiology – inflammation, coagulation and immunosuppression. Each module was then considered an independent gene set and tested for significance of transcriptional difference between non-survivors and survivors by permutation-based testing of pathway score.

### 6.4 Selection of data sets from GEO

Electronic search was performed in NCBI GEO (Gene Expression Omnibus) on 06th December 2017, with the search string: “sepsis or septic shock or survivor or non-survivor”. Application of human filter (species-human) resulted in 9 transcriptome data sets (Figure S9; Table S4). One study (GSE78929) was rejected as it contained data from a tissue (muscle tissue) and not blood. Blood was chosen as it captures the systemic cellular response in systemic inflammatory disease such as sepsis [25]. Hierarchical clustering of the gene expression data led to clear age-specific segregation of sepsis transcriptome (Figure S10). The data from our SCB cohort consisted only of adult sepsis cases. Accordingly, all subsequent analysis for validation was performed on adult data sets only.

### 6.5 SCB (Srirama Chandra Bhanja Medical College) cohort

Transcriptomic profiling was performed on sepsis cases and on matched (by age and gender) healthy controls, who were not suffering from any inflammatory diseases, and were not related to the patients. Samples were collected at two different time points in the cases. A comparison between survivors and non-survivors was planned to identify genomic changes that are specifically associated with survival. Patients suspected of sepsis were recruited into the study with defined inclusion and exclusion criteria mentioned in Table S2. Two blood samples from the sepsis cases were collected, the first at the time of diagnosis (D1), and a second sample after 24 hours (D2). Representative single blood sample was collected from each of the healthy control individuals. Blood samples were collected from the patients of SS and healthy subjects after obtaining approval from the Institutional Ethical Committees of the National Institute of Biomedical Genomics, Kalyani and SCB Medical College, Cuttack. All the methods were carried out in accordance with the approved guidelines. Informed written consent was obtained from all subjects who participated in the study. A total of 27 patients (23 with paired transcriptome data for two time points) and 12 healthy control subjects (for the single time point) were recruited in this study. For each of the cases and matched control samples quality of RNA (isolated from whole blood) was assessed by the following criteria: ratio of absorbance of light at two wavelengths (A_260_/A_280_) should be between 1.8 - 2 and RNA Integrity Number (RIN) of 6 or higher. Another criterion for selection of subjects was based on temporal analysis, which requires that the two temporal samples from each subject should pass the quality assessment mentioned above.

#### 6.5.1 Sample processing and quality assessment

Venous blood was collected from patients in PAXgene Blood collection tubes from BD, Franklin Lakes, New Jersey, USA (cat. no. 762165), containing stabilising reagent that keeps the blood cells fixed and preserves cellular RNA before further use. PAXgene Blood RNA isolation Kit from Qiagen, Hilden, Germany (cat. no. 762164) was used to isolate the RNA from the whole blood according to manufacturer’s instructions. Spectroscopy was done in Nanodrop-2000 (ND-2000, from Thermo Fisher Scientific, Waltham, Massachusetts, USA) for checking RNA quality and concentration. Agilent Bioanalyzer RNA nano kit (Agilent RNA 6000 Nano) was used for checking RNA quality. Illumina TotalPrep RNA Amplification Kit was used to convert the total RNA to biotinylated cRNA for hybridisation to microarray chip. Transcriptome profiling was done on the following microarray platforms: Affymetrix (GeneChip^TM^ Human Gene 2.0 ST Array, cat. no. 902113) and Illumina Microarray Chips (HumanHT-12 v4 Expression BeadChip Kit, cat. no. BD-103-0204)

#### 6.5.2 Software

All analysis were done in R [26], a language and environment for statistical computing in Linux operating system.

#### 6.5.3 Data availability

A vignette with reproducible code chunks for analysis is provided in the supplementary data. All data and code are available at (https://figshare.com/) under project (“ssnibmgsurv”).

## 7 Author contributions

**SM:** contributed to Data Curation, Microarray Hybridization, Investigation, Analysis, Methodology, Software, Validation, Visualization, Writing, Reviewing and Editing of the original draft.

**PT:** contributed to Data Curation, Investigation, Reviewing the original draft.

**BD:** contributed to Data Curation, Investigation, Funding Acquisition, Project Administration, Resources, Supervision, Reviewing the original draft.

**SKM:** conceptualized the study and contributed to Funding Acquisition, Project Administration, Resources, Supervision, Analysis, Methodology, Software, Reviewing and editing of the original draft.

All authors contributed to Manuscript Reviewing and approved of the final manuscript.

## 8 Acknowledgments

This work was supported by an intramural grant from NIBMG, Kalyani and an extramural grant (No. BT/PR5548/MED/29/571/2012, duration of 3 years, sanctioned on 27-05-2013) from the Department of Biotechnology, Government of India. S.M. acknowledges research fellowship (No. F.2-45/2011(SA-I)) provided by the University Grants Commission of India. We are grateful to the patients of SCB Medical College Hospital for contributing blood samples for the study. We acknowledge the kind suggestion given by Prof. Partha P Majumder during the analysis of this data. We acknowledge the kind suggestions of Dr. Samsiddhi Bhattacharjee regarding the metaanalysis of published data. We acknowledge the kind suggestions by Prof. B. Ravindran for his kind discussion on many topics of sepsis. We thank our colleagues at the NIBMG core facility and the ILS laboratory for timely processing of the samples, often at short notice.

## 10 Supplementary Text 1

### 10.1 Immunostimulatory therapy in sepsis

Our result is supportive of Ono *et al*. [19] who argued in favour of immunostimulation over targeting inflammation as a possible therapeutic startegy to treat patients suffering from sepsis. As the immunosuppression involves chiefly the adaptive immune system, there have been attempts to boost adaptive immunity through IFN*γ*, granulocyte-macrophage colony-stimulating factor (GMCSF), or granulocyte colony-stimulating factor GCSF [27]. However these attempts have failed to demonstrate a clear survival benefit to those who received these therapies. A meta-analysis [28] reports that GCSF and GMCSF failed to show survival benefit in sepsis patients. Two Interleukins IL-7 [29] and IL-15 [30] have been shown to have adaptive immunostimulatory function in sepsis patients. IL7 treatment to sepsis patients restored IFN-gamma secretion and T-cell proliferation [29] in them. PD-1 is another interesting target molecule for immunostimulatory therapy, as it causes immunosuppression through IL-10 expression [19]. Anti PD-1 [31] and PDL-1 [32] therapy in bacterial and fungal murine sepsis showed increased survival. Disruption of the PD-1/PDL-1 axis seems to be a novel approach for restoring immune function in sepsis patients [19].

## 11 Supplementary Text 2

### Immunosuppression

This module (Figure S1) consists of MHC molecules with co-stimulators (CD4, CD8), transporters (TAP1/2), and receptors (KIR, TCR) resulting in NK-cell mediated cytotoxicity and T-cell receptor signaling. TAP1, TAP2 help in transport of the processed cytosolic pathogenic antigen with the help of MHC class I. NK cells are the part of innate immunity that detects the pathogen infected cell by MHC class I and interact through Killer-cell immunoglobulin-like receptors (KIRs) located in the plasma membrane of NK cells. The NK cells mostly play an instrumental role in removal of virus infected cells. MHC class II mediated antigen presentation, observed to be down-regulated in non-survivors leading to impaired TCR signalling with MHC class II and CD4, CD8 all were significantly down-regulated in non-survivors. This is consistent with general and perhaps reversible immune paralysis/ suppression in non-survivors.

**Figure S1.**
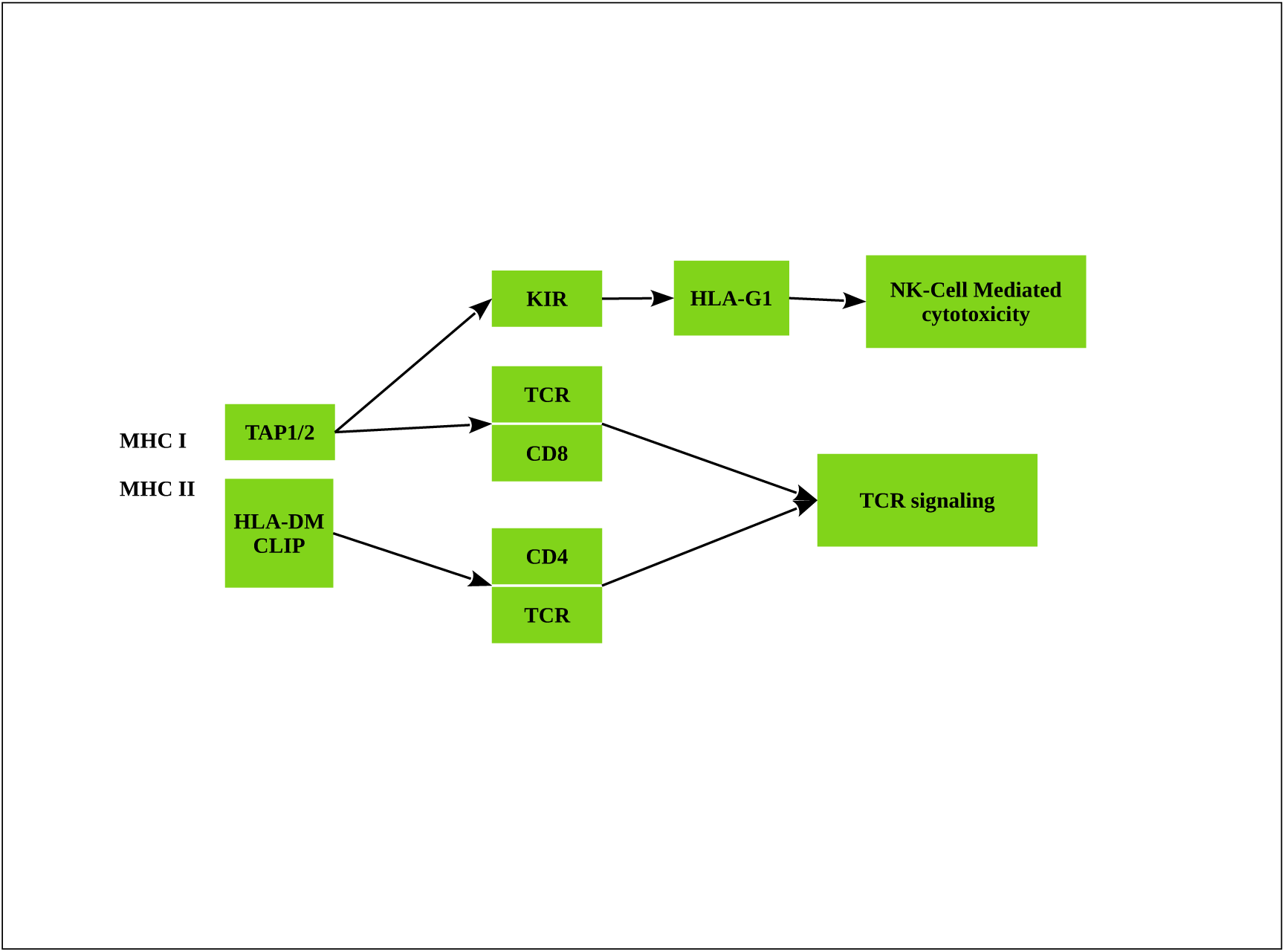
Immunosuppression module. A schematic representation of the connected nodes in the immunosuppression module.

### Coagulation

Platelets can detect vascular injury through collagen (up-regulated in non-survivors and promoting clot formation by platelet activation [33]. This module (Figure S2) consists of collagen-induced platelet activation and clotting factors both causing formation of fibrin and a persistent pro-coagulant phenotype. GPVI (up-regulated in non-survivors) is a key receptor protein that plays an important role in platelet activation after the collagen is bound to it. Up-regulation (GSEA p = 0.03) of extracellular matrix (ECM) pathways suggests damage to the ECM. This activates platelets to the site of injury that contribute to repair of the damaged tissue at that site [33].

**Figure S2.**
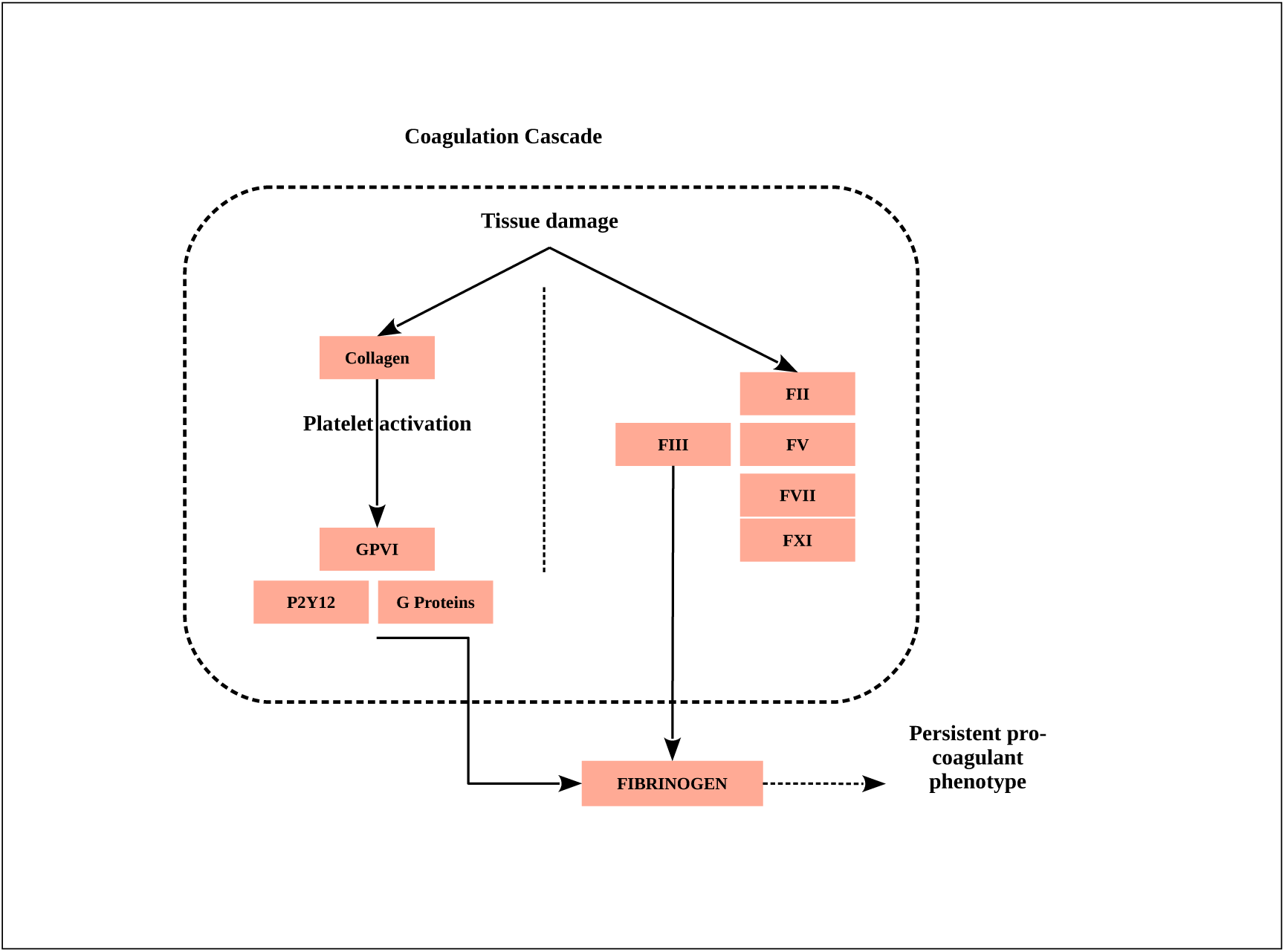
**Coagulation Module**: A schematic representation of the connected nodes in the coagulation module.

### Inflammation

This module (Figure S3) consists of pattern recognition receptors, chemokines, signaling molecules (PI3K/AKT, MAP Kinases), leading to regulation of actin cytoskeleton and leukocyte migration. As expected, there is up-regulation of genes associated with inflammation.

The main input signal of the inflammation emanates from not only the PAMPS (that recognise microbial antigens), but also the damage-associated molecular patterns (DAMPs); that indicates tissue damage due to host inflammation. Toll like receptors recognize pathogen-associated molecular patterns (PAMPs) [34] as well as DAMPs.

**Figure S3.**
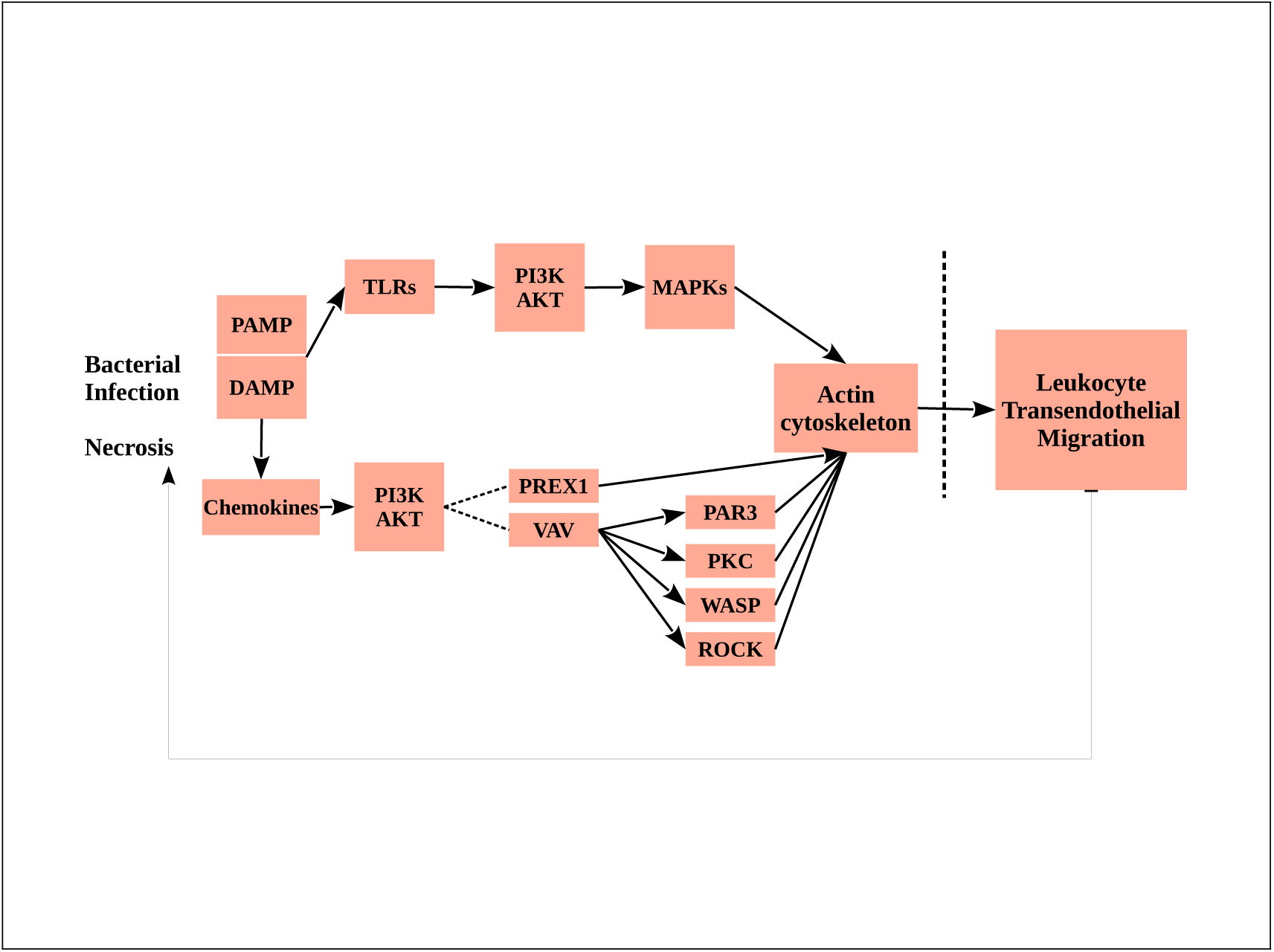
**Inflammation Module**: A schematic representation of the connected nodes of the inflammation module.

## 12 Supplementary Figures and Tables

**Figure S4.**
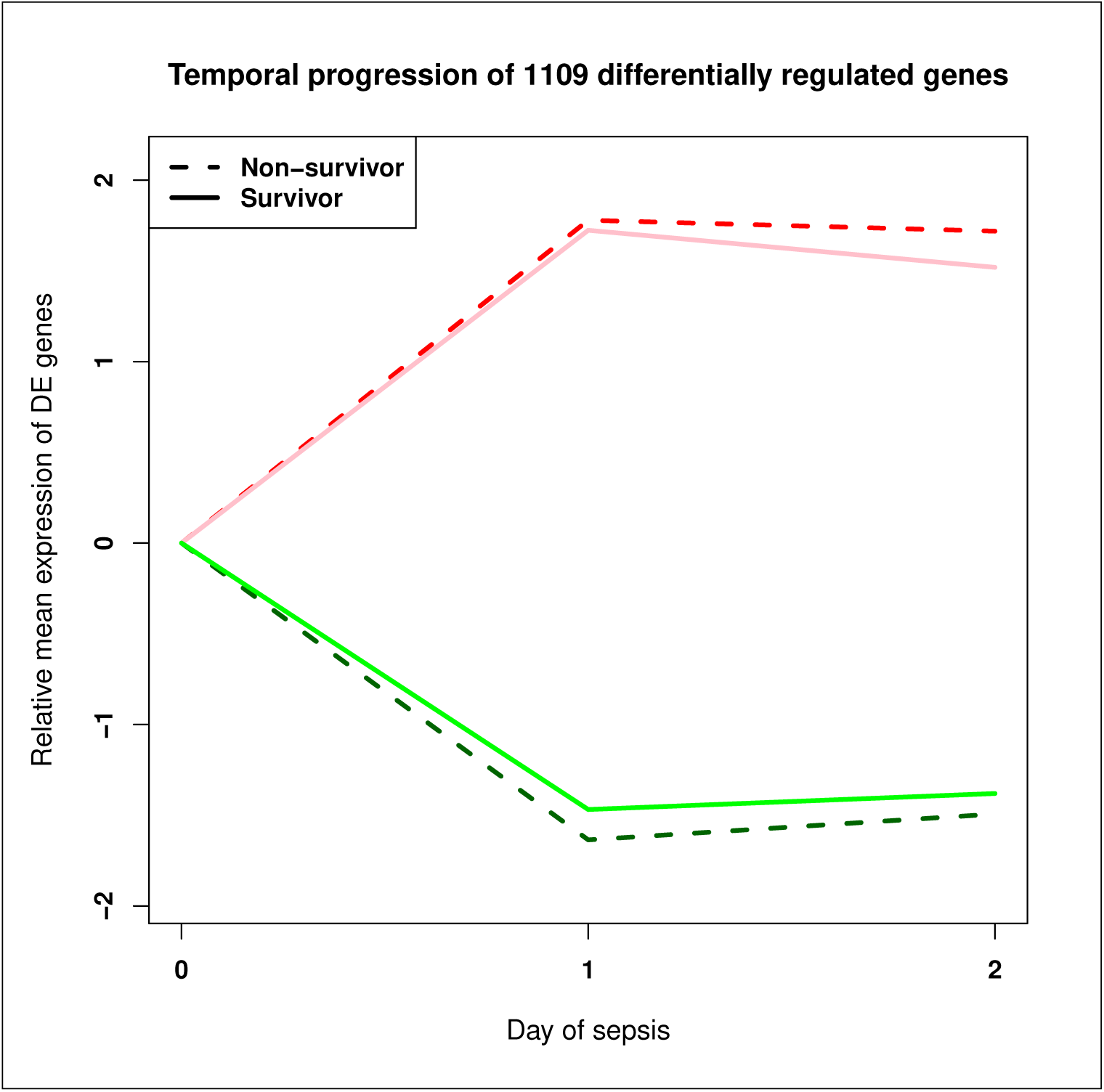
**Trajectory of DE genes:** This is a line plot of mean expression of DE genes in SS patients compared to control. Compared to the survivor group, generally the non-survivor gene expression is more deviated from controls and continues to diverge with time.

**Figure S5.**
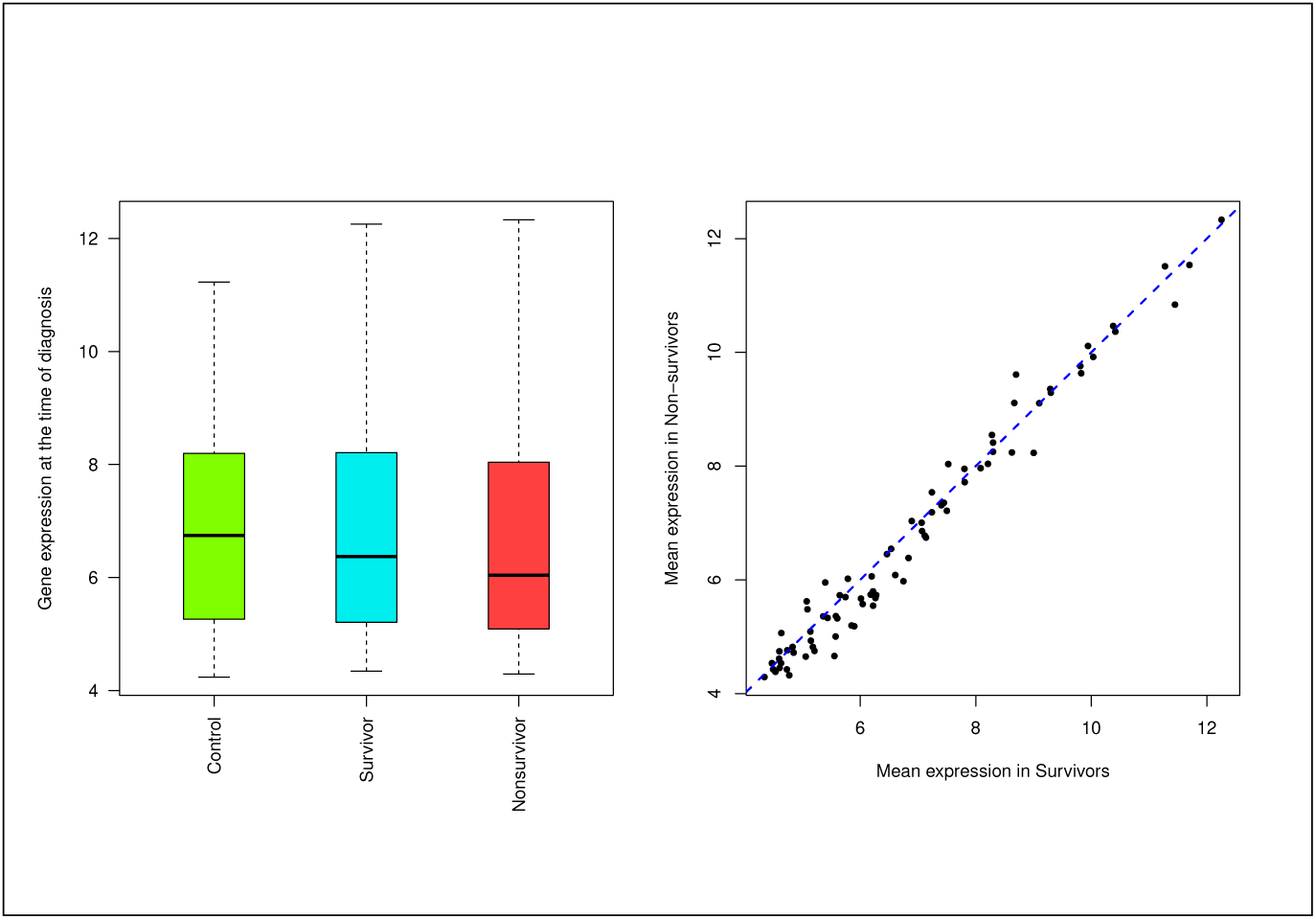
**NF-*κ*B pathway:** Boxplot and scatterplot of NF-*κ*B pathway in SCB cohort. The pathway is monotonically down-regulated from control through survivors to non-survivors on Day-1.

**Figure S6.**
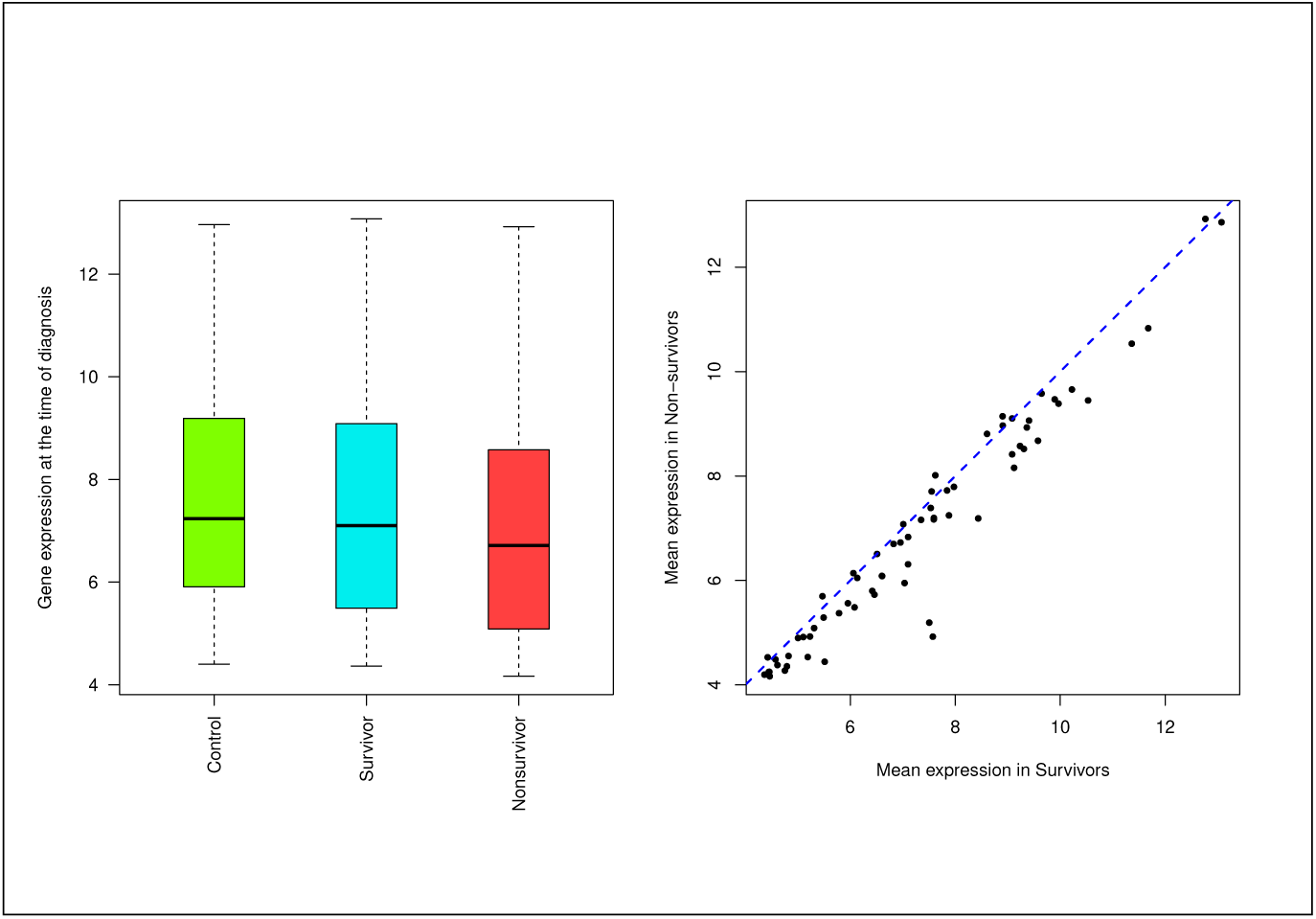
**Antigen Processing and Presentation Pathway:** Boxplot and scatterplot of Antigen Processing and Presentation Pathway pathway in SCB cohort.

**Figure S7.**
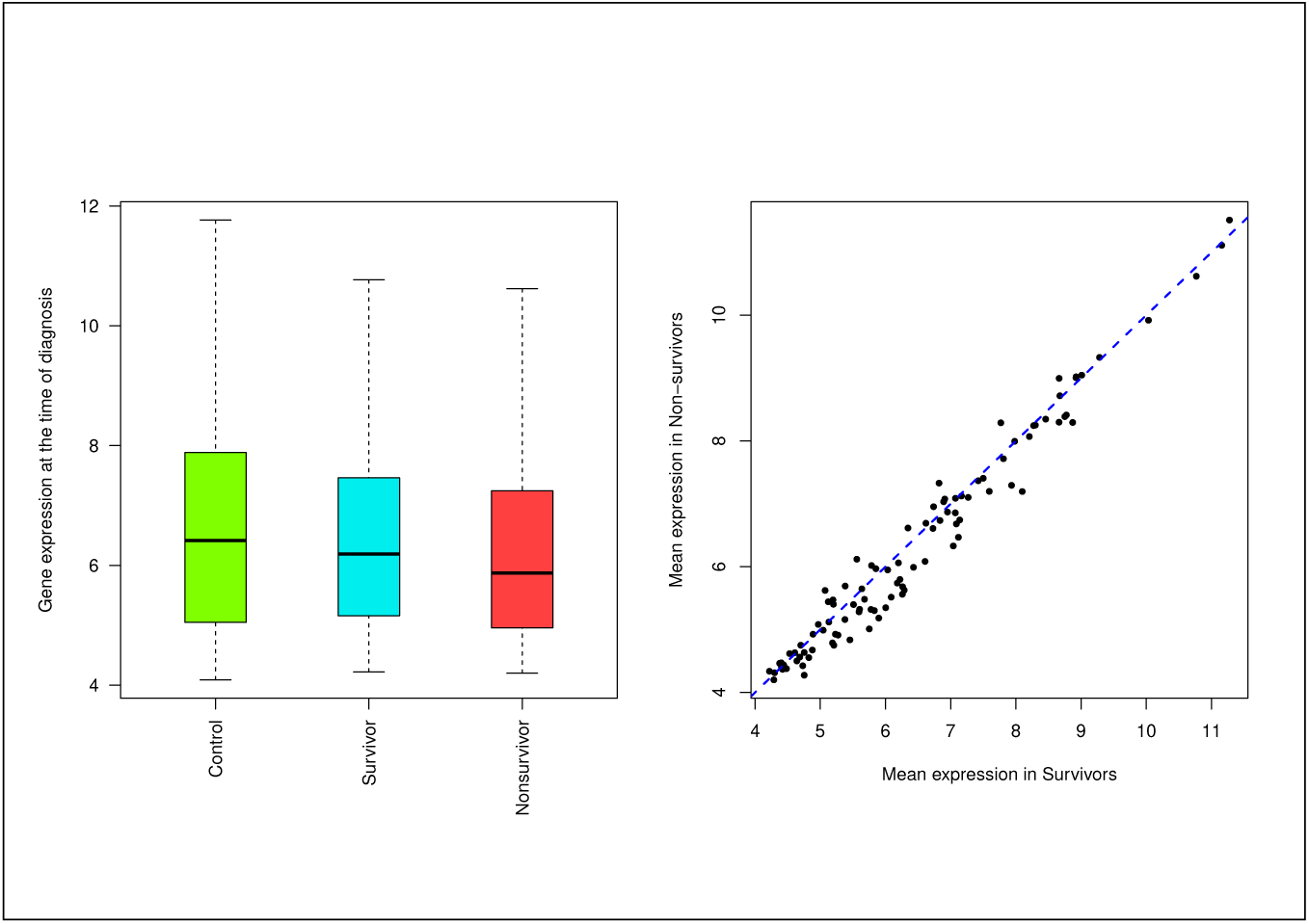
**TCR signaling Pathway:** Boxplot and scatterplot of TCR signaling pathway in SCB cohort. The pathway is monotonically down-regulated from control through survivors to non-survivors on Day-1.

**Figure S8.**
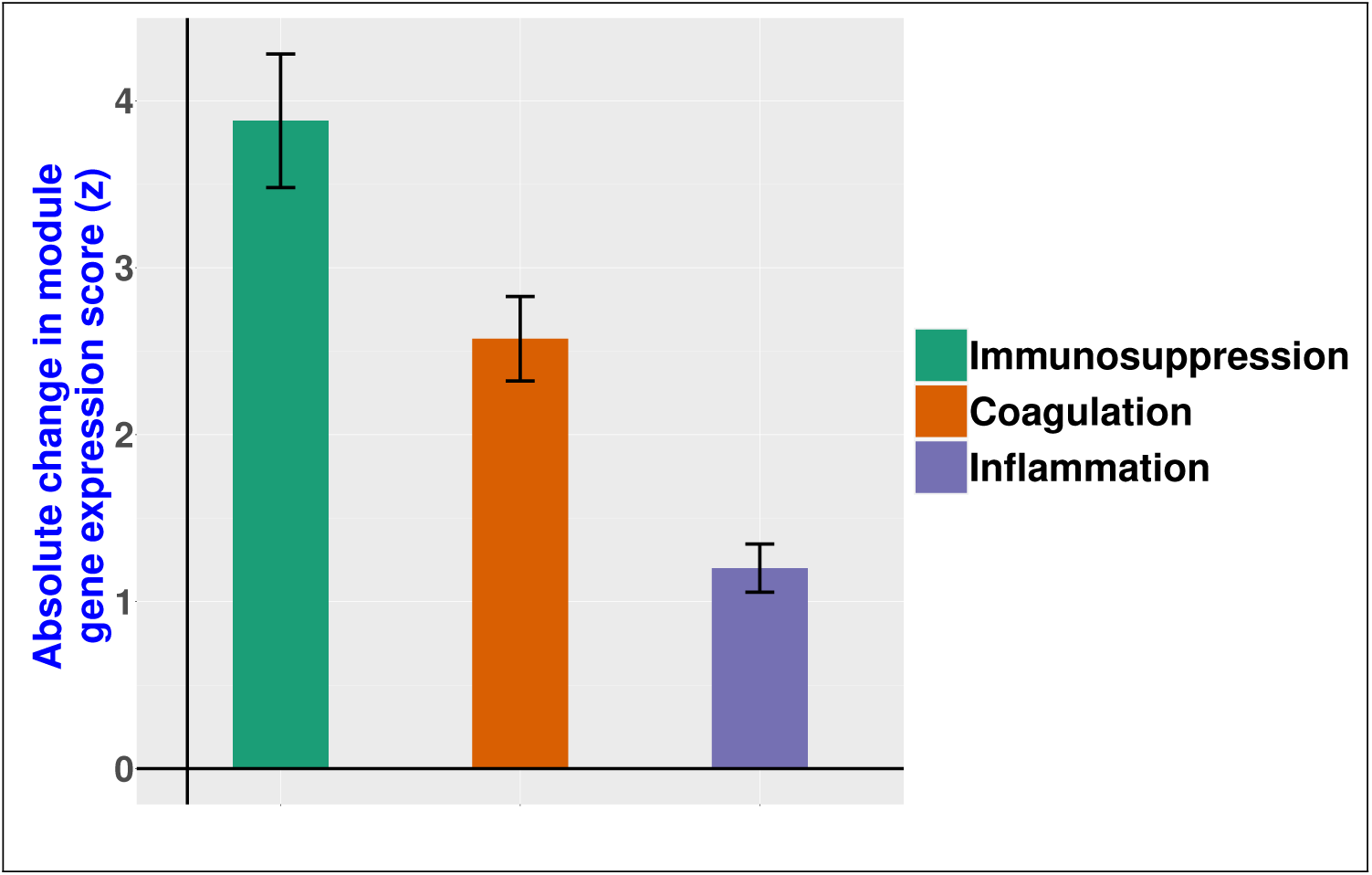
**Mean module score in SCB cohort:** Barplot shows the magnitude of gene expression of 3 key modules in each sepsis patients. For each module a gene expression score (z) was calculated to capture the relative expression of the genes of the module in the sepsis patients compared to the healthy subjects. Quantitatively, magnitude of change in expression is highest for the immunosuppresion module.

**Figure S9.**
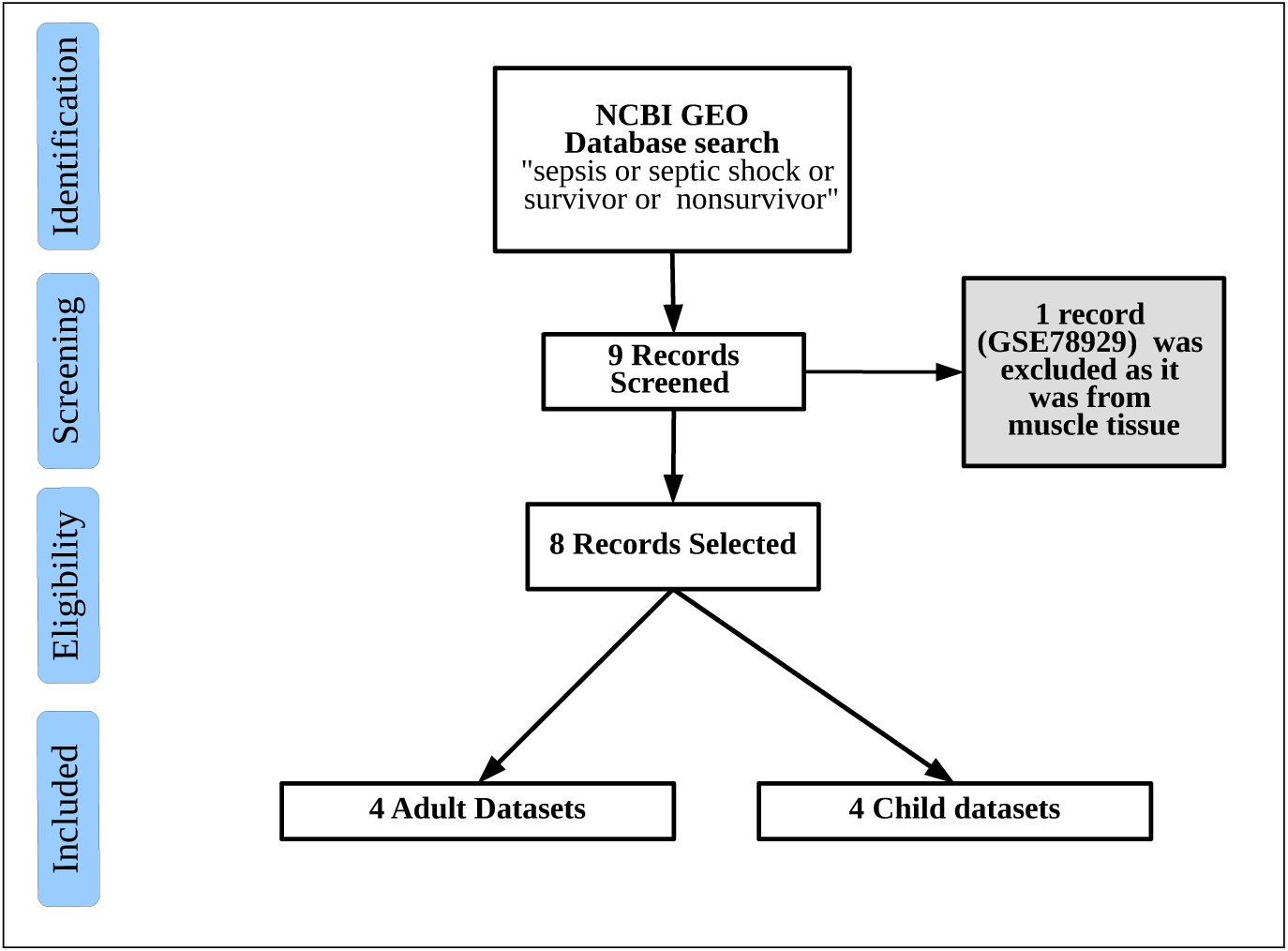
**PRISMA flowchart:** For selection and screening of the data sets from NCBI GEO.

**Figure S10.**
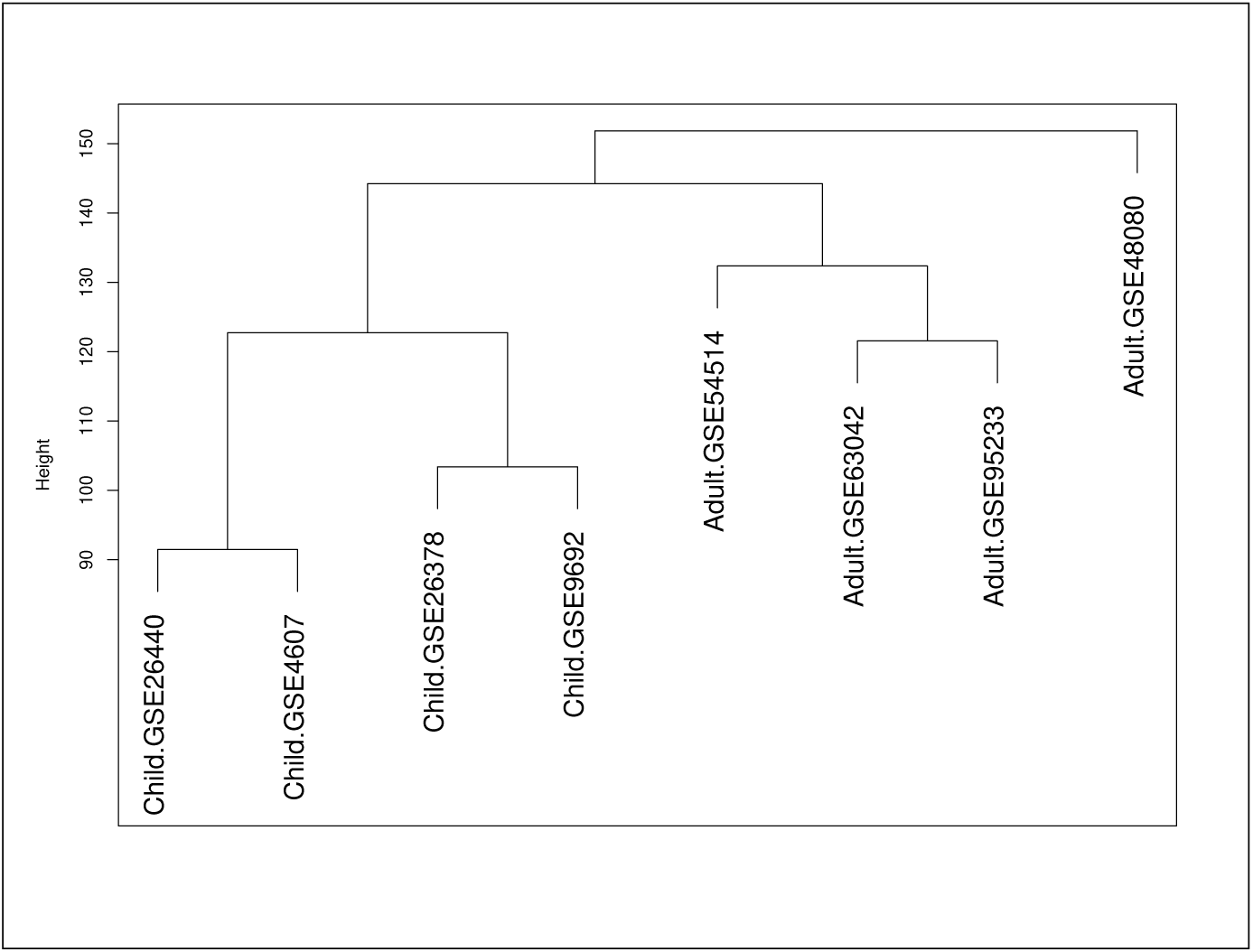
**Hierarchical Clustering:** Using expression of the common genes in the 8 data sets from GEO. The paediatric data (annotated with prefix “Child”) are clearly segregated from the “Adult” studies.

**Table S1.** Genomic change in available SS reports: This table contains genomic change in available reports in SS and our results of re-analysis of data downloaded from NCBI GEO. Details are available in Materials and Methods.

| Author_year | Data Source | Disease | Country | # Probes assayed | # DE probes | % change | FDR p threshold | Fold change threshold | Pubmed ID |
| --- | --- | --- | --- | --- | --- | --- | --- | --- | --- |
| Parnell_2013 | GSE54514 | Sepsis | Australia | 48804 | 3677 | 7.53 | 0.05 | NA | 23807251 |
| Severino_2014 | GSE48080 | Sepsis | Brazil | 19596 | 151 | 0.77 | 0.05 | 1.7 | 24667684 |
| Wong_2007 | GSE4607 | Septic shock | USA | 54675 | 2482 | 4.54 | 0.05 | NA | 17374846 |
| Wong_2009 | GSE13904 | Septic shock | USA | 54675 | 1867 | 3.41 | 0.001 | 2 | 17374846 |
| Wong_2010 | NA | Septic shock | USA | 54675 | 2355 | 4.31 | 0.05 | 1.5 | 20009785 |
| Calvano_2005 | GlueGrant | Endotoxin Shock | USA | 44000 | 5093 | 11.58 | 0.1 | NA | 16136080 |
| Xiao_2011 | GSE11375 | Critical Injury | USA | 47000 | 5136 | 10.93 | 0.001 | 2 | 22110166 |
| This manuscript | NA | Severe sepsis / septic shock | India | 17513 | 4221 | 24.10 | 0.05 | NA | NA |
| Re-analysis * | GSE4607 | Septic shock | USA | 22017 | 6179 | 28.1 % | 0.05 | NA | NA |
|  | GSE13904 | Septic shock | USA | 22017 | 7420 | 33.7 % | 0.05 | NA | NA |
|  | GSE26378 | Septic shock | USA | 22017 | 8143 | 37 % | 0.05 | NA | NA |
|  | GSE26440 | Septic shock | USA | 22017 | 10014 | 45.5 % | 0.05 | NA | NA |
|  | GSE8121 | Septic shock | USA | 22017 | 6517 | 29.6 % | 0.05 | NA | NA |
|  | GSE9692 | Septic shock | USA | 22024 | 12202 | 55.4 % | 0.05 | NA | NA |
|  | GSE95233 | Septic shock | France | 22017 | 11580 | 52.6 % | 0.05 | NA | NA |

**Table S2.** **Inclusion and exclusion criteria:** Clinical and laboratory features for recruitment of sepsis patients for the study

|  | Clinical features |
| --- | --- |
| **Inclusion criteria** | **Any two of the following clinical parameters is present in patient**<br>– Body temperature >38 °C or <36 °C<br>– Heart rate >90 beats per minute<br>– Respiratory rate >20 breaths/min or arterial PaCO<sub>2</sub> <32 mmHg<br>– WBC count >12 x 10<sup>9</sup> per microlitre or <4 x 10<sup>9</sup> per microlitre. |
| **Exclusion criteria** | Viral or fungal infection<br>Chronic inflammatory diseases such as asthma, rheumatoid arthritis etc.<br>Nosocomial (hospital acquired) infections |

**Table S3.**
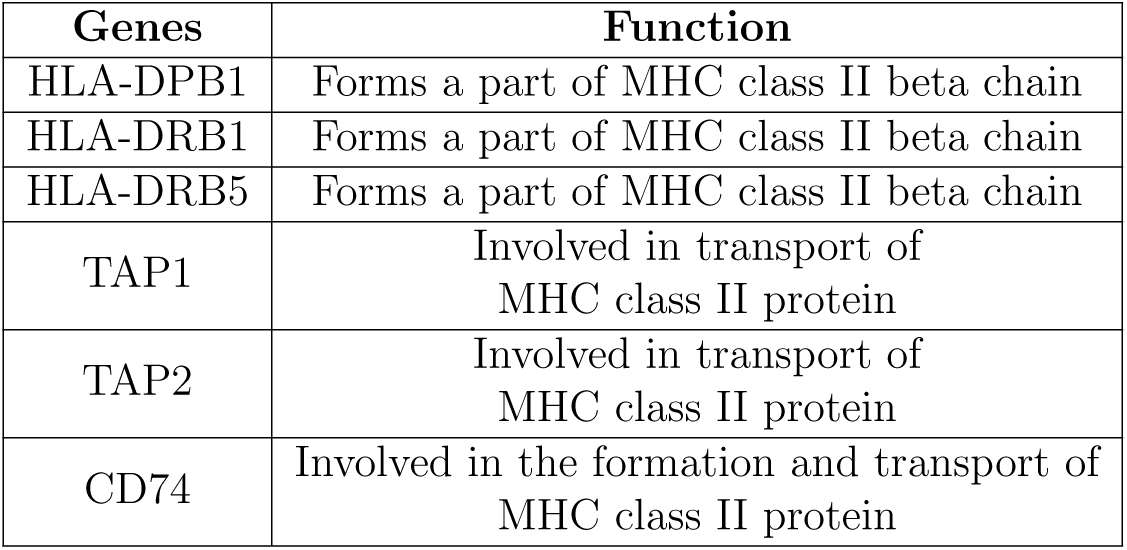
Annoatated table for MHCII genes with their function.

**Table S4.**
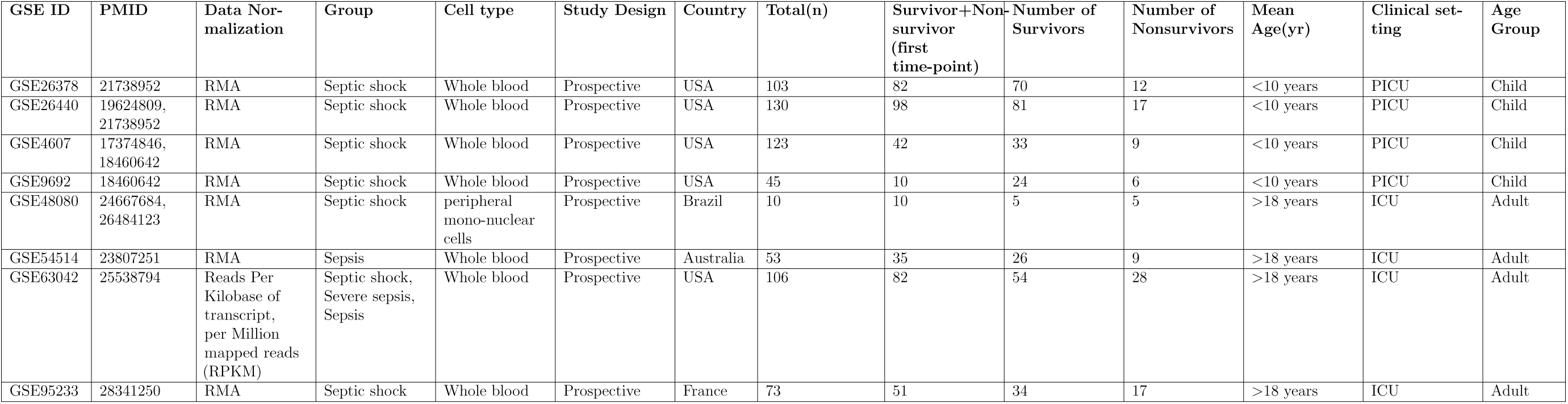
Study characteristics of all the datasets selected for survival meta-analysis.

| GSE ID | PMID | Data Normalization | Group | Cell type | Study Design | Country | Total(n) | Survivor+Non-survivor (first time-point) | Number of Survivors | Number of Nonsurvivors | Mean Age(yr) | Clinical setting | Age Group |
| --- | --- | --- | --- | --- | --- | --- | --- | --- | --- | --- | --- | --- | --- |
| GSE26378 | 21738952 | RMA | Septic shock | Whole blood | Prospective | USA | 103 | 82 | 70 | 12 | <10 years | PICU | Child |
| GSE26440 | 19624809, 21738952 | RMA | Septic shock | Whole blood | Prospective | USA | 130 | 98 | 81 | 17 | <10 years | PICU | Child |
| GSE4607 | 17374846, 18460642 | RMA | Septic shock | Whole blood | Prospective | USA | 123 | 42 | 33 | 9 | <10 years | PICU | Child |
| GSE9692 | 18460642 | RMA | Septic shock | Whole blood | Prospective | USA | 45 | 10 | 24 | 6 | <10 years | PICU | Child |
| GSE48080 | 24667684, 26484123 | RMA | Septic shock | peripheral mono-nuclear cells | Prospective | Brazil | 10 | 10 | 5 | 5 | >18 years | ICU | Adult |
| GSE54514 | 23807251 | RMA | Sepsis | Whole blood | Prospective | Australia | 53 | 35 | 26 | 9 | >18 years | ICU | Adult |
| GSE63042 | 25538794 | Reads Per Kilobase of transcript, per Million mapped reads (RPKM) | Septic shock, Severe sepsis, Sepsis | Whole blood | Prospective | USA | 106 | 82 | 54 | 28 | >18 years | ICU | Adult |
| GSE95233 | 28341250 | RMA | Septic shock | Whole blood | Prospective | France | 73 | 51 | 34 | 17 | >18 years | ICU | Adult |

## 1 General summary

Sepsis is one of the leading diseases associated with high mortality and morbidity, due to the systemic nature of this illness, transcriptomic technology is particularly suited to investigation of molecular underpinning of survival from sepsis episodes. We adopted an analysis approach that combined published transcriptome data and data generated in our laboratory from Indian sepsis patients leading to the discovery of key immune pathways to be altered in non-survivors compared to survivors. This is the first clinical transcriptomic study on sepsis from India, showing that non-survival is associated with down-regulated adaptive immune pathways and significant M2-specific immune-suppression, possibly regulated by NF-κB signalling. Three Biological processes related to sepsis were observed to be significantly altered in non-survivors. A patient-specific analysis reveals up-regulation of coagulation and inflammation but a strong down-regulation of immunosuppression modules in Indian sepsis patients.

**Figure 1:**
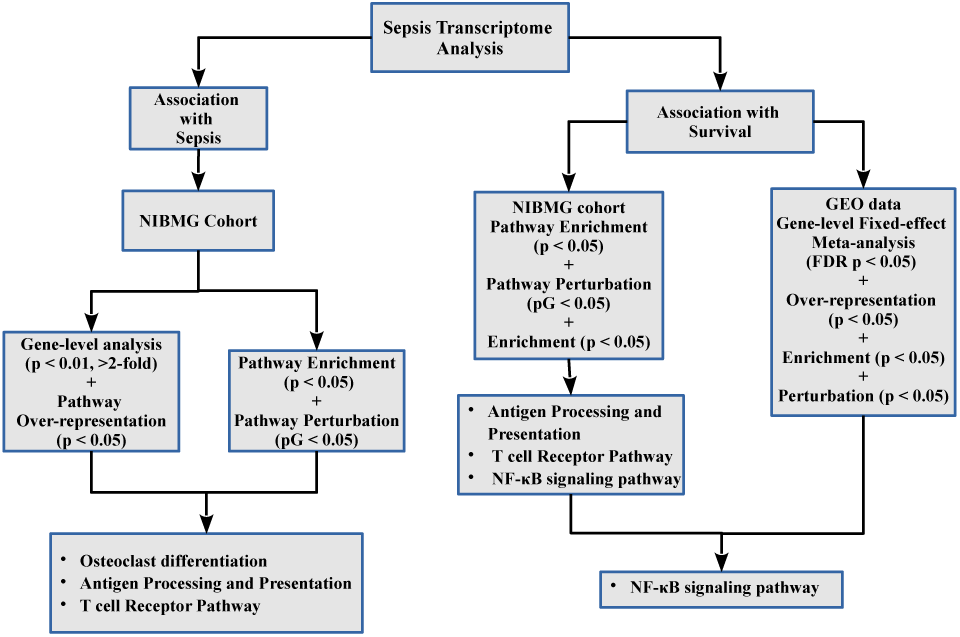
Analysis flow with results: The left arm describes the steps for case-vs-control analysis whereas the right arm describes the steps for identification of pathways associated with survival

## 2 Getting the data

Availability of data and materials Data and R code are available at the following link (https://figshare.com/) (search for the project **ssnibmgsurv**). The data are accessed through the two data packages listed below. The code can be downloaded as a single zip file (**ssnibmgsurvdoc.zip**). Upon uncomperssing the zip, install the two data packages as described below and run the subsequent code to generate the appropriate output.

### 2.1 Installation of the data package ssnibmgsurv

Dowload the file **ssnibmgsurv_1.0.tar.gz** from https://figshare.com/ (search for **ssnibmgsurv**). Change the directory to where you saved the file. Start R. At the R prompt, issue the following command:

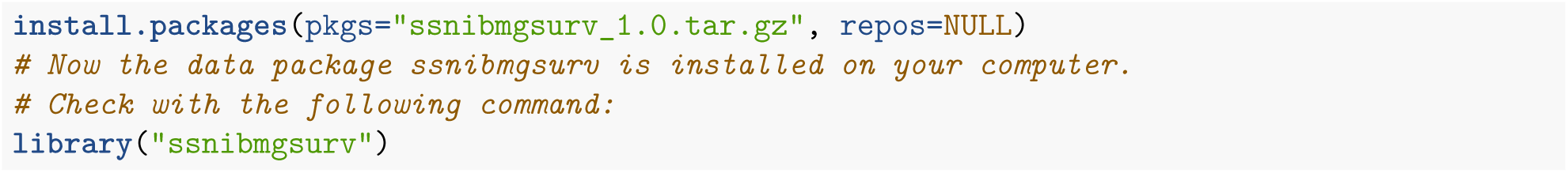

### 2.2 Installation of the data package ssgeosurv

Dowload the file **ssgeosurv_1.0.tar.gz** from https://figshare.com/ (search for **ssgeosurv**). Change the directory to where you saved the file. Start R. At the R prompt, issue the following command:

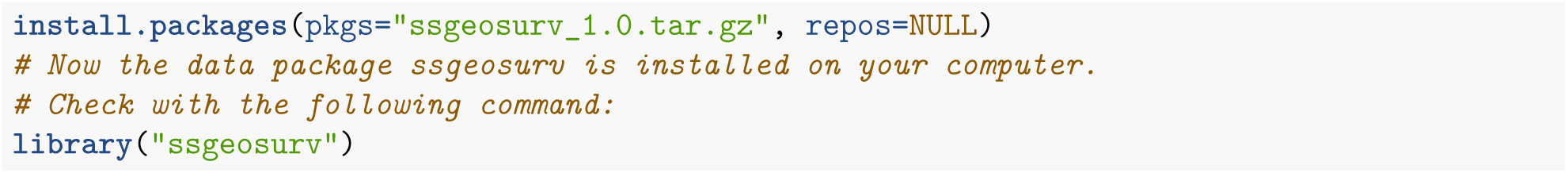

Now both the data packages are installed in your computer; let’s focus on starting the analysis.

### 2.3 Running the analysis: Setting working directory

It is assumed that you have access to a folder **ssnibmgsurvdoc**. Start R and set the working directory to **ssnibmgsurvdoc**. Run the code chunks as follow.

## 3 Case control analysis: Identifying genes and pathways DE in sepsis

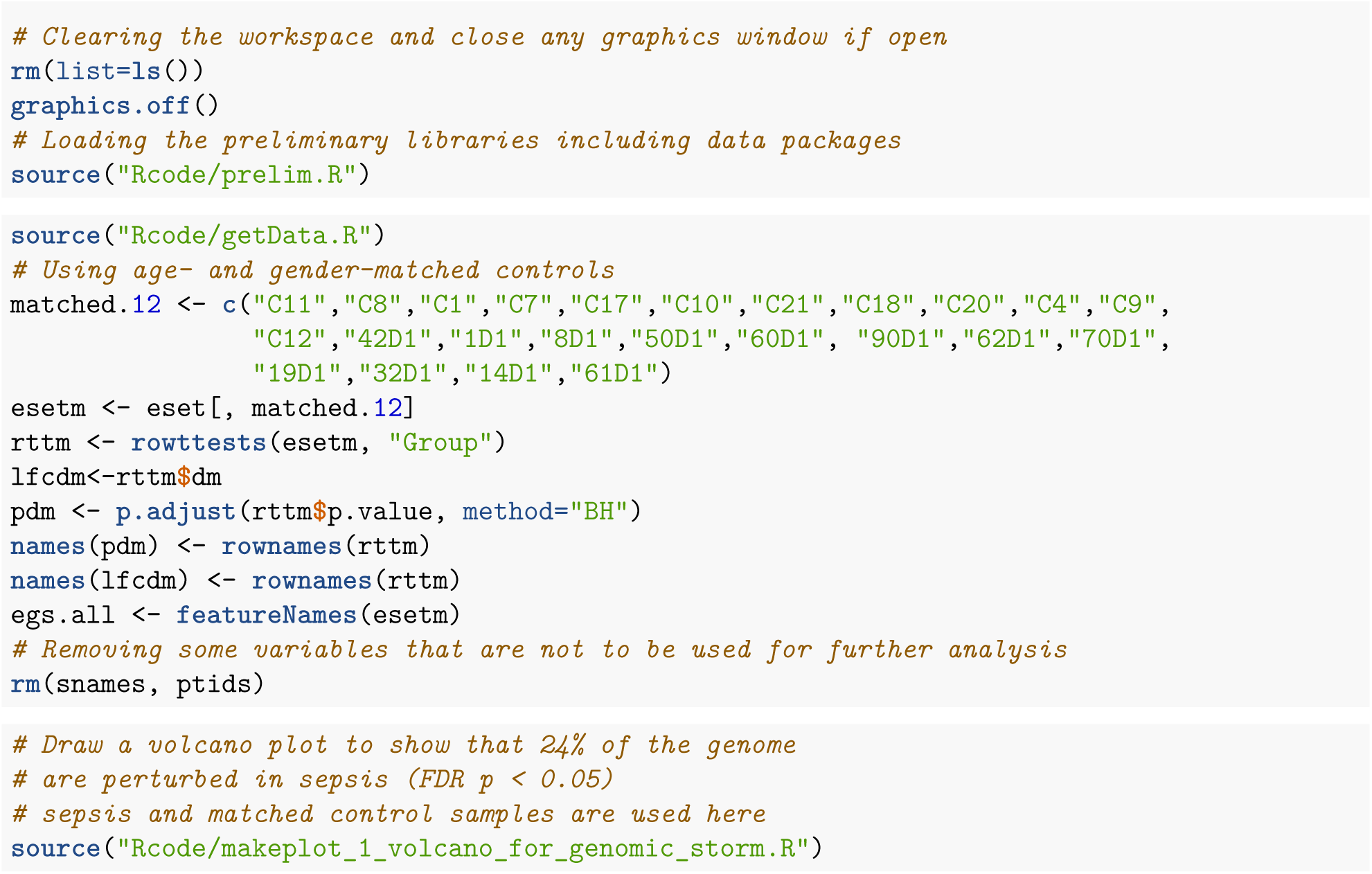

**Figure 2:**
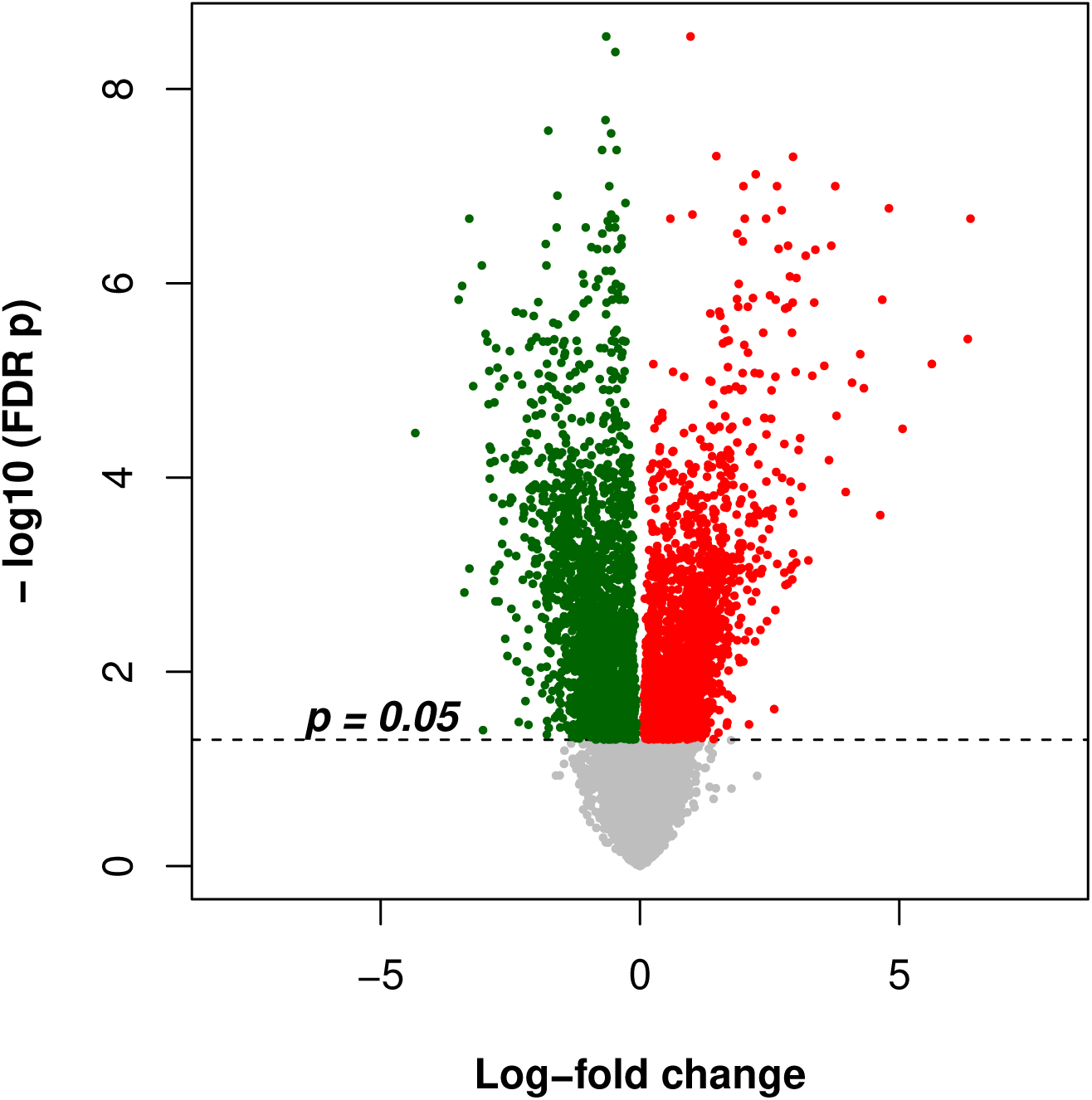
Volcano plot showing 24 percent of the genome perturbed in sepsis compared to healthy control (FDR p < 0.05). This establishes large scale change in gene expression in sepsis, and possible multiple pathways being perturbed.

### 3.1 Genome-level changes in gene expression

Differential gene expression analysis revealed 1109 genes to be altered in sepsis patients compared to age- and gender-matched healthy controls. Volcano plot (Figure 2) showed 24% of the genome perturbed in sepsis compared to healthy control (FDR p < 0.05). This establishes large scale change in gene expression in sepsis, and possible multiple pathways being perturbed.

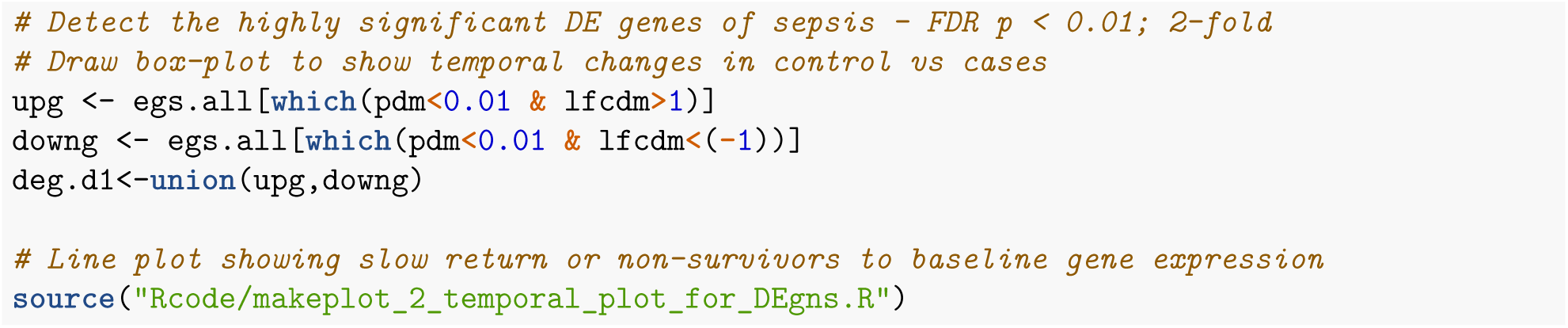

**Figure 3:**
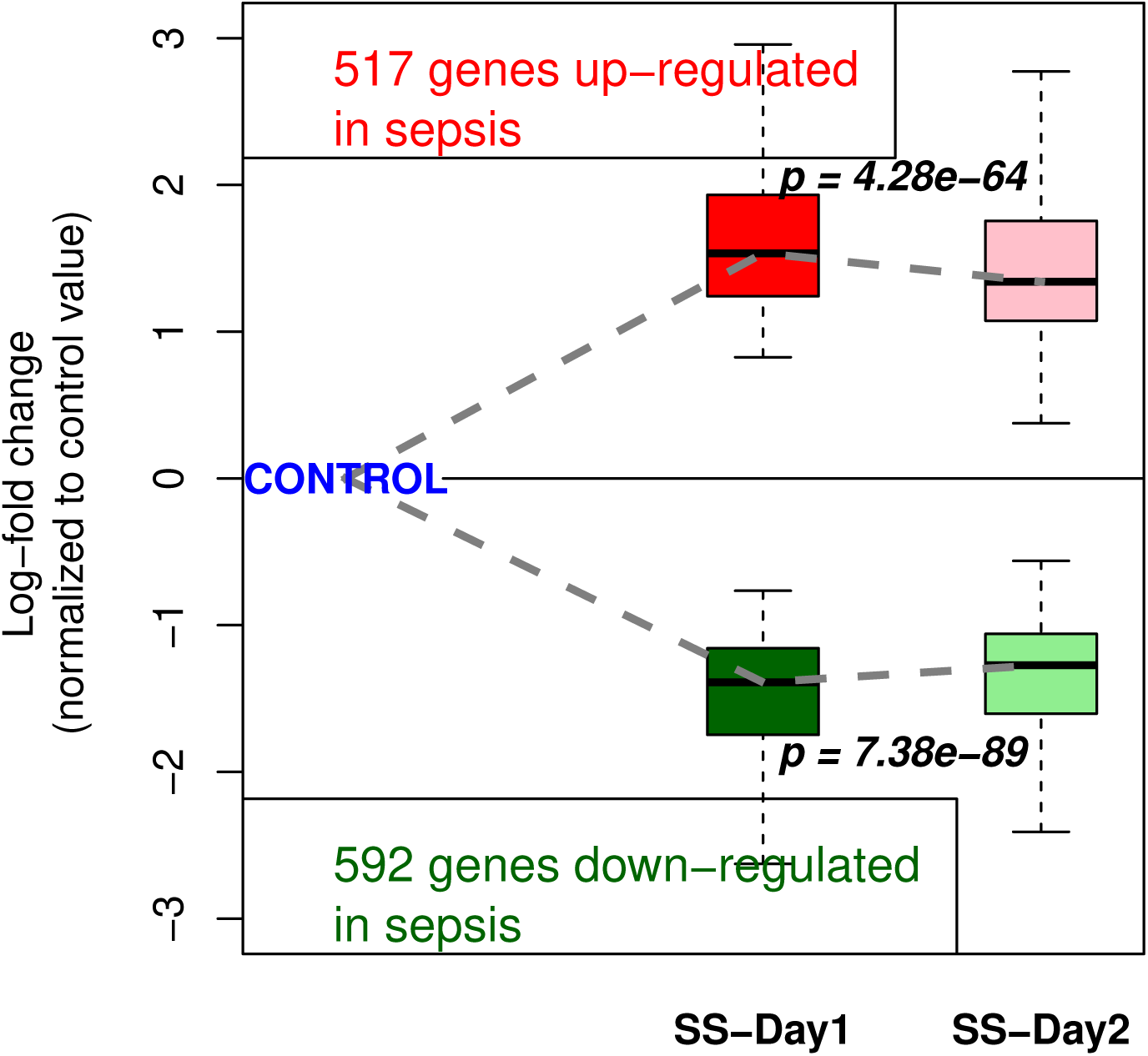
Temporal change of DE genes (FDR p < 0.05, 2 fold-change or more), there is a non-random trend toward the baseline with time (p-values from paired t-tests are provided in the legend). This is consistent with earlier findings from patients with trauma.

**Figure 4:**
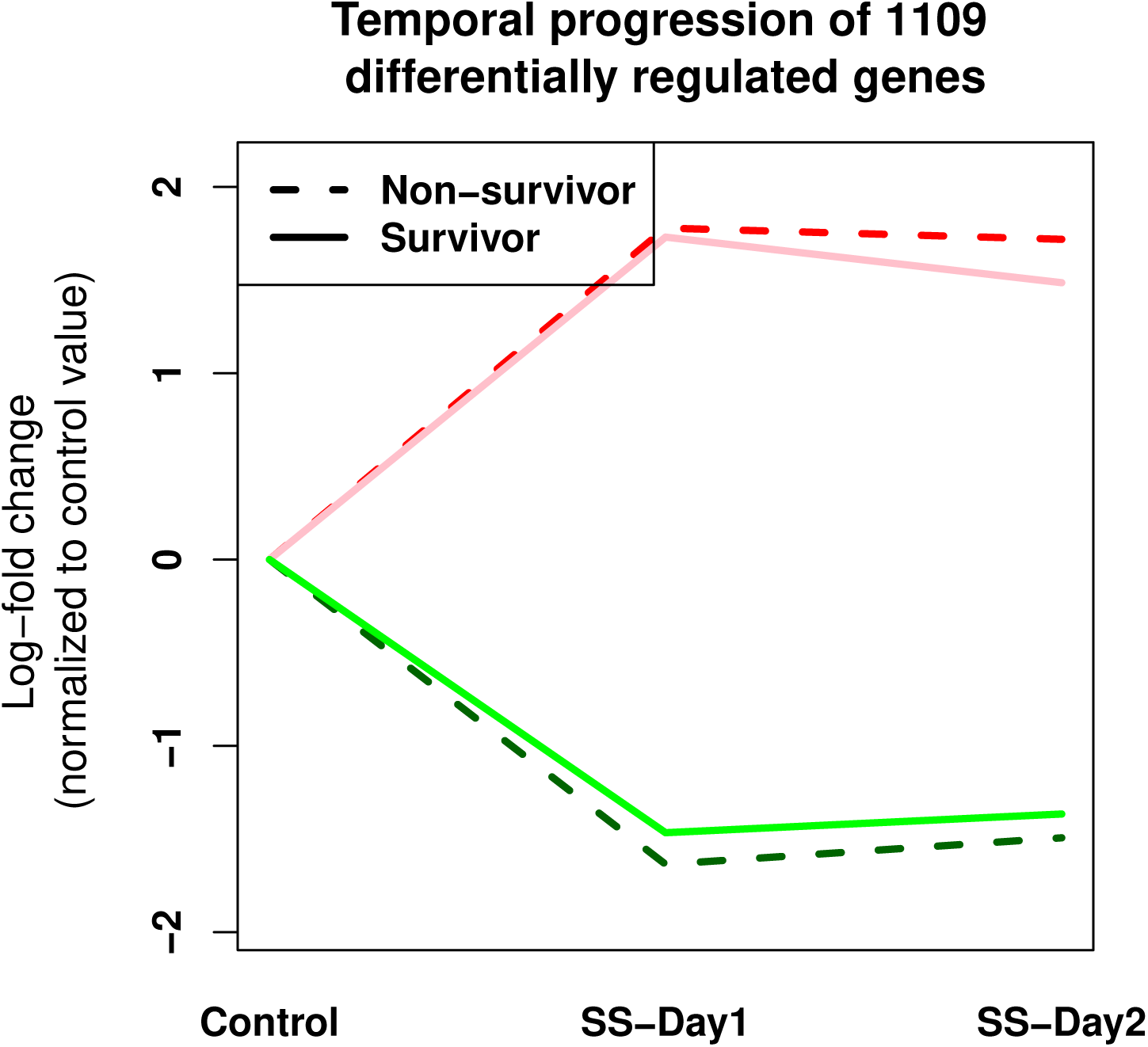
Temporal change of DE genes (FDR p < 0.05, 2 fold-change or more), there is a non-random trend toward the baseline with time (p-values from paired t-tests are provided in the legend). The delayed return to baseline is associated with non-recovery from sepsis.

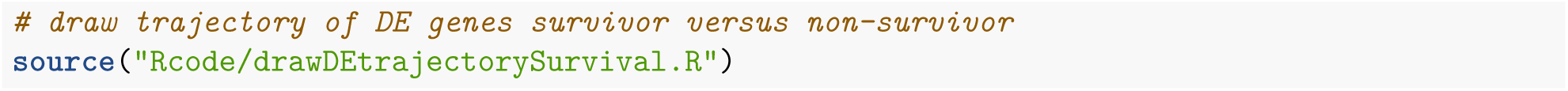

### 3.2 Pathway analysis results

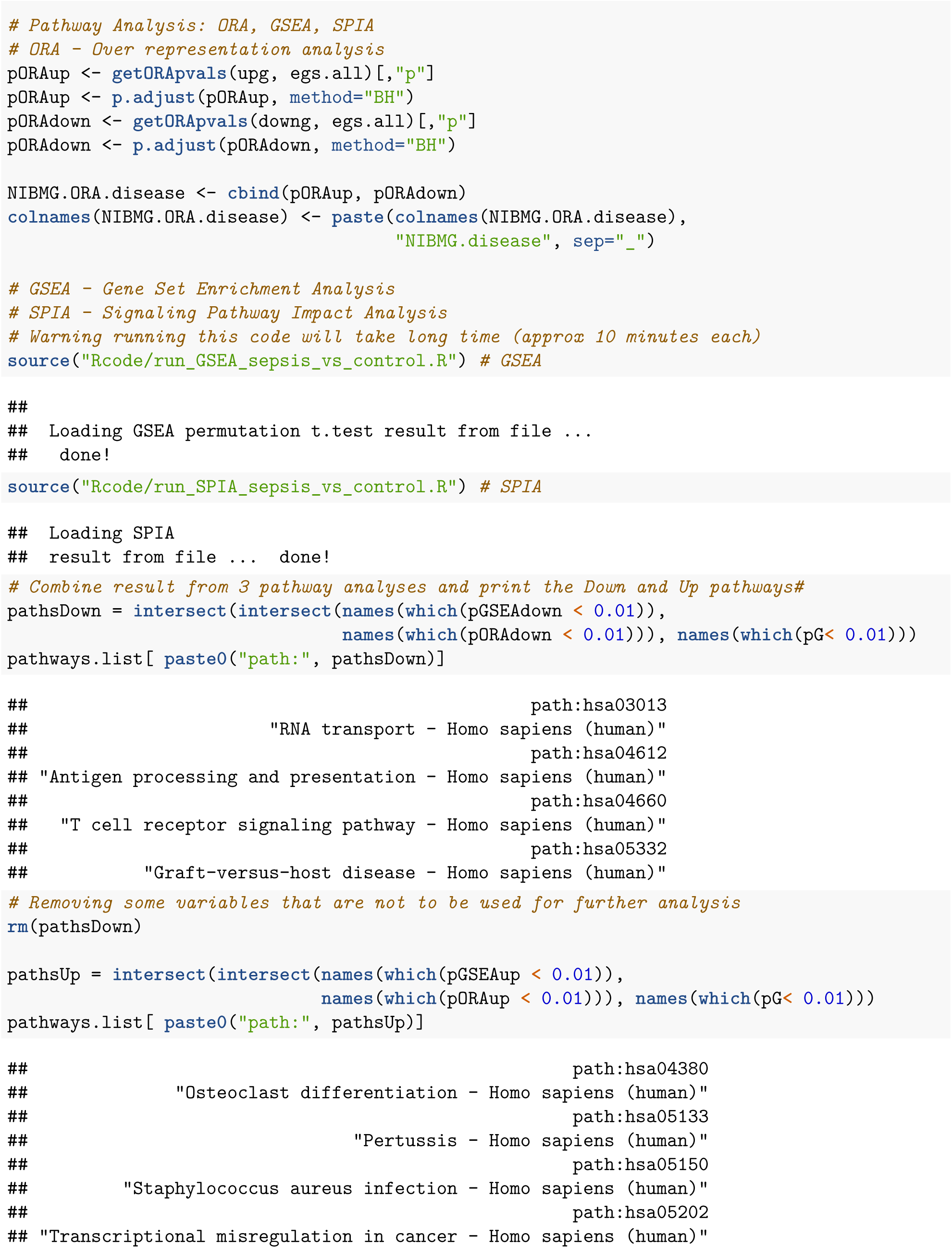

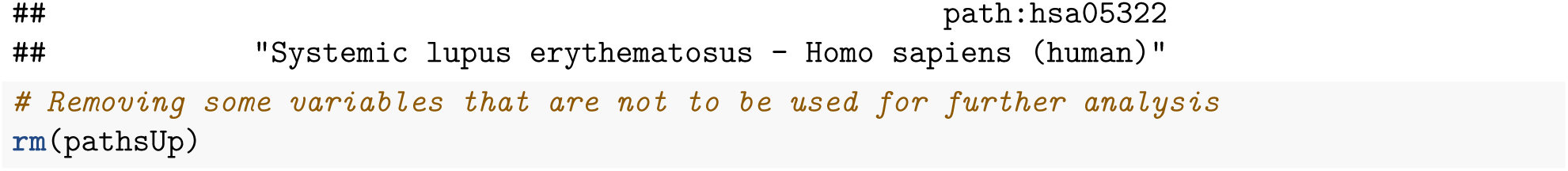

**Figure 5:**
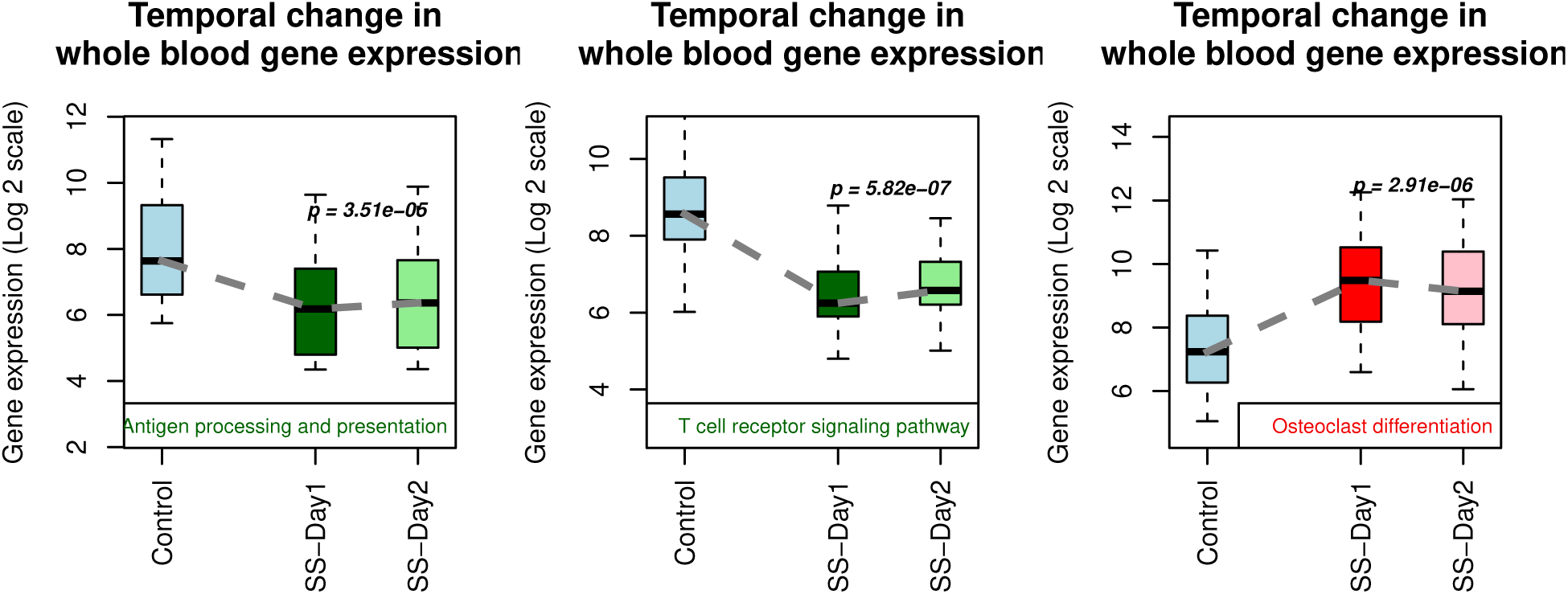
Temporal boxplot of 3 key pathwayes altered in Sepsis by combined pathway analysis.

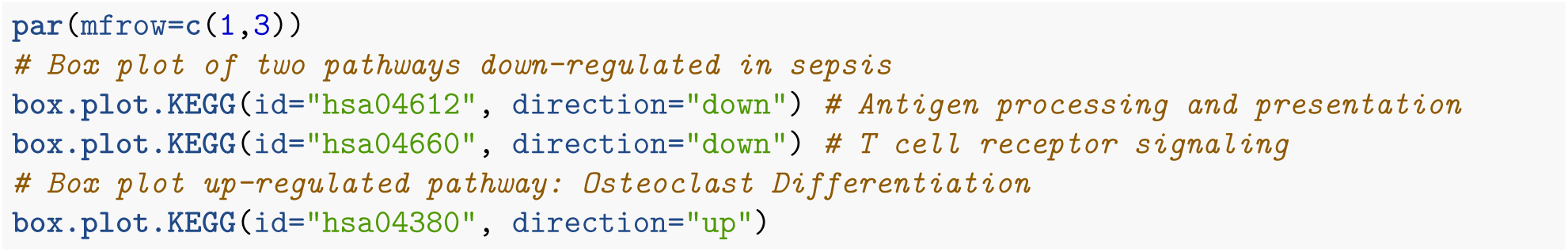

## 4 Survival analysis

### 4.1 Survival analysis using published transcriptome data (from NCBI GEO)

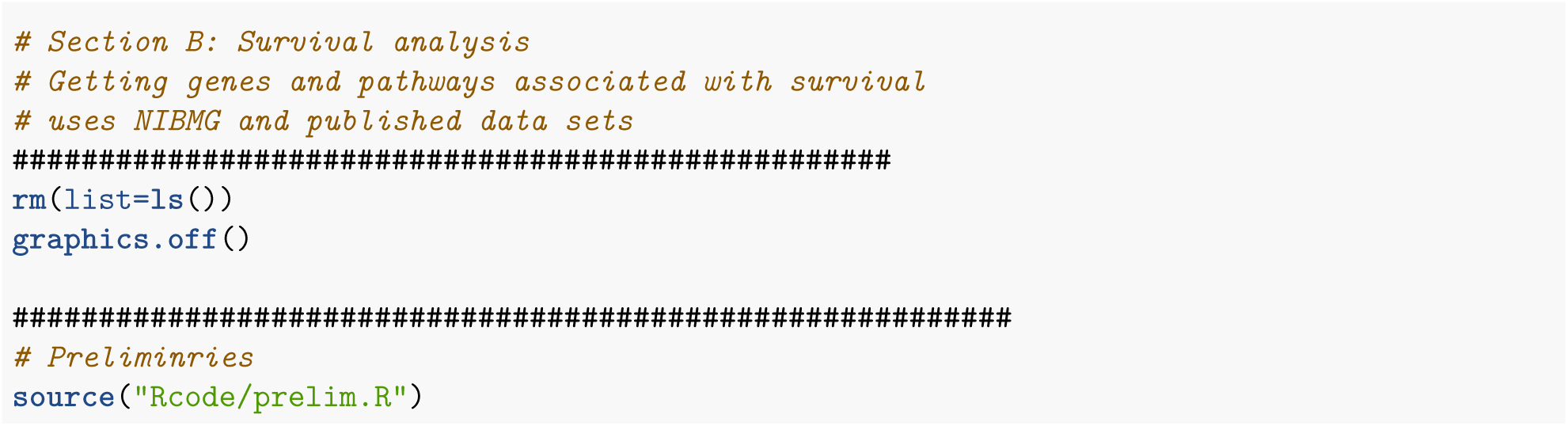

**Figure 6:**
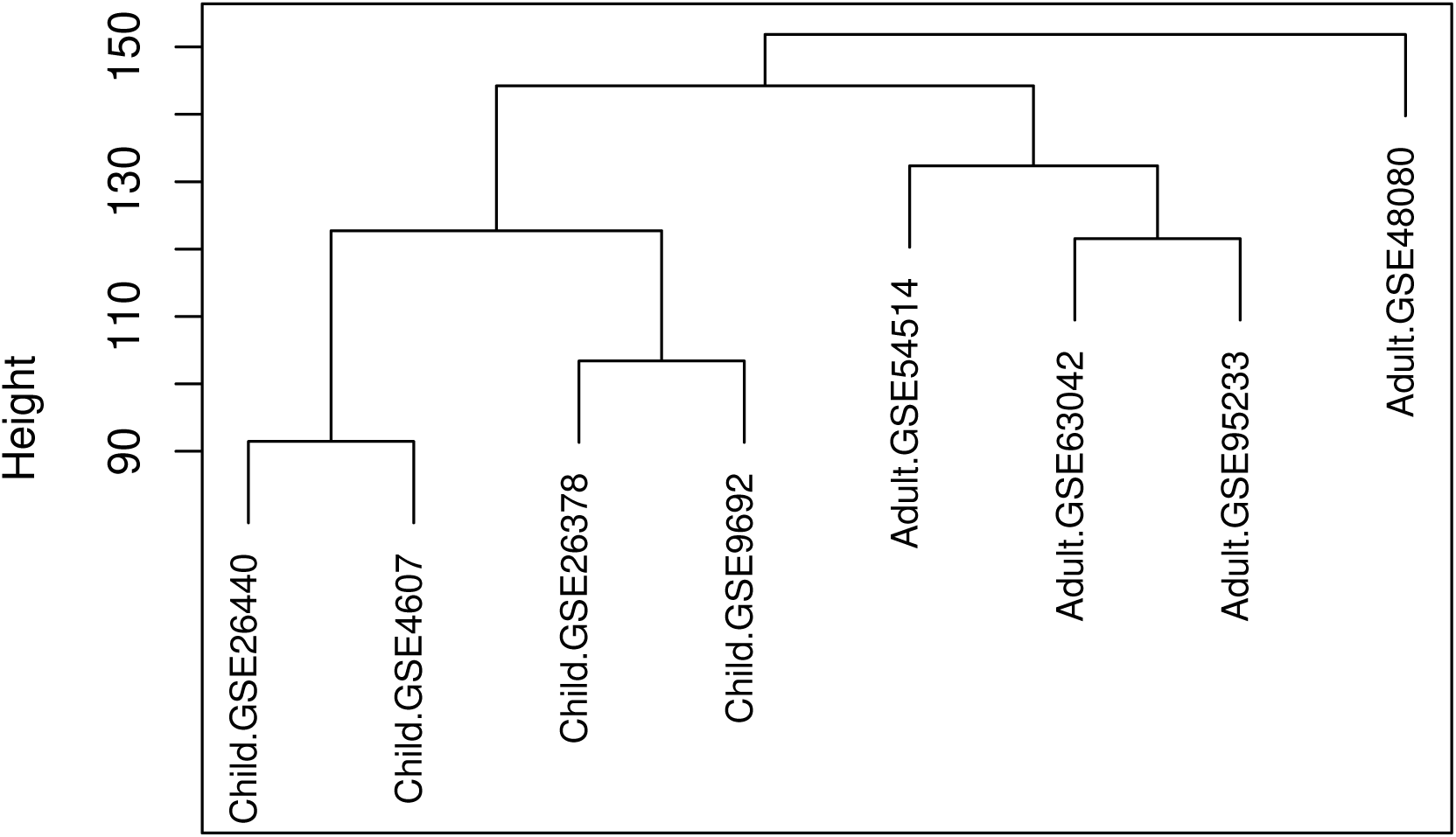
Survival analysis with eight published data sets: human adults and children with sepsis; hierarchical clustering with log-fold change in gene expression led to evidence of developmental age-specific differential perturbation; i.e., separate clusters for adult and child data sets. In view of this difference, further analysis was confined to to Adult data sets when combined with NIBMG data.

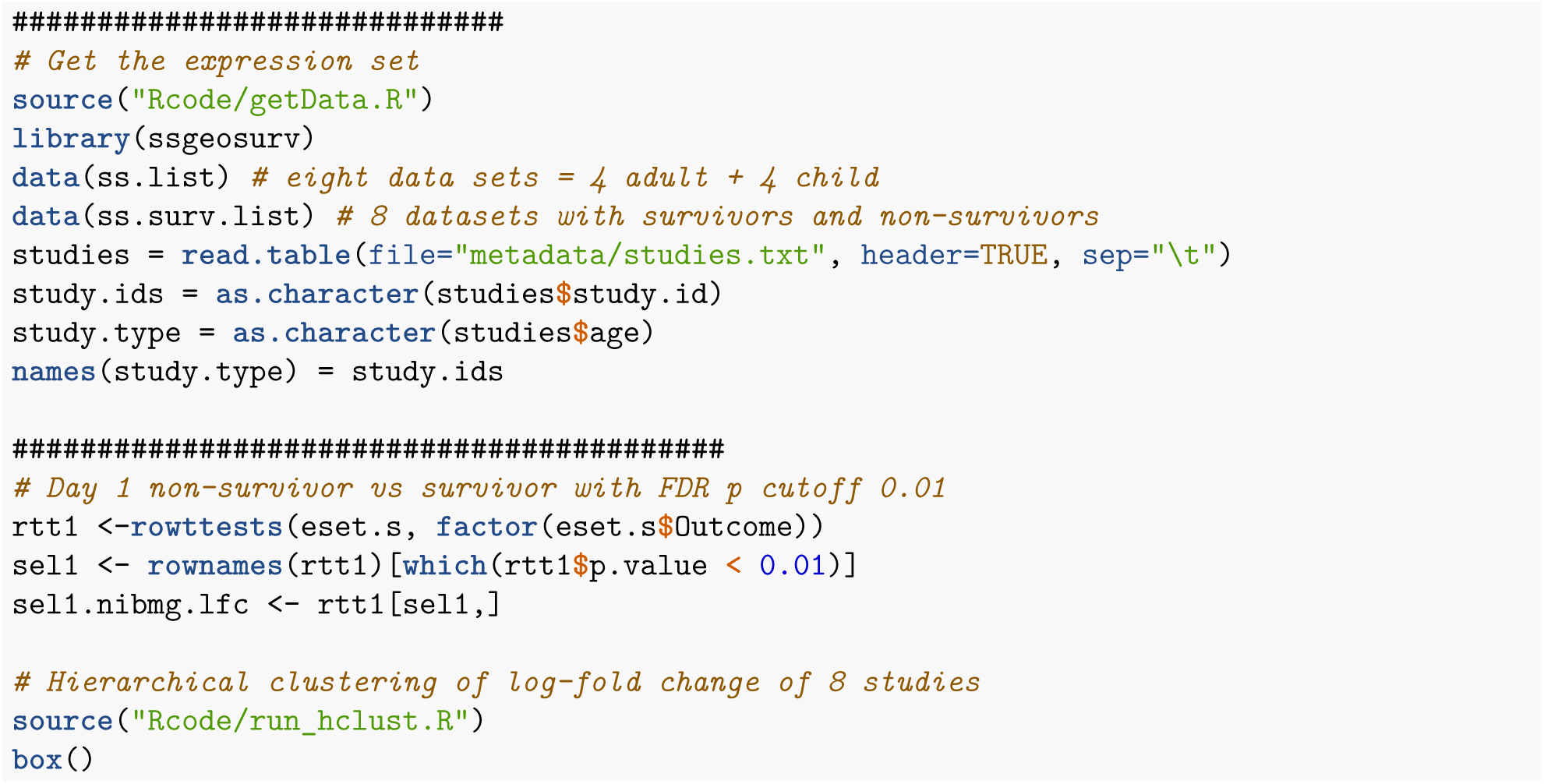

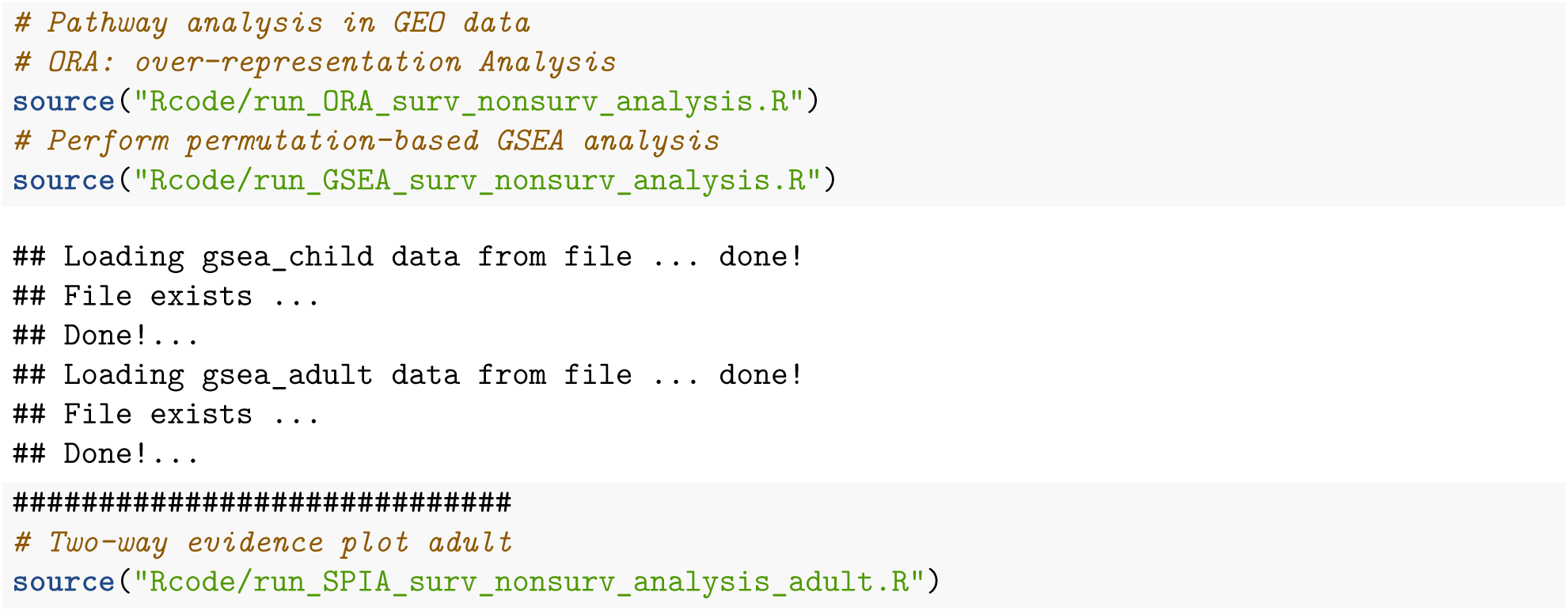

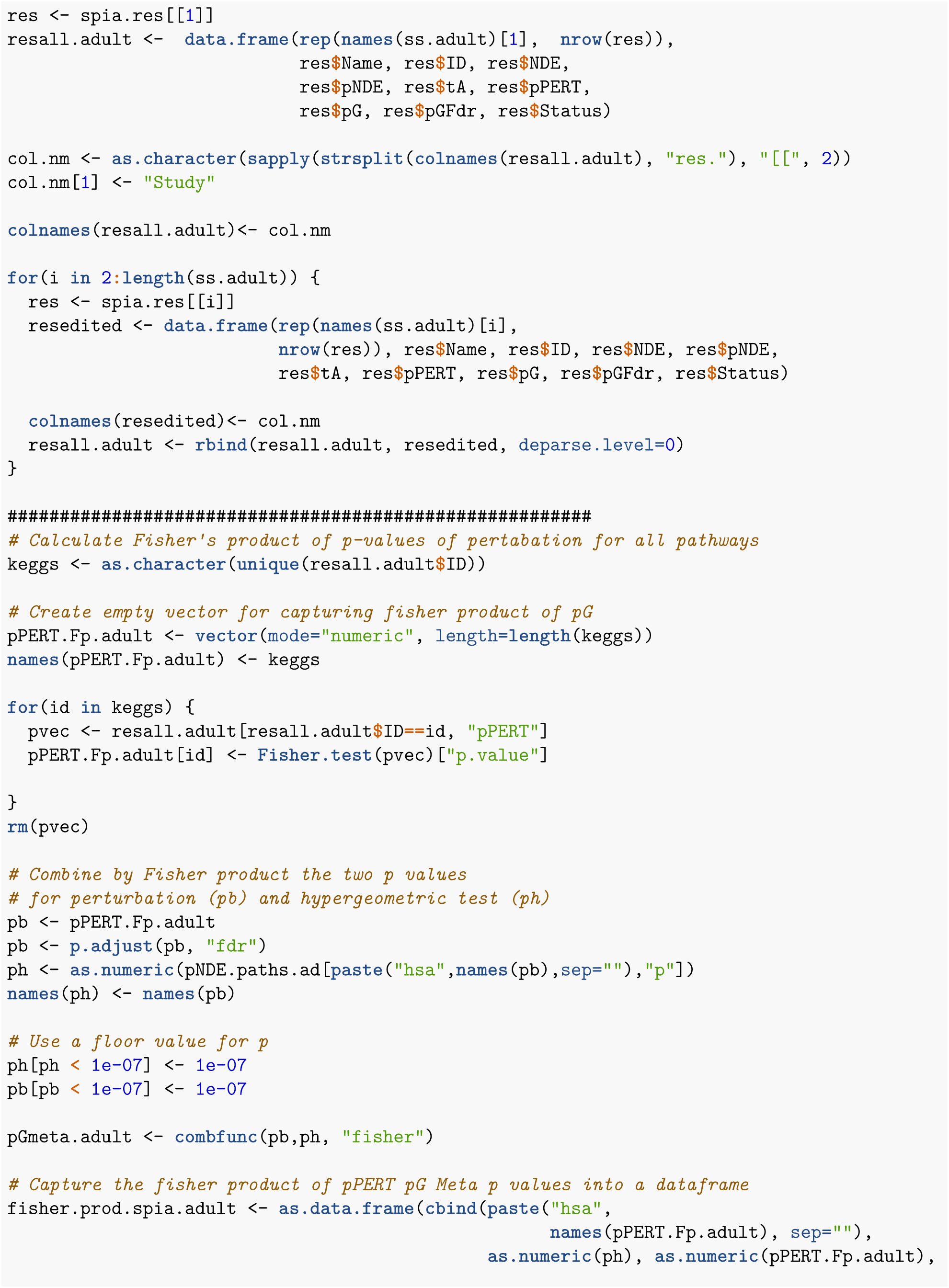

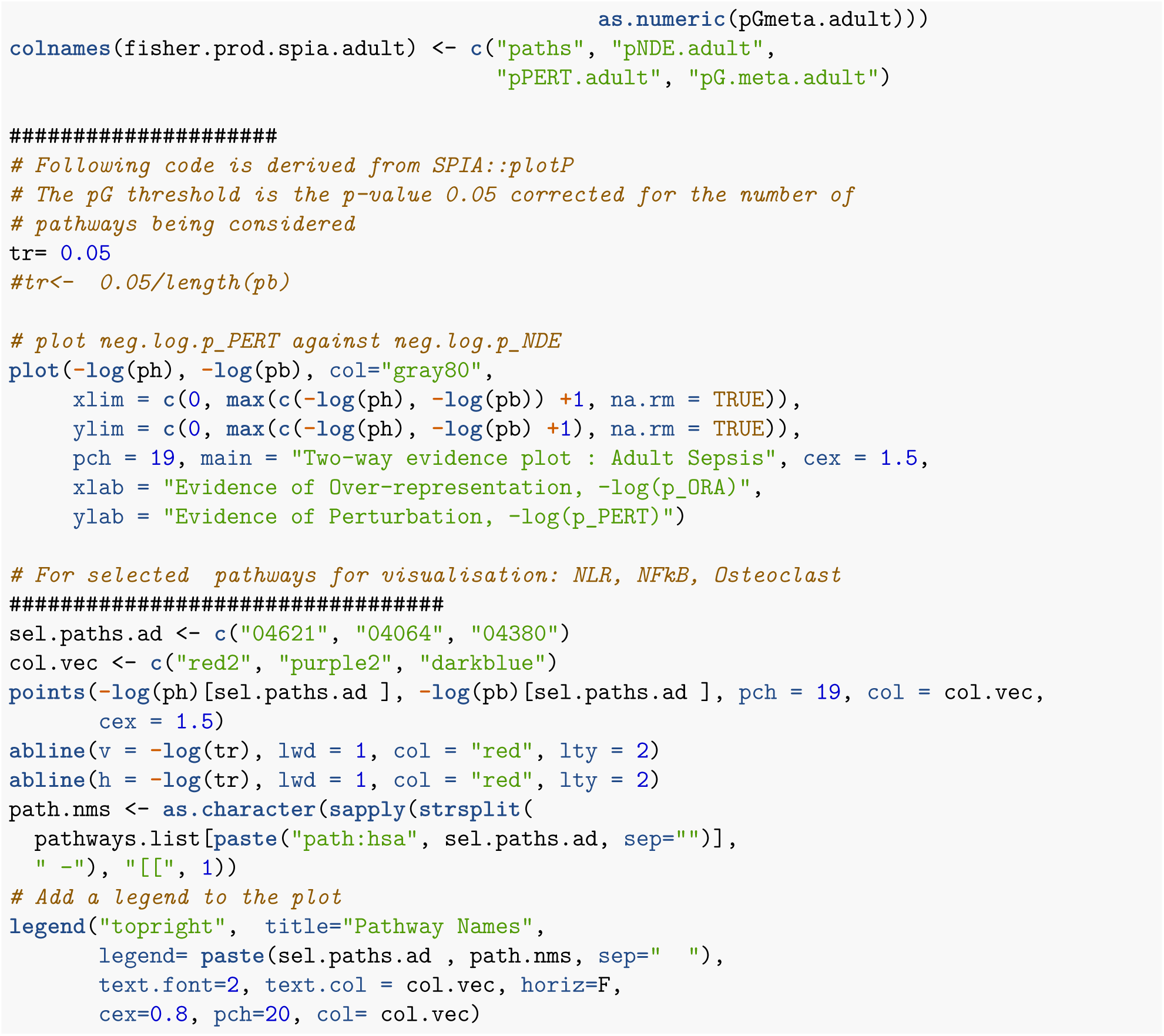

**Figure 7:**
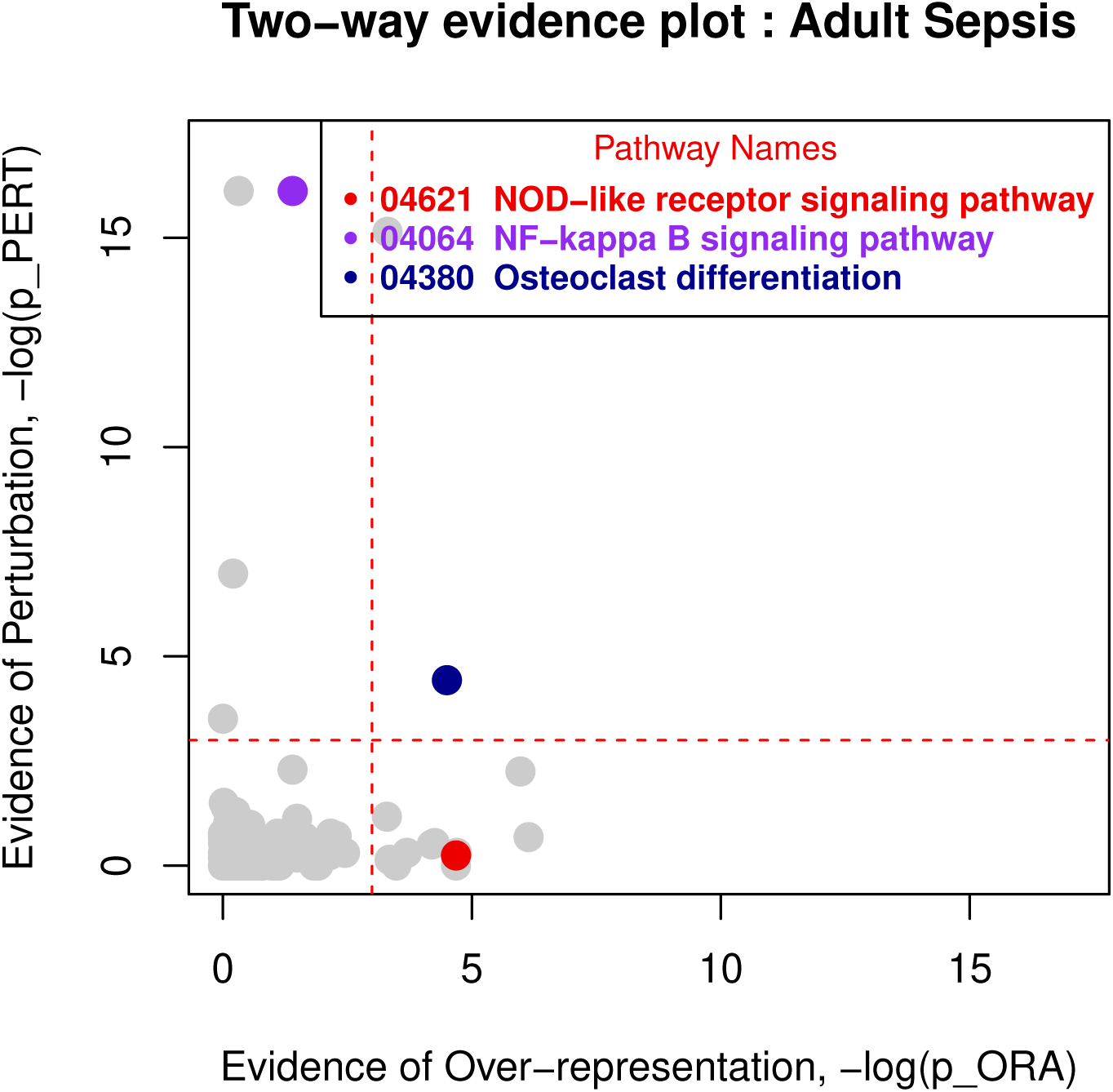
KEGG pathways associated with survival in GEO data; NF-kappaB signalling pathway, Osteoclast differentiation and NOD-like receptor signalling pathway.

**Figure 8:**
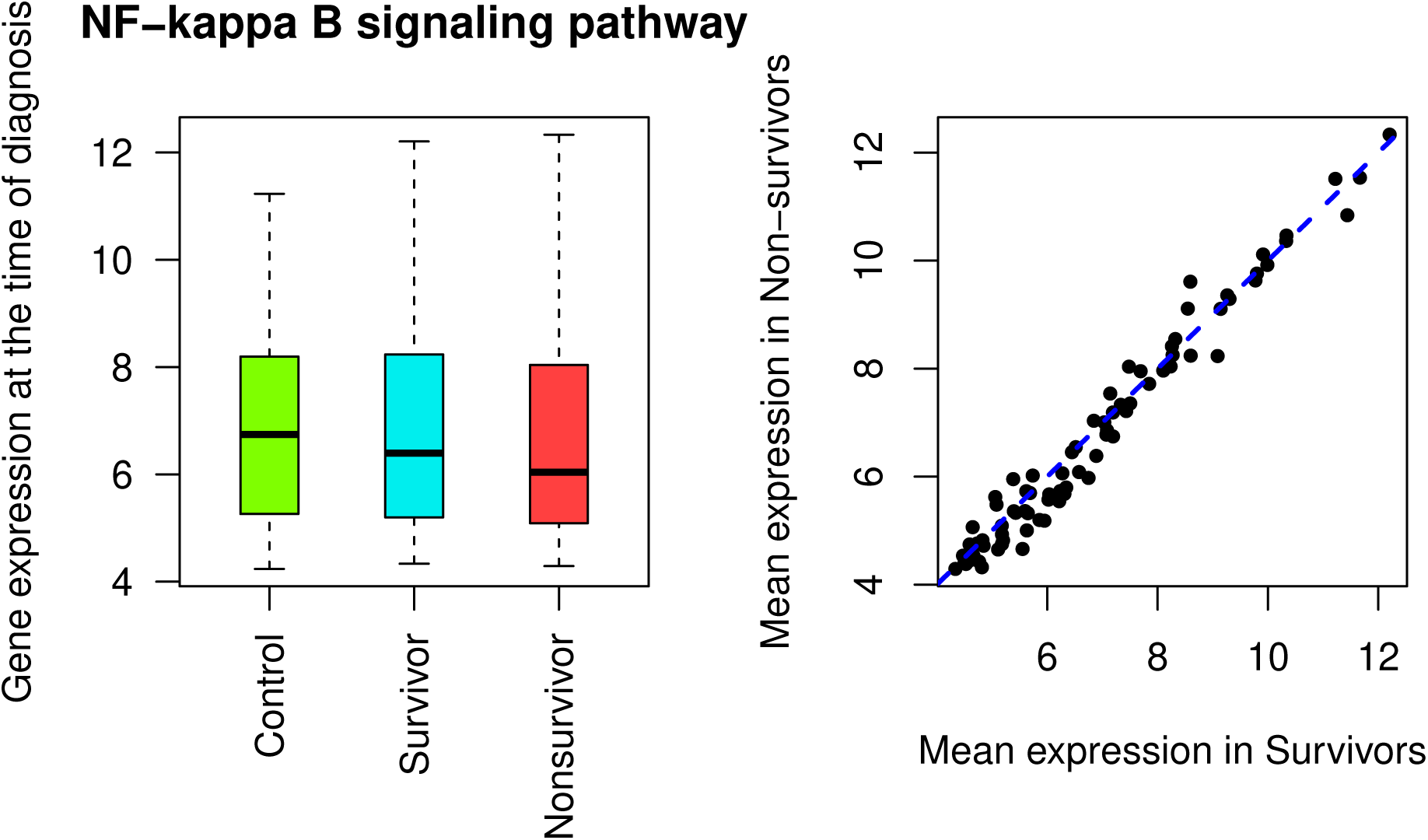
NF-kappa B signalling pathway boxplot and scatterplot

### 4.2 Down-regulation of NF-**κ** B signalling pathway genes in non-survivors

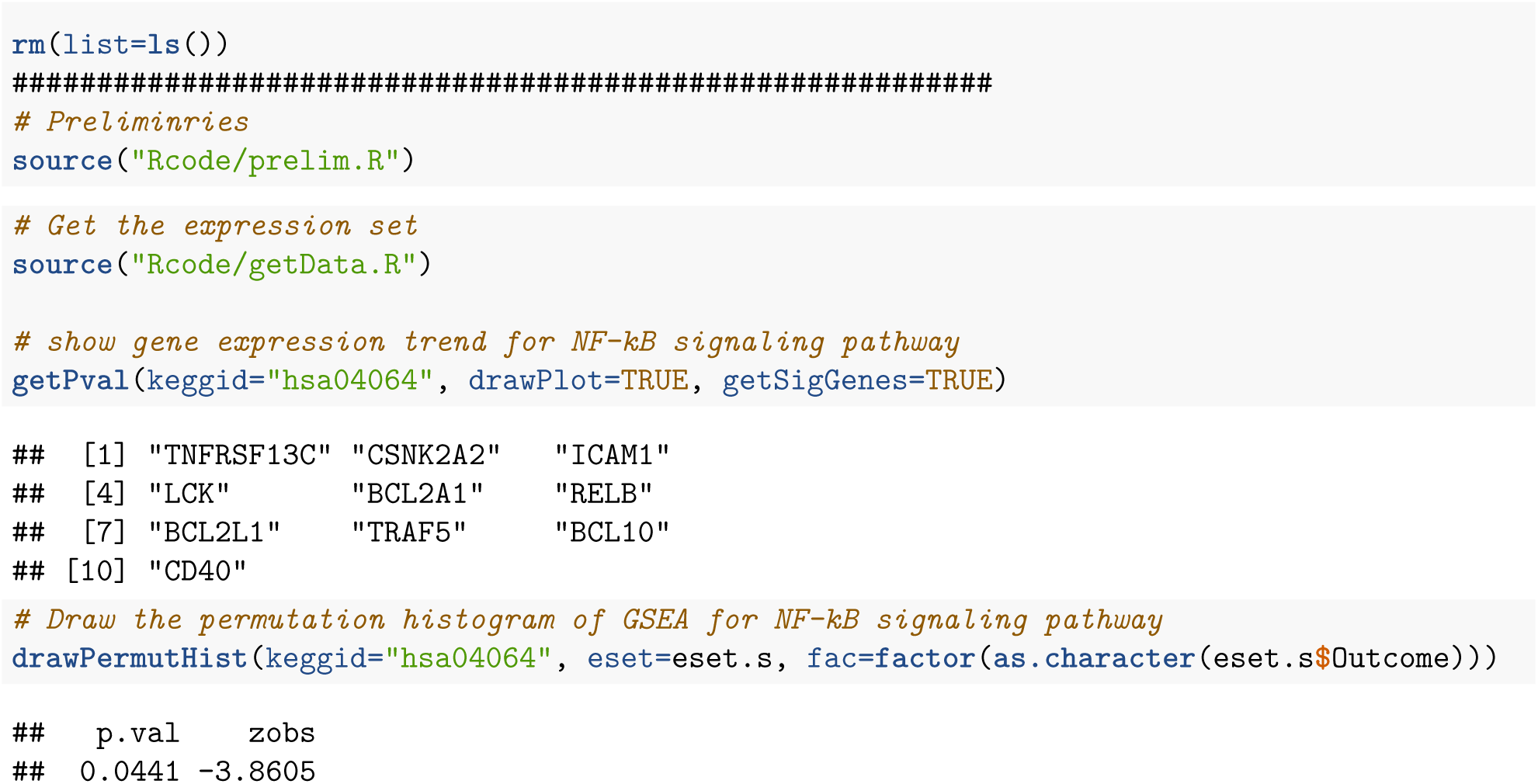

**Figure 9:**
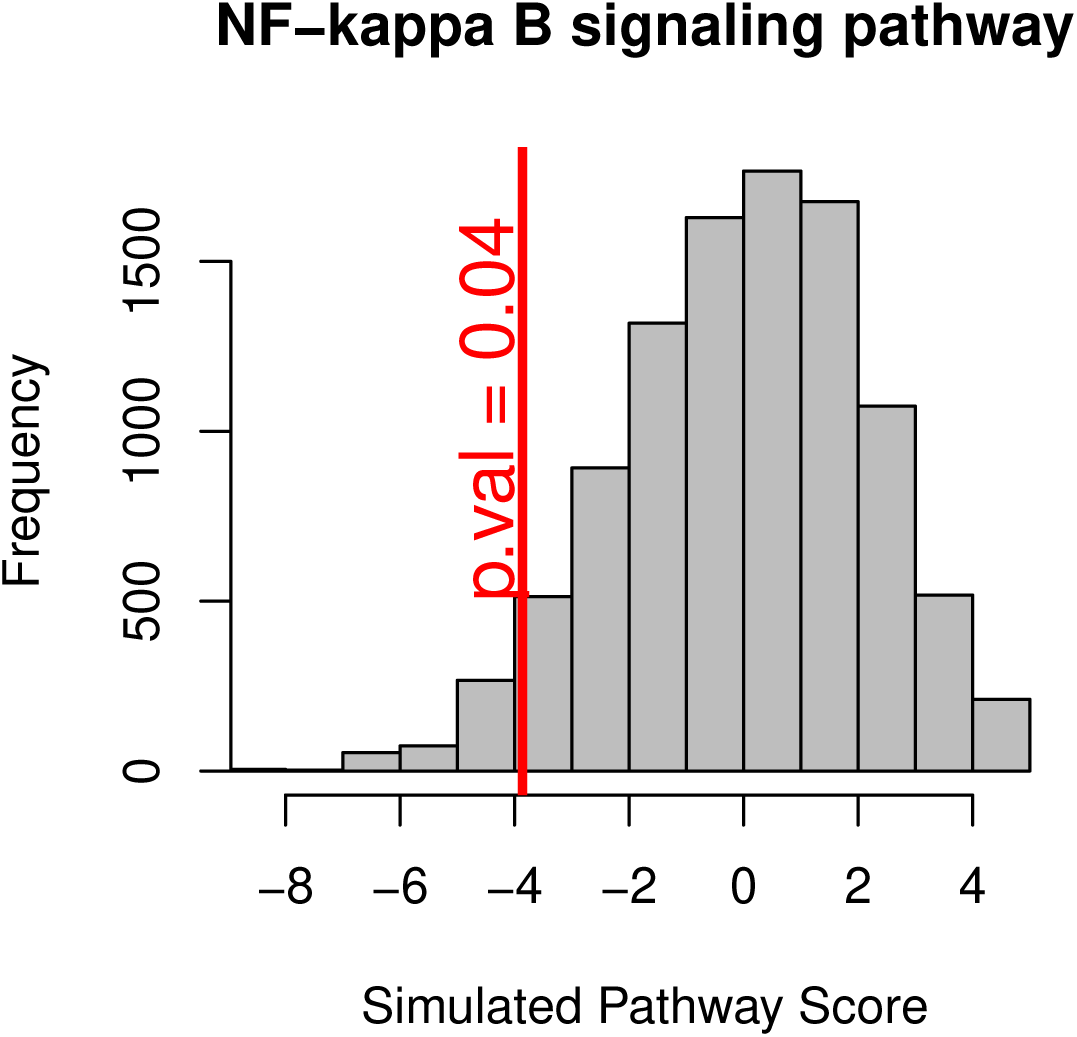
NF-kappa B signalling pathway histogram. Permutation bases Gene set enrichment analysis creats a histogram of simulated pathway scores (in gray bars). The red line shows the observed pathway score in nonsurvivors when compared to survivor.

### 4.3 Relative gene expression of the targets of NF-**κ**B

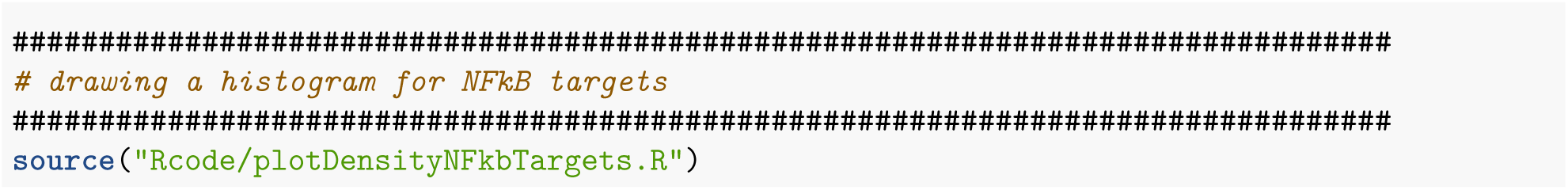

**Figure 10:**
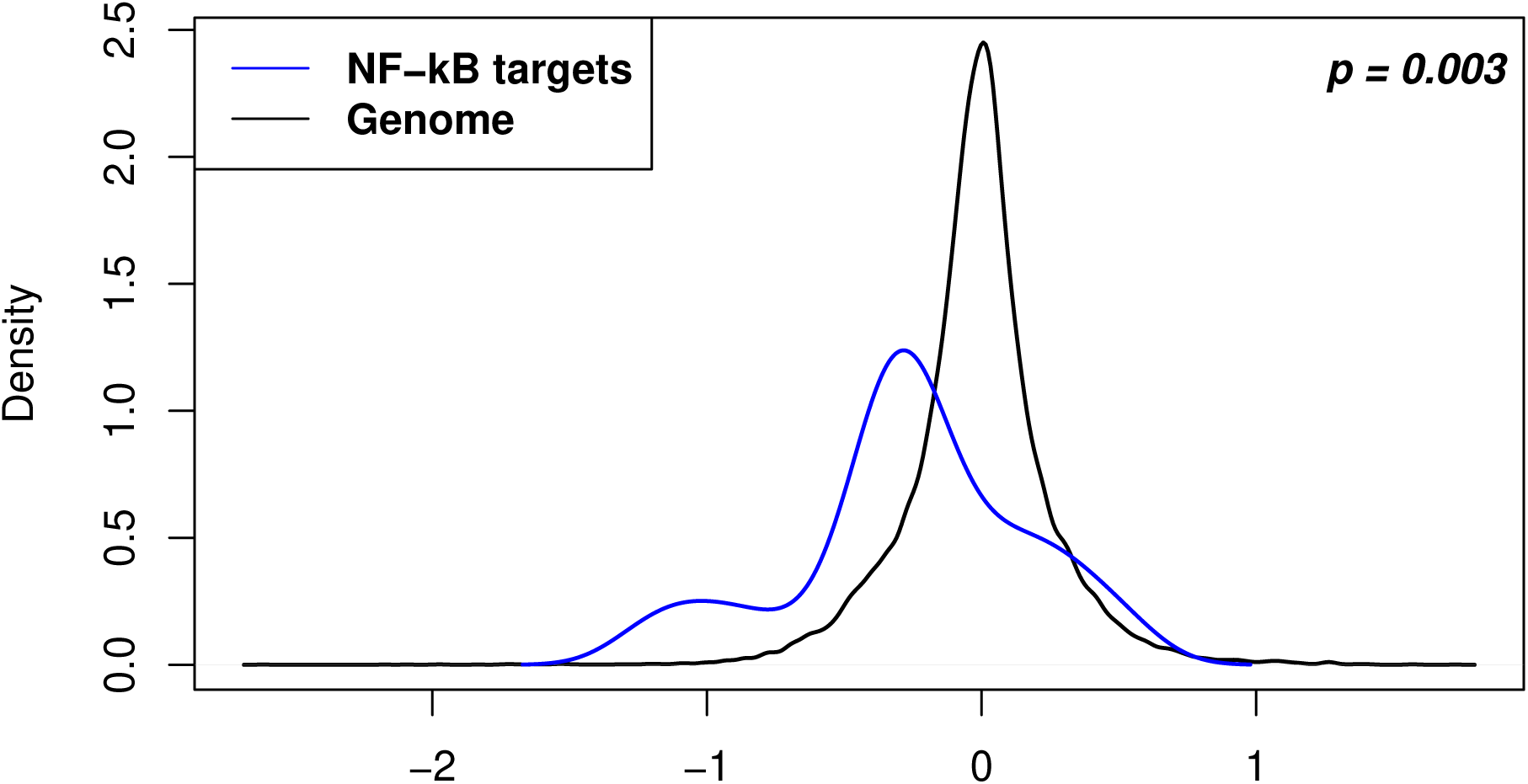
Relative gene expression of the targets of NFkB (i.e., Antigen processing and presentation genes and Immune receptor Genes). The gray peak in the background representsdistribution of all genes in the genome. There is significant down-regulation of the targets in the non-survivors (blue line).

### 4.4 M2 macrophage-specific down-regulation of gene expression in non-survivors

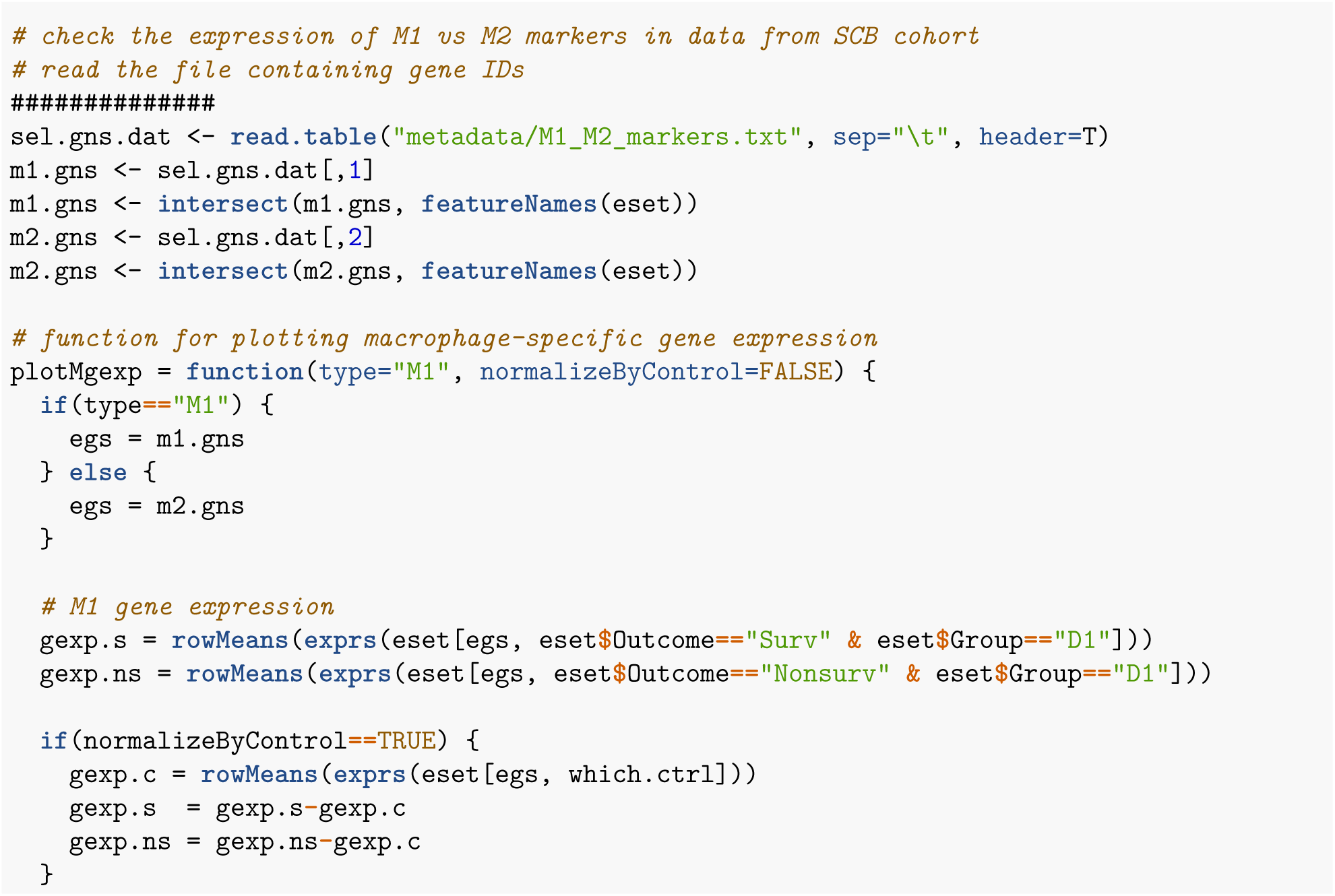

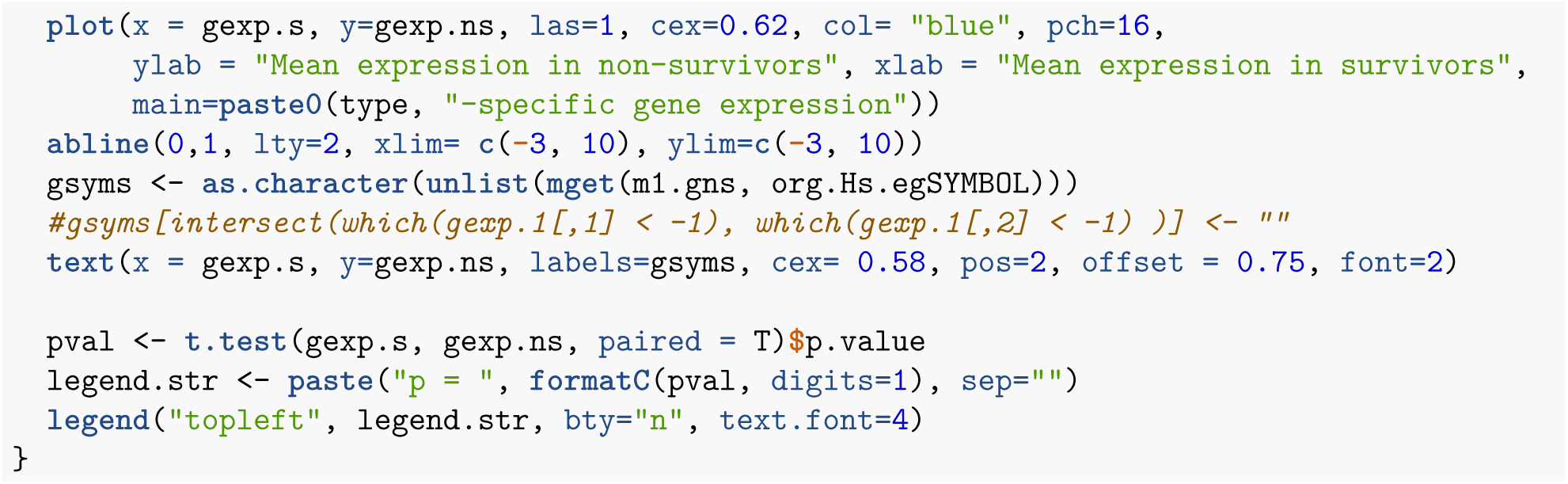

**Figure 11:**
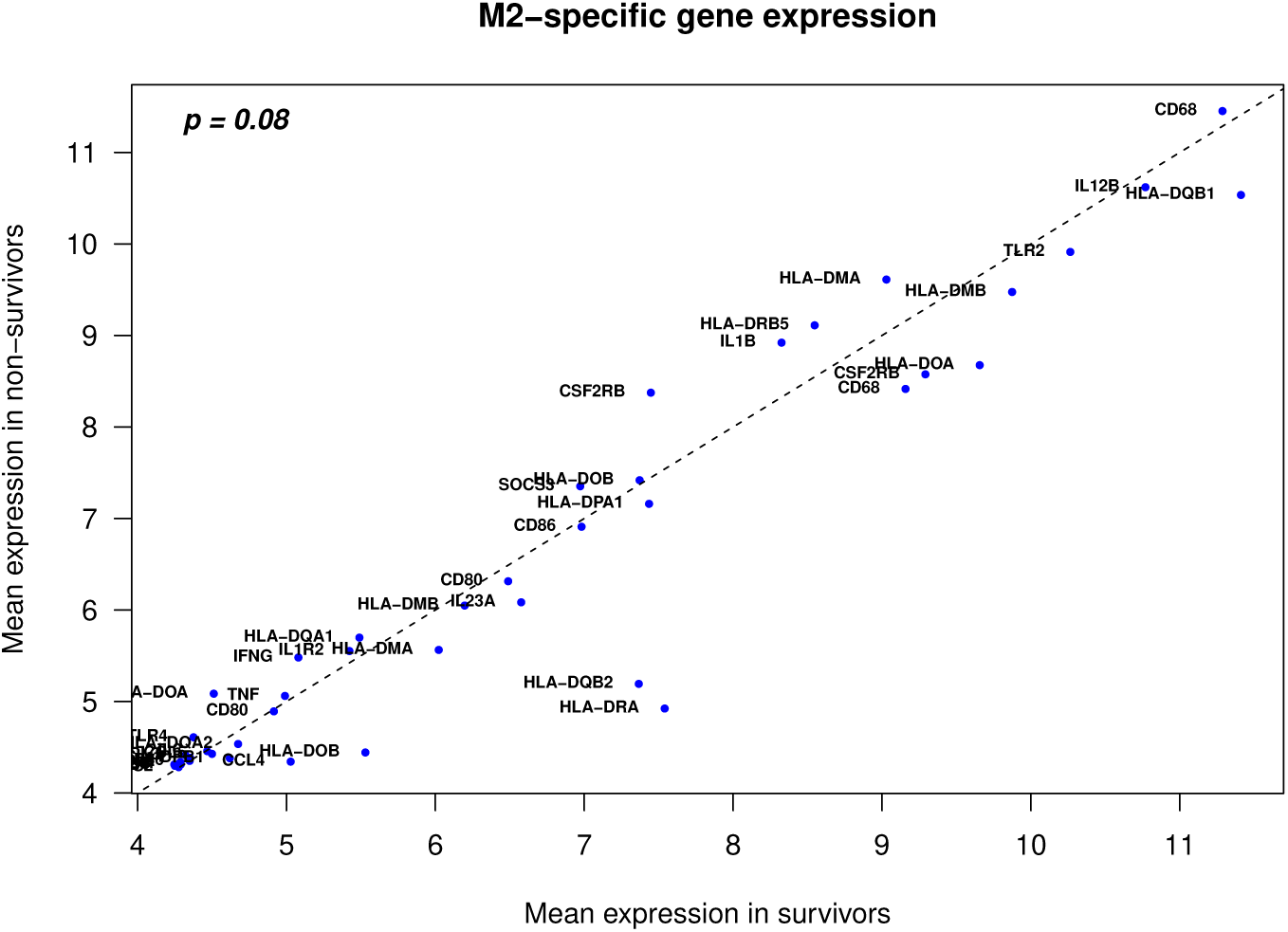
M1 macrophages (classically activated macrophages) are pro-inflammatory, important in host defence against the pathogens, phagocytosis, secretion of pro-inflammatory cytokines and microbicidal molecules. M2 macrophages (alternatively activated macrophages) participate in regulation of resolution of inflammation and repair of damaged tissues. M2-specific under-expression is observed in non-survivors (p = 0.02).

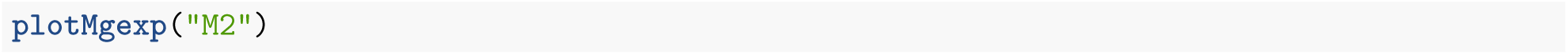

### 4.5 Down-regulation of Antigen processing and presentation signalling pathway genes in non-survivors

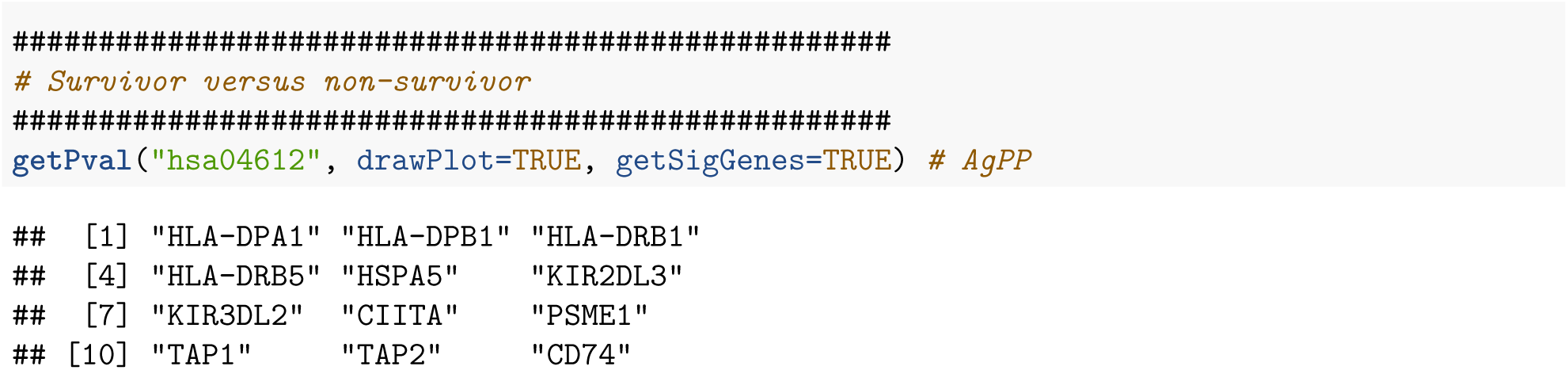

**Figure 12:**
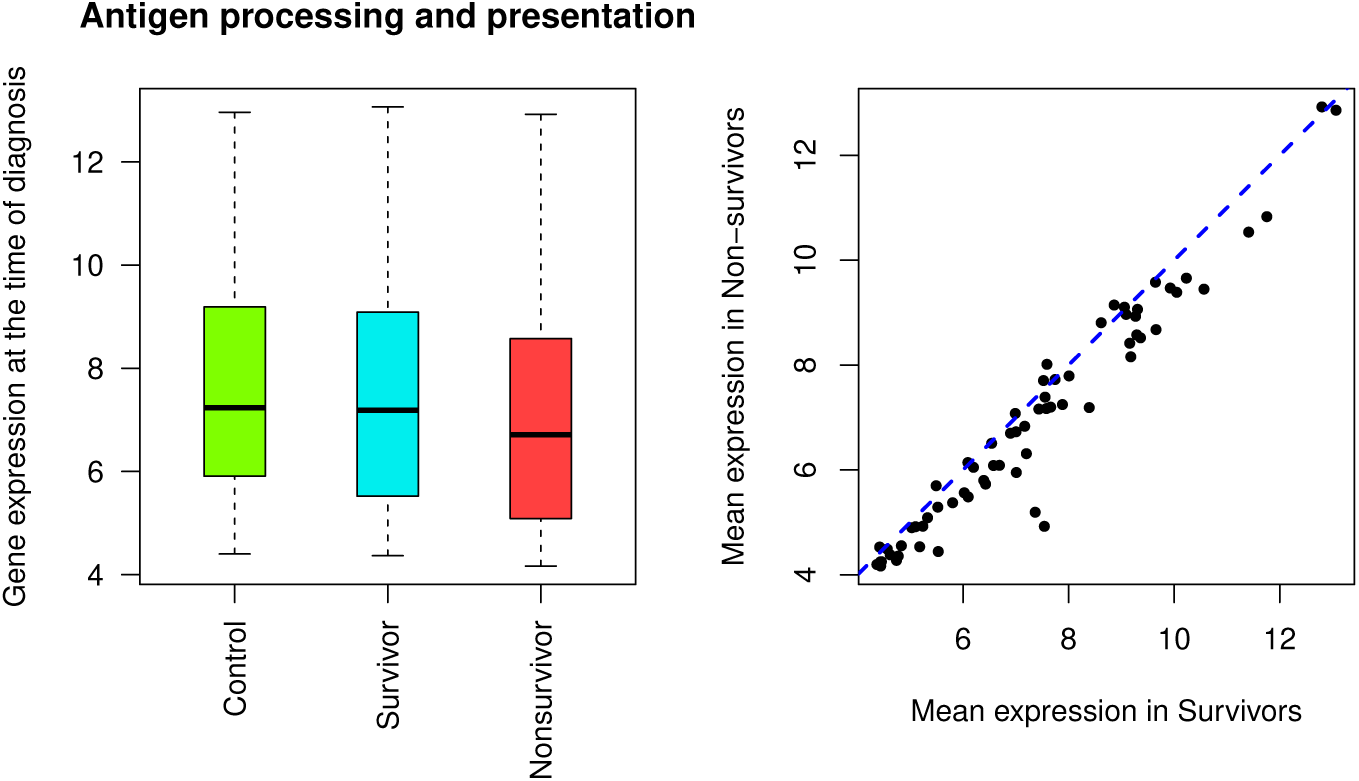
Status of AgPP pathway in NIBMG data with Box/Scatterplot

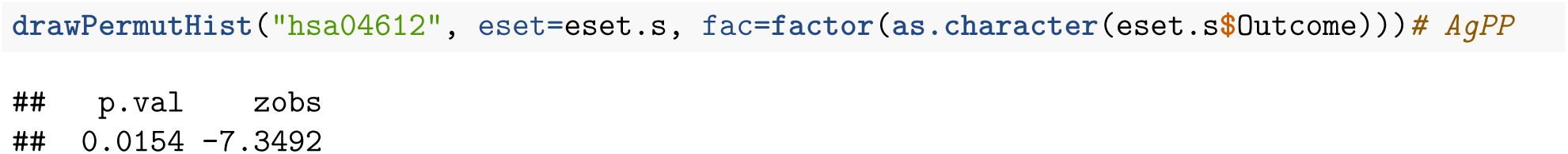

**Figure 13:**
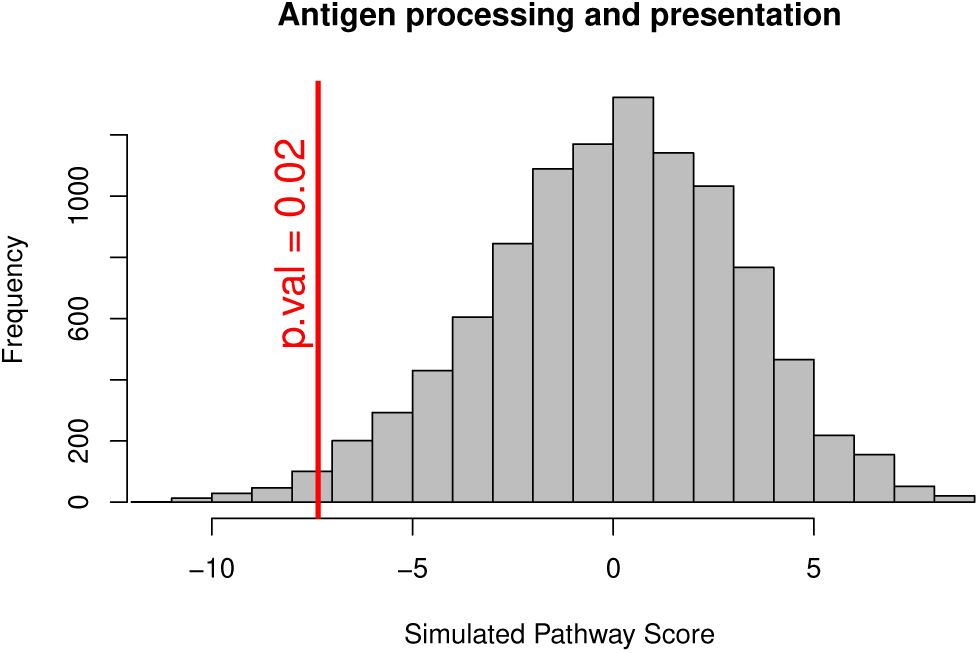
Status of AgPP pathway in NIBMG data, permutation based Gene set enrichment histogram.

### 4.6 Down-regulation of T cell receptor signalling pathway genes in non-survivors

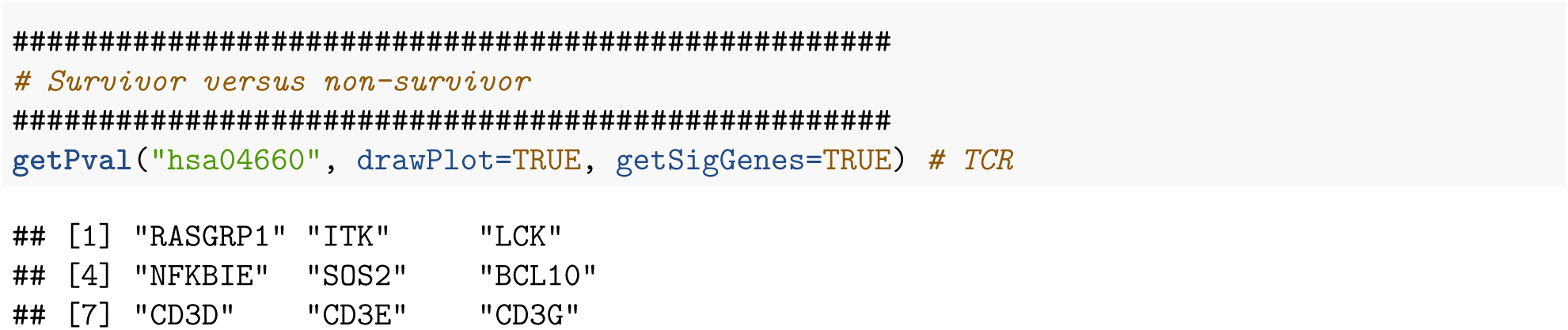

**Figure 14:**
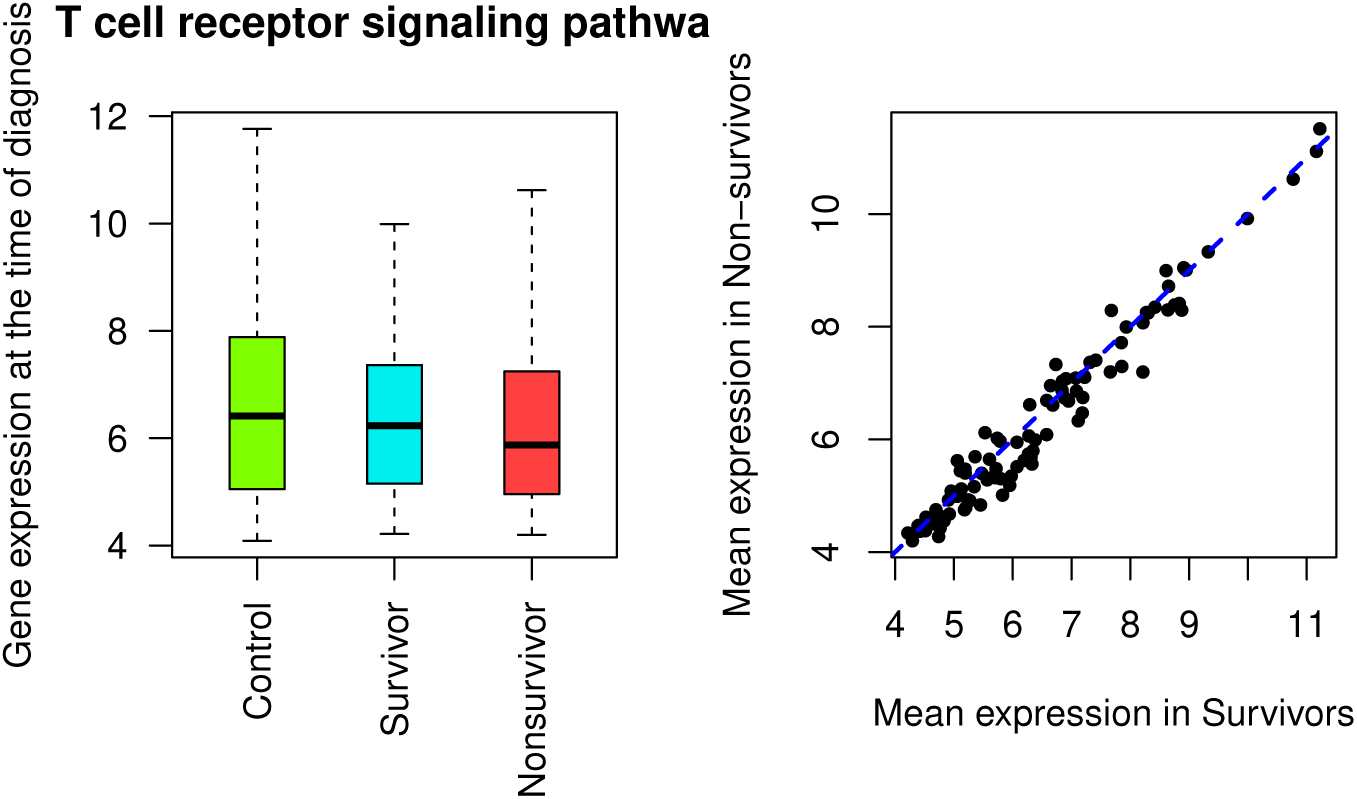
Box plot and scatter plot showing down-regulation of TCR pathway in non-survivors

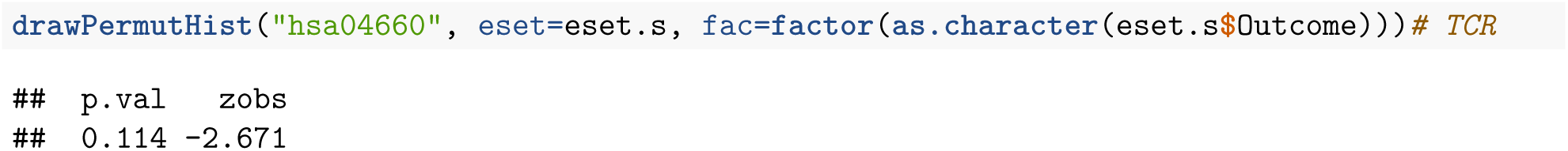

**Figure 15:**
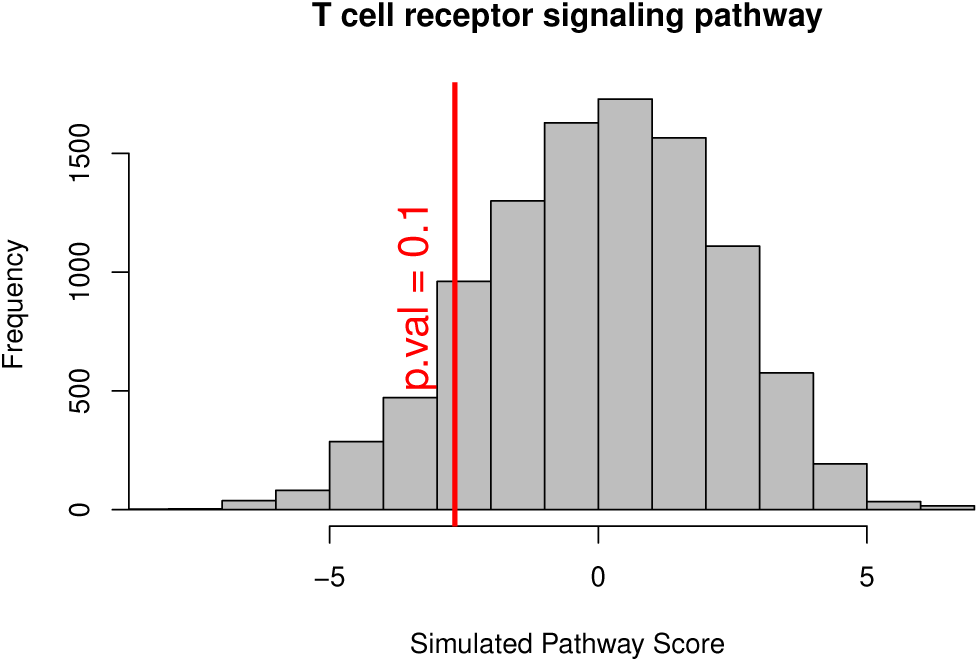
Down-regulation of TCR pathway in SCB cohort, with histogram from permutation based Gene set enrichment test.

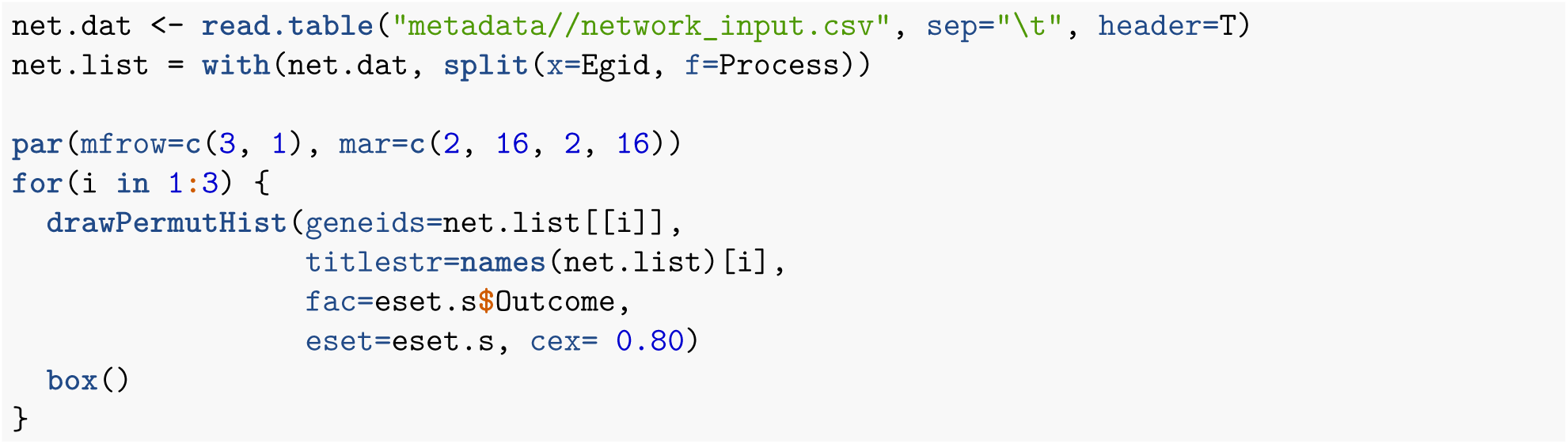

**Figure 16:**
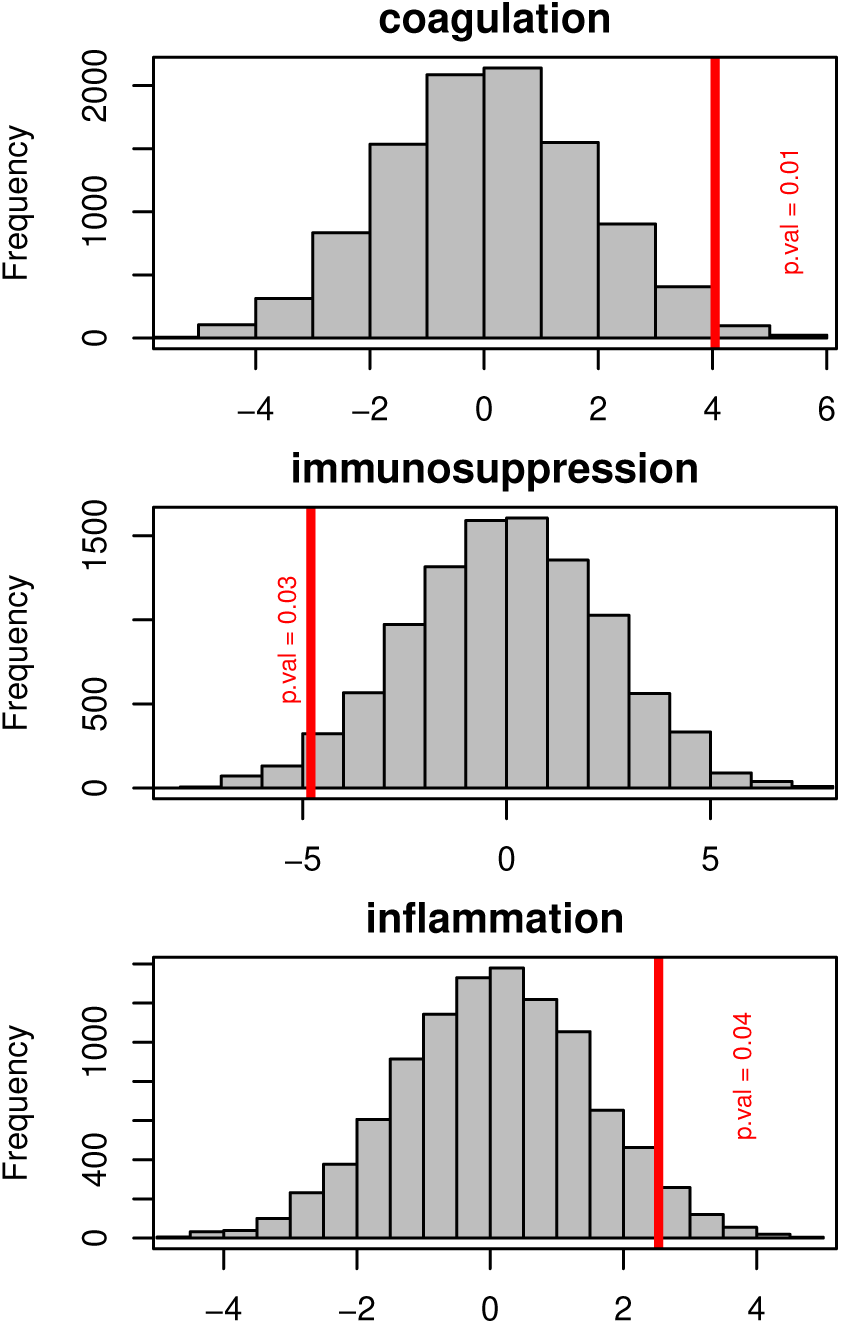
Three Biological processes found to be differentially enriched in Nonsurvivors. Refer to the main manuscript for more details.

## 5 Module scores in SCB cohort

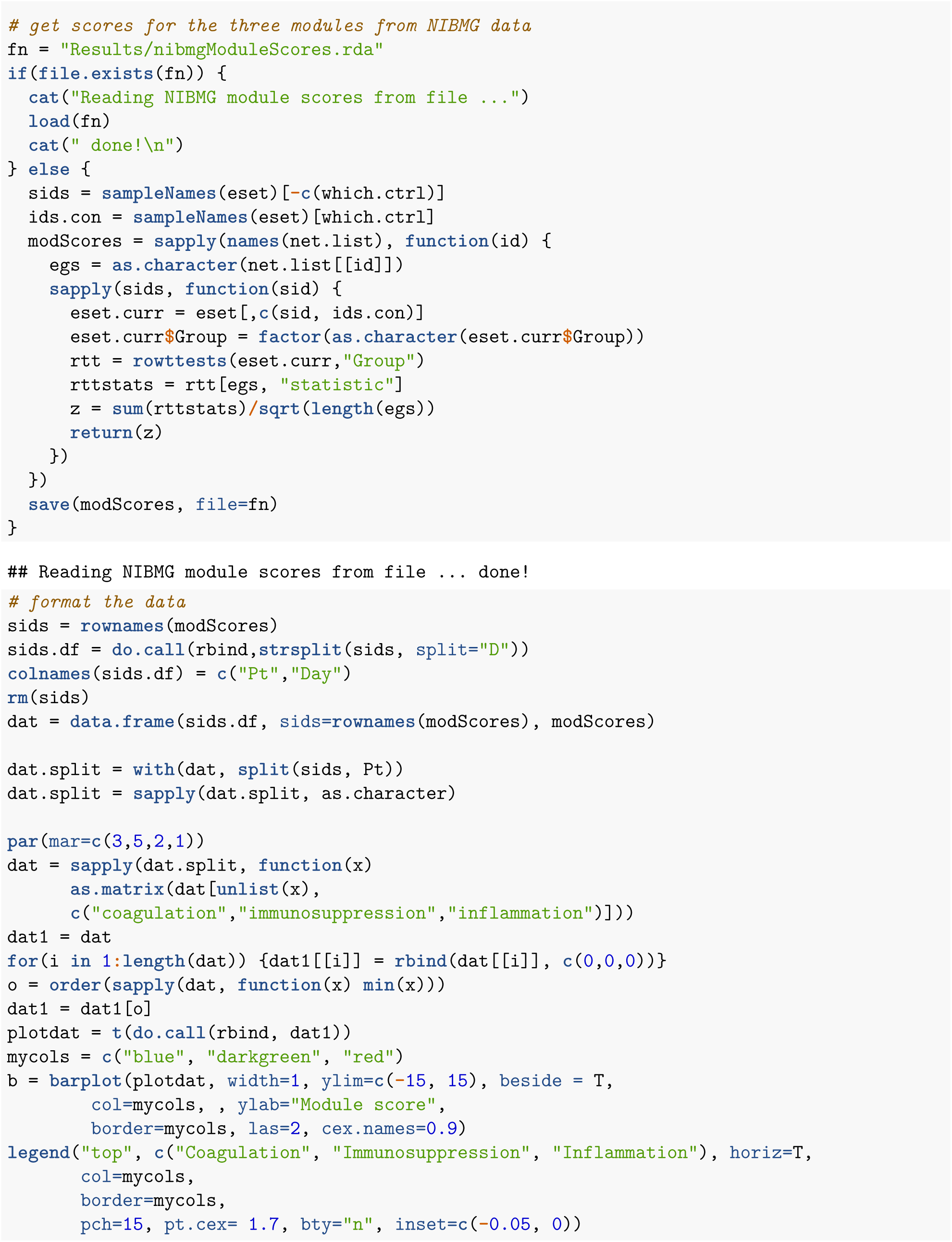

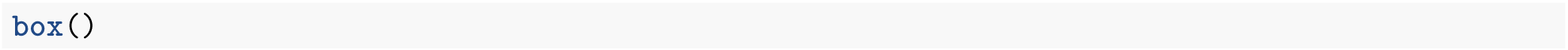

**Figure 17:**
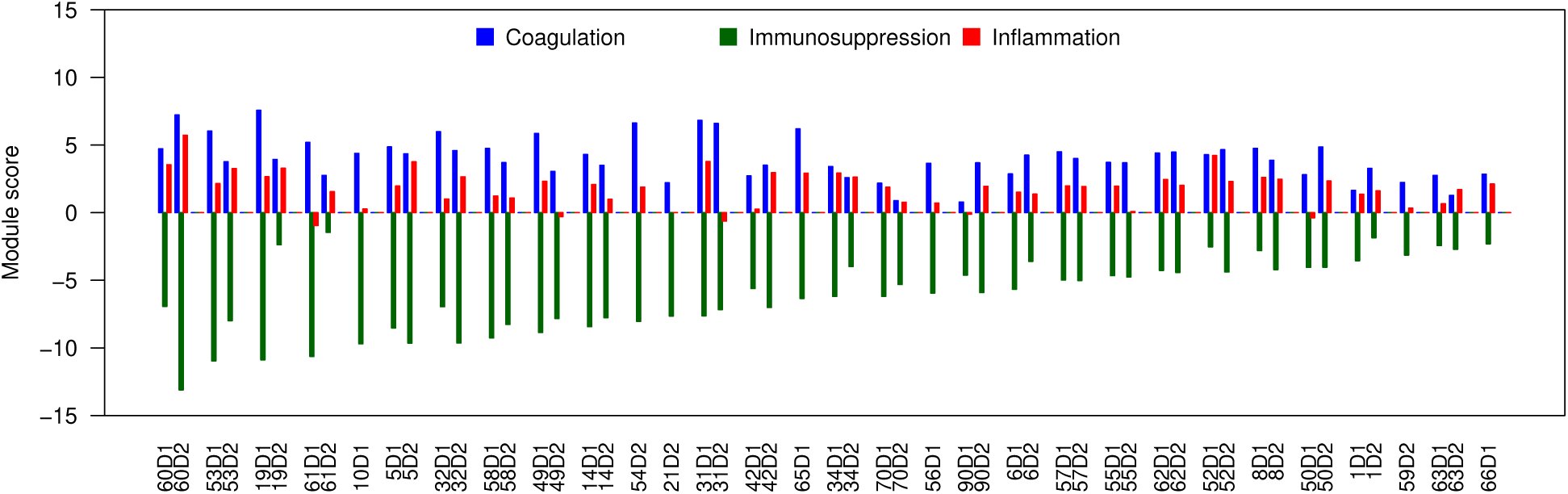
Barplot showing the score of 3 key modules in each sepsis patient

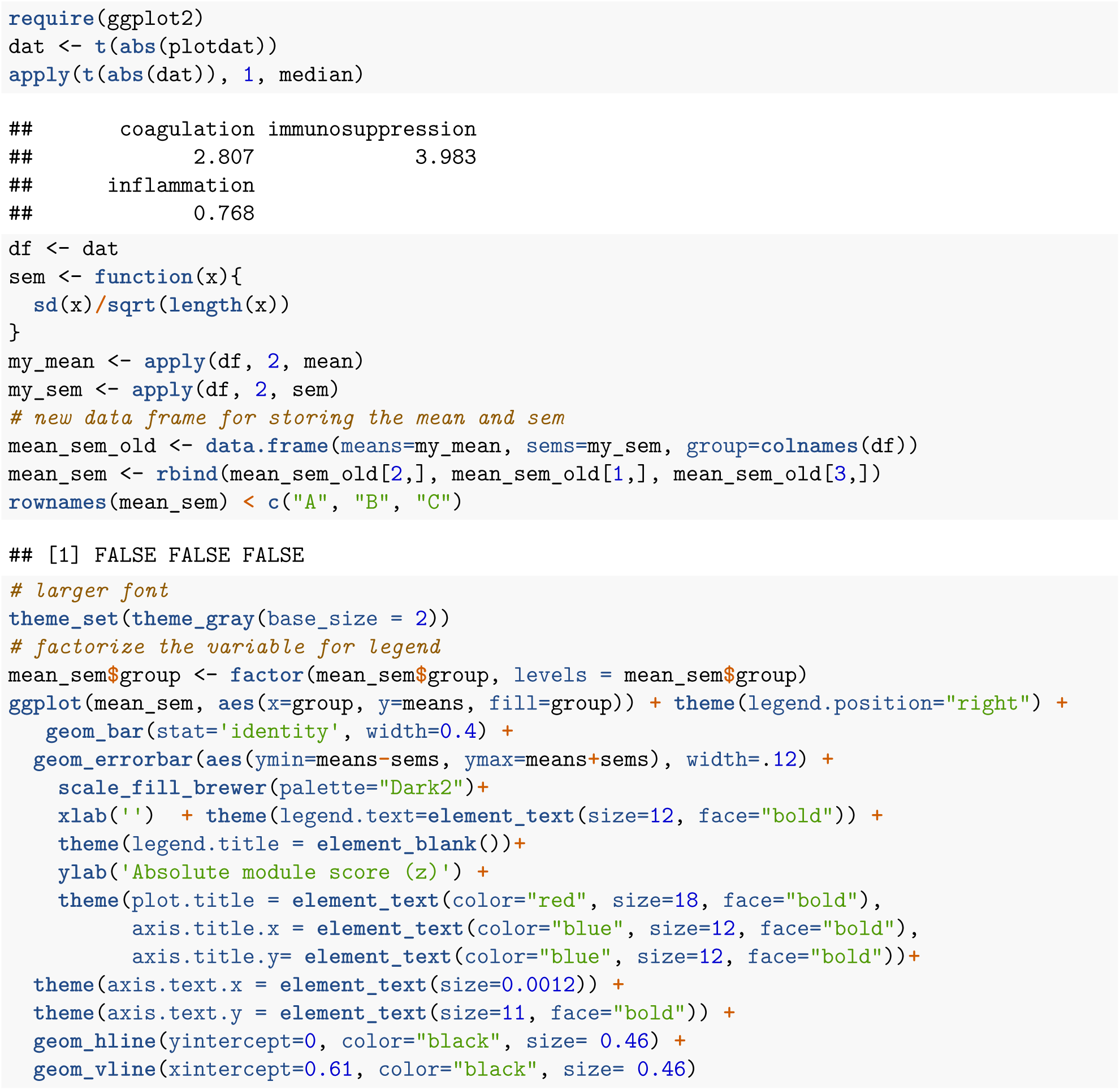

**Figure 18:**
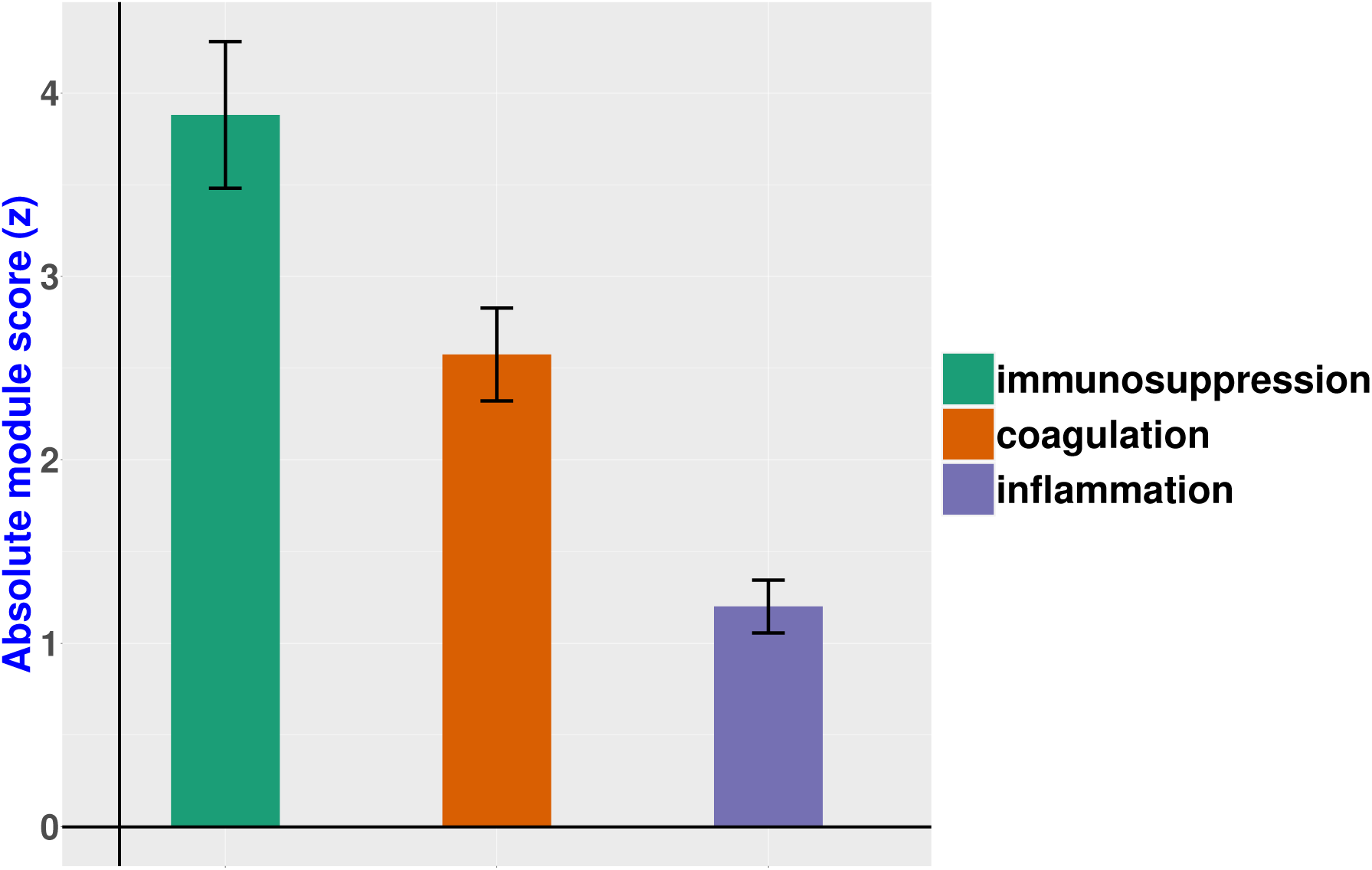
Barplot showing the magnitude of 3 key modules in each sepsis patients. Each bar represents mean of patient-level module score with SEM as the error bar. For all three modules, absolute z-scores have been used. It is clear that there is much greater immunnosuppression compared to inflammtion.

## 6 Session Information

~~~
## R version 3.4.4 (2018-03-15)
## Platform: x86_64-pc-linux-gnu (64-bit)
## Running under: Ubuntu 18.04.1 LTS
##
## Matrix products: default
## BLAS: /usr/lib/x86_64-linux-gnu/openblas/libblas.so.3
## LAPACK: /usr/lib/x86_64-linux-gnu/libopenblasp-r0.2.20.so
##
## locale:
## [1] LC_CTYPE=en_US.UTF-8
## [2] LC_NUMERIC=C
## [3] LC_TIME=en_IN.UTF-8
## [4] LC_COLLATE=en_US.UTF-8
## [5] LC_MONETARY=en_IN.UTF-8
## [6] LC_MESSAGES=en_US.UTF-8
## [7] LC_PAPER=en_IN.UTF-8
## [8] LC_NAME=C
## [9] LC_ADDRESS=C
## [10] LC_TELEPHONE=C
## [11] LC_MEASUREMENT=en_IN.UTF-8
## [12] LC_IDENTIFICATION=C
##
## attached base packages:
## [1] stats4 parallel stats
## [4] graphics grDevices utils
## [7] datasets methods base
##
## other attached packages:
## [1] ssgeosurv_1.0
## [2] ssnibmgsurv_1.0
## [3] ggplot2_3.1.0
## [4] KEGG.db_3.2.3
## [5] pathview_1.18.2
## [6] stringr_1.4.0
## [7] gplots_3.0.1
## [8] pca3d_0.10
## [9] rgl_0.99.16
## [10] KEGGREST_1.18.1
## [11] SPIA_2.30.0
## [12] KEGGgraph_1.38.0
## [13] Category_2.44.0
## [14] Matrix_1.2-15
## [15] GSEABase_1.40.1
## [16] graph_1.56.0
## [17] annotate_1.56.2
## [18] XML_3.98-1.19
## [19] illuminaHumanv2.db_1.26.0
## [20] hgu133plus2.db_3.2.3
## [21] org.Hs.eg.db_3.5.0
## [22] AnnotationDbi_1.40.0
## [23] IRanges_2.12.0
## [24] S4Vectors_0.16.0<colcnt=3>
## [25] genefilter_1.60.0
## [26] limma_3.34.9
## [27] GEOquery_2.46.15
## [28] Biobase_2.38.0
## [29] BiocGenerics_0.24.0
##
## loaded via a namespace (and not attached):<colcnt=3>
## [1] bitops_1.0-6
## [2] bit64_0.9-7
## [3] RColorBrewer_1.1-2
## [4] webshot_0.5.1
## [5] httr_1.4.0
## [6] Rgraphviz_2.22.0
## [7] tools_3.4.4
## [8] R6_2.4.0
## [9] KernSmooth_2.23-15
## [10] lazyeval_0.2.1
## [11] colorspace_1.4-0
## [12] DBI_1.0.0
## [13] manipulateWidget_0.10.0
## [14] withr_2.1.2
## [15] tidyselect_0.2.5
## [16] bit_1.1-14
## [17] compiler_3.4.4
## [18] xml2_1.2.0
## [19] labeling_0.3
## [20] caTools_1.17.1.1
## [21] scales_1.0.0
## [22] readr_1.3.1
## [23] RBGL_1.54.0
## [24] digest_0.6.18
## [25] rmarkdown_1.13
## [26] XVector_0.18.0
## [27] pkgconfig_2.0.2
## [28] htmltools_0.3.6
## [29] htmlwidgets_1.3
## [30] rlang_0.3.1
## [31] RSQLite_2.1.1
## [32] shiny_1.2.0
## [33] bindr_0.1.1
## [34] jsonlite_1.6
## [35] crosstalk_1.0.0
## [36] gtools_3.8.1
## [37] dplyr_0.7.8
## [38] RCurl_1.95-4.12
## [39] magrittr_1.5
## [40] Rcpp_1.0.0
## [41] munsell_0.5.0
## [42] stringi_1.4.3
## [43] yaml_2.2.0
## [44] zlibbioc_1.24.0
## [45] plyr_1.8.4
## [46] grid_3.4.4
## [47] blob_1.1.1
## [48] gdata_2.18.0
## [49] promises_1.0.1
## [50] crayon_1.3.4
## [51] miniUI_0.1.1.1
## [52] lattice_0.20-38
## [53] Biostrings_2.46.0
## [54] splines_3.4.4
## [55] hms_0.4.2
## [56] knitr_1.23
## [57] pillar_1.3.1
## [58] codetools_0.2-16
## [59] glue_1.3.1
## [60] evaluate_0.14
## [61] png_0.1-7
## [62] httpuv_1.4.5.1
## [63] gtable_0.2.0
## [64] purrr_0.2.5
## [65] tidyr_0.8.2
## [66] assertthat_0.2.0
## [67] xfun_0.8
## [68] mime_0.6
## [69] xtable_1.8-3
## [70] later_0.7.5
## [71] survival_2.43-3
## [72] tibble_2.0.1
## [73] memoise_1.1.0
## [74] ellipse_0.4.1
## [75] bindrcpp_0.2.2
~~~

## Notes

https://figshare.com/projects/ssnibmgsurv/67721

